# Single-cell profiling reveals the intratumor heterogeneity and immunosuppressive microenvironment in cervical adenocarcinoma

**DOI:** 10.1101/2024.03.20.586024

**Authors:** Yang Peng, Jing Yang, Jixing Ao, Yilin Li, Jia Shen, Xiang He, Dihong Tang, Chaonan Chu, Congrong Liu, Liang Weng

**Affiliations:** Fourth Department of Gynecologic Oncology, Hunan Cancer Hospital/The Affiliated Cancer Hospital of Xiangya School of Medicine, Central South University, 283 Tongzipo Road, Changsha 410013, Hunan, China; Department of Pathology, Third Hospital, School of Basic Medical Sciences, Peking University Health Science Center, Beijing, 100191, China; Department of Gynecologic Oncology, Changsha Kexin Cancer Hospital, 292 Fenglinsan Road, Changsha 410004, Hunan, China; Department of Pathology, Hunan Cancer Hospital/The Affiliated Cancer Hospital of Xiangya School of Medicine, Central South University, 283 Tongzipo Road, Changsha 410013, Hunan, China; Xiangya Cancer Center, Xiangya Hospital, Central South University, Changsha 410008, China; Key Laboratory of Molecular Radiation Oncology of Hunan Province, Changsha 410008, China; Hunan International Science and Technology Collaboration Base of Precision Medicine for Cancer, Changsha 410008, China

## Abstract

**Background:** Cervical adenocarcinoma (ADC) is more aggressive compared to other types of cervical cancer (CC), such as squamous cell carcinoma (SCC). The tumor immune microenvironment (TIME) and tumor heterogeneity are recognized as pivotal factors in cancer progression and therapy. However, the disparities in TIME and heterogeneity between ADC and SCC are poorly understood.

**Methods:** We performed single-cell RNA sequencing on 11 samples of ADC tumor tissues, with other 4 SCC samples served as controls. The immunochemistry and multiplexed immunofluorescence were conducted to validate our findings.

**Results:** Compared to SCC, ADC exhibited unique enrichments in several sub-clusters of epithelial cells with elevated stemness and hyper-malignant features, including the Epi_10_CYSTM1 cluster. ADC displayed a highly immunosuppressive environment characterized by the enrichment of regulatory T cells (Tregs) and tumor-promoting neutrophils. The Epi_10_CYSTM1 cluster recruits Tregs via ALCAM-CD6 signaling, while Tregs reciprocally induce stemness in the Epi_10_CYSTM1 cluster through TGFβ signaling. Importantly, our study revealed that the Epi_10_CYSTM1 cluster could serve as a valuable predictor of lymph node metastasis for CC patients.

**Conclusions:** This study highlights the significance of ADC-specific cell clusters in establishing a highly immunosuppressive microenvironment, ultimately contributing to the heightened aggressiveness and poorer prognosis of ADC compared to SCC.

## Introduction

Cervical cancer (CC) is the fourth leading cause of cancer-related death among all types of women malignancies (Sung et al., 2021). The two major histological types of CC are squamous cell carcinoma of cervix (SCC) and adenocarcinoma of cervix (ADC), accounting for 70% and 25% of all cases, respectively (Cohen, Jhingran, Oaknin, & Denny, 2019). Compared to SCC, ADC patients have a higher rate of rapid progression, relapse and insensitivity to chemotherapy, radiotherapy and even immunotherapy. Therefore, ADC is considered as a more aggressive phenotype than SCC (Sasieni, Castanon, & Cuzick, 2009). Unfortunately, the current standard treatment for CC is not stratified based on pathologic types. As a result, ADC patients often have suffered from therapeutic failures and experienced short-term recurrence and metastasis, even finishing standard treatment procedures (Sasieni et al., 2009). However, due to the low morbidity of ADC, in-depth investigations are insufficient. Furthermore, the molecular mechanisms elucidating its aggressiveness and the associated biomarkers are not yet well understood. Therefore, this study aims to comprehensively compare the intratumor heterogeneity of ADC, hoping to find out the potential molecule markers specifically associated with ADC.

For early-stage CC patients, it is a crucial and unavoidable issue to have post-surgical upstaging. Over 10% of patients, who are initially diagnosed with early stage and deemed suitable for radical surgery, end up being upstaged due to lymph node metastasis discovered after the surgery (Dabi et al., 2018; Thelissen et al., 2022). This indicates misdiagnosis, leading to different therapeutic strategies and outcome. According to NCCN (National Comprehensive Cancer Network) guidelines, these misdiagnosed patients require additional radiotherapy after surgery, even though they have underwent surgery (Bhatla, Aoki, Sharma, & Sankaranarayanan, 2018). However, combo-treatment of surgery and radiotherapy may cause additional complications and heavier financial burdens, without offering more benefits than radiotherapy alone (Cushman et al., 2018; Kashima et al., 2023). Actually, the optimal therapeutic approach should be radiotherapy alone if the presence of lymph node metastasis could have been accurately diagnosed beforehand (Abu-Rustum et al., 2020). To avoid this problem, the International Federation of Gynecology and Obstetrics (FIGO) in 2018 updated a new stage as IIIC by including lymph node (LN) status and stratified this stage into two sub-groups: IIICp (p stands for pathological findings of LN, which means misdiagnosis) and IIICr (r stands for radiological findings of LN, which means pre-surgical evidence) (Bhatla et al., 2018). Unfortunately, radiological imaging tools do have limitations, since CT or MRI can only detect suspiciously metastatic LN when the minimum diameter is ≥ 10mm (Saleh et al., 2020). Therefore, together with radiological diagnosis, it is urgent to search for potential molecular predictors of LN metastasis, in order to decrease the rate of post-surgical upstaging.

Persistent infection with human papillomavirus (HPV) is the primary cause of CC (Mann, Singh, & Kumar, 2023). However, there are about 5%-15% of patients testing negative for HPV (Yang et al., 2023). It is reported that HPV-negative patients tend to experience worse clinical outcomes (Fernandes, Viveros-Carreño, Hoegl, Ávila, & Pareja, 2022). However, current therapeutic strategies for CC do not take into account the HPV infection status as a factor for stratified therapy. Therefore, it is imperative to investigate the precise mechanisms underlying the development of HPV non-infected tumors.

The tumor immune microenvironment (TIME), consisting of tumor cells, immune cells, cytokines and their interactions, predominantly determines the efficacy of immunotherapy (Lv et al., 2022). In cervical cancer, while it has been reported that 30%-50% of patients exhibit positive PD-L1 expression, however, only an estimated 20%-30% are expected to respond positively to PD-1/PD-L1 blockade therapy (Omenai, Ajani, & Okolo, 2022; Sun et al., 2020). Therefore, cervical cancer, especially the pathological type of ADC, is characterized as an immune-insensitive cancer type, due to atypical modulation of the TIME (Cao et al., 2023). The mechanisms of tumor immune evasion is basically ascribed to dysregulation of TIME and are similar across different types of cancer, involving the deactivation of immune surveillance pathways or recruitment of immunosuppressive cells (Vinay et al., 2015). However, the specific characteristics of the TIME in ADC remain poorly understood. Therefore, in order to optimize precise therapeutic strategies and to enhance the efficacy of immunotherapy, it is indeed imperious to investigate the intricacies of TIME in ADC.

To extensively investigate the cellular and molecular heterogeneity of CC, particularly ADC, the utilization of single-cell RNA sequencing (scRNA-seq) technique is crucially needed. In the current study, we focused on the remodeling of TIME in ADC and performed scRNA-seq analysis on 11 ADC samples, with another 4 SCC samples included as controls. By high-throughput analysis, we are able to characterize the landscape of TIME in ADC, with different types of tumor-promoting cells being recruited in the vicinity of the tumor, thus creating an immunosuppressive environment in ADC. The cellular crosstalk analysis provides valuable insights into the interactions between different cell types within the TIME. Furthermore, we have conducted a screening of novel signature genes as prognostic biomarkers for ADC. These biomarkers have the potential to possibly predict lymph node metastasis and may guide personalized treatment decisions more effectively.

## Methods

### Clinical sample collection

The 15 cases of fresh cervical cancer tissues were collected from Hunan Cancer Hospital, with informed consent obtained from 15 independent patients. The inclusion criteria are as follows: 1) The diagnosis of CC, with pathological type of ADC or SCC, should be confirmed by pathologists; 2) Pre-surgical FIGO stages should be within the range from IB (which means the tumor mass should be detectable via CT or MRI) to IIA (which means no parametrial involvement, nor distant metastasis), in which patients have explicit indication for radical surgery following NCCN guidelines; 3) No neo-adjuvant treatments should be given before the surgery, including chemotherapy and radiotherapy; 4) The surgery options should be type-C radical hysterectomy and include pelvic lymphadenectomy with/without para-aortic lymphadenectomy. Detailed information of the 15 patients was listed in **Supplementary Table 1**. This study was supervised and approved by the Ethics Committee Board of Hunan Cancer Hospital.

### Sample preparation for single-cell isolation

Immediately after surgical removal, the fresh tumor tissues were washed by saline for 3 times and stored in cold storage solution (MACS® Tissue Storage Solution) for transportation. To prepare high-quality samples, the tissues were cut into pieces to 1mm^3^ and were incubated at 37°C for 30 minutes. 0.25% trypsin solution was used to digest the tissue pieces at 37°C for 10 minutes. Then single-cell samples were filtered through a 70-μm meshed filter and suspended in a red blood cell lysis buffer (MACS® Red Blood Cell Lysis Solution) for dissolution. Before instrumental sequencing, suspension of single cells were visualizing by TC20^TM^ Automated Cell Counter to assess the cell viability. Each sample was confirmed to have a cell viability rate of over 80%.

### Single-cell RNA sequencing

The suspension of single cells was transferred onto the 10X Chromium Single-Cell instrument to generate single-cell beads in the emulsion (GEMs). Then the scRNA-seq libraries were constructed by using the Chromium Controller and Chromium Single Cell 3’Reagent Kits (v3 chemistry CG000183) and sequenced by using the sequencer Novaseq6000 (Illumina, USA). All procedures were following the manufacturer’s protocol.

### Data processing, dimension reduction and cell clustering

The raw scRNA-seq reads were processed for barcode processing, genome mapping and the gene expression matrixes were generated by using the Cell Ranger toolkit (version 5.0.0) of 10X Genomics platform. The GRCh38 human reference genome was utilized in the read alignment process. The unique molecular identifiers (UMIs) were counted in each cell. As for data processing, cells with low qualities, which showed <200 expressed genes or >25% mitochondrial UMIs, were excluded. The package of Seurat R (version 4.0.5 R) was applied for quality control. The Scrublet software (version 0.2.2) was utilized to identify and remove potential doublets. As for data normalization and conversion, the LogNormalize method was implemented in the NormalizeData function, and then the normalized counts were log-transformed. With the function of RunPCA, the principal component analysis (PCA) was used for data dimension reduction. The functions of FindNeighbors and FindClusters were used for cell clustering. For visualization, the RunUMAP function was performed. In each unsupervised cell cluster, the gene markers were identified by the function of FindAllMarkers with comparison to other clusters.

### Cell type annotation

The cell typing was conducted and annotated according to the selected gene markers via CellMarker database and publications (C. Li et al., 2022; Qiu et al., 2023; Xue et al., 2022). The expression of differential expressed genes (DEGs) was used to determine each cell type following the rules: Log_2_Foldchange should be > 0.2 and adjusted-*P* values should be < 0.05.

### Pseudotime trajectory analysis

To identify the developmental changes of epithelial cells, the pseudotime trajectory analyses were performed by using Monocle2 R package (version 2.22.0), in which Seurat was used as input. Calculated by the Monocle algorithm, the top 50-1000 genes were conducted for differential GeneTest. Then, the DDR-Tree and default parameters were generated to visualize the translational relationship and developmental orders among different epithelial sub-clusters.

### Cell malignancy scoring

Among these epithelial cells, the malignancy score of each single cell was predicted by using scCancer R package (Guo et al., 2021). The malignant cells were distinguished from non-malignant cells by inferring large-scale copy-number variations (CNVs), which was calculated by the interCNV R package as described (Guo et al., 2021; Xue et al., 2022).

### Signaling pathway enrichment analysis

The biological signaling pathway of each cell cluster was identified by performing Gene Ontology (GO) analyses and Hallmark Pathway enrichment analyses, according to the Molecular Signature Database (MSigDB version 7.5.1). Then DEGs of each cell cluster were converted to ClusterProfiler (version 4.2.2) package for analyses of predicted functions.

### CytoTRACE

In order to predict the differential state of cells, scRNA-seq derived data were pass to the CytoTRACE software (version 0.3.3) with R package for further analyses.

### Survival analysis

The CESC dataset from TCGA database was downloaded from Xena, with clinical characteristics, survival endpoints and treatments were included. The Kaplan-Meier survival curves were generated by using R software.

### Cellular communication analysis

The possible interactions, with ligand-to-receptor communicating signaling pathway, between two types of cells were analyzed using the R packages of CellChat (version 1.5.0). Seurat was used for normalization and the rank with significance was calculated. The probability of ligand-to-receptor interaction in each signaling pathway was plotted by *P*-value and intensity.

### Immunohistochemistry (IHC)

The paraffin-embedded ADC samples (whether post-surgical or biopsy samples) were sectioned into 4μm slides. The protocol of IHC strictly followed our previous published works (Peng et al., 2019). The primary antibody of SLC26A3 was purchased from Santa Cruz Biotechnology (Cat# sc-376187) and diluted at the ratio of 1:20. The protein expression of SLC26A3 was determined using the method of *H*-scoring: *H*-score = intensity score × percentage of positivity. The intensity was scored as: Negative=0, Weak=1, Moderate=2, Strong=3. Samples were grouped as the higher expression group (H-score ≥ 150) and the lower expression (H-score < 150) group of SLC26A3.

### Multiplexed immunofluorescence (multi-IF)

The multi-IF was performed using the Opal 6-Plex Manual Detection Kit (NEL811001KT, Akoya), following the manufacturer’s instructions. Generally, the slides were stained with each primary antibody sequentially following the staining steps in each cycle as: (1) antigen retrieval by citrate (pH=6) at water bath for 30 minutes; (2) independent primary antibody incubation at 4°C atmosphere overnight or 37°C atmosphere for 1 hour; (3) common HRP-crosslinked secondary antibody incubation at 37°C atmosphere for 30 minutes; (4) independent opal reactive fluorophore solution at intended wavelength (opal 570 was excluded in order to save for FOXP3 signal). Each independent primary antibody started a new cycle. After all cycles finished, the slides were prepared for incubation of FOXP3 monoclonal antibody with eFluor™ 570 (Thermo Fisher, Cat#41-4777-82) conjugated, as well as DAPI (Sigma) for nucleus staining.

### Statistics

Specific analyses of scRNA-seq data were decoded and visualized by using the R software. Statistics were processed using IBM-SPSS Statistics 20.0 software and R software as well, including Student’s *t*-test, two-sided Wilcoxon test, etc. Correlations analysis was performed by the *Chi*-square test. Survival analysis was performed by the Kaplan-Meier method. *P* values < 0.05 were considered as statistically significant.

## Results

### Single-cell transcriptomic landscape of cervical cancer

We collected 15 samples of cervical cancer tissues from 15 individual patients immediately after undergoing radical hysterectomy surgery. Among these cases, 11 were ADC types, with 5 being HPV-negative and 6 being HPV-positive. The remaining 4 cases were SCC types, with an even distribution of HPV-positive and HPV-negative status (**Supplementary Table 1**). Prior to surgery, all 15 patients were diagnosed as early FIGO stages (stage I ∼IIA) based on evaluation of CT, MRI, and physical examination. However, after surgery, 5 cases turned out to be upstaged with pathological evidence of lymph node metastasis (i.e. FIGO stage IIICP), requiring theses patients to receive radiation therapy according to NCCN guidelines. Therefore, in addition to investigate the difference between histological types and HPV infection status, we also focused on the issue of post-surgical upstaging by comparing different FIGO stages.

In order to analyze the genomic profiling, we resolved these samples at the single-cell level using the 10X genomics chromium platform. The cell viability and sample quality were ensured beforehand, and a total of 102,012 qualified cells were utilized for further analysis (**Fig. 1A**). The technique of Uniform Manifold Approximation and Projection (UMAP) was utilized to reduce the dimension of gene profiling data for visualization. The obtained cells were primarily categorized into 12 cell clusters, which were further re-clustered into 9 distinct cell types, based on specific gene markers (C. Li et al., 2022) (**Fig. 1B-C**). These cell types included T cells (38,627 cells in total, proportioning 37.87%, marked with PTPRC, CD3D, CD3E and CD3G), epithelial cells (32,199 cells in total, proportioning 31.56%, marked with EPCAM, SLP1 and CD24), neutrophils (12,586 cells in total, proportioning 12.34%, marked with CSF3R), macrophages (6,798 cells in total, proportioning 12.34%, marked with CD68 and CD163), fibroblasts (4,258 cells in total, proportioning 4.17%, marked with COL1A2 and DCN), plasma cells (4,044 cells in total, proportioning 3.96%, marked with JCHAIN), endothelial cells (1,389 cells in total, proportioning 1.36%, marked with ENG and VWF), B cells (1,326 cells in total, proportioning 1.30%, marked with CD79A and MS4A1) and mast cells (785 cells in total, proportioning 0.77%, marked with MS4A2) (**Fig. 1D-E** and **Supplementary Table 2-3**).

**Figure 1.**
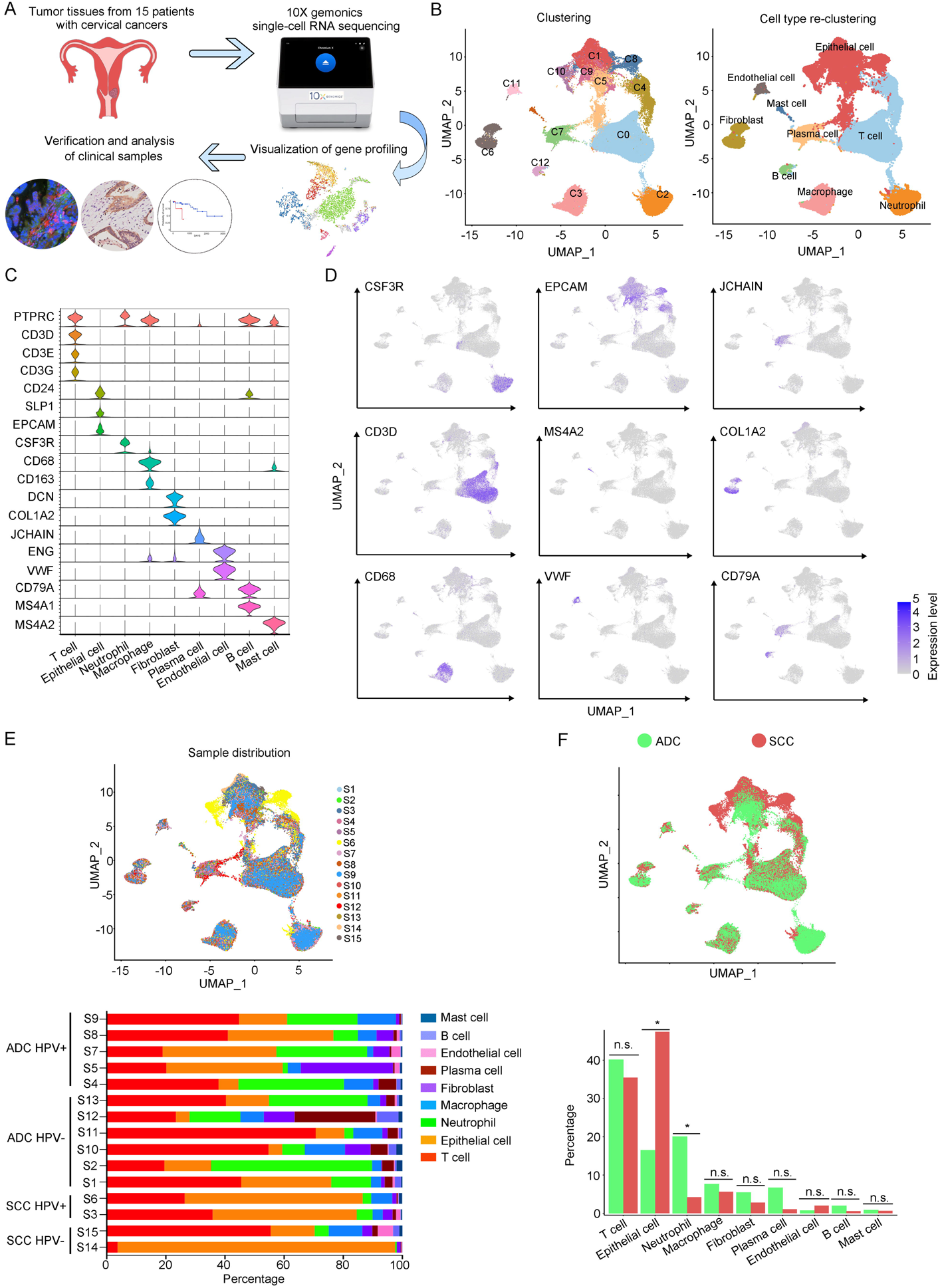
The single-cell genomic atlas of cervical cancer. **A.** The schematic design of sample collection, single-cell RNA sequencing, data processing and clinical validating through 10X genomics platform (The sketch of scRNA-sq machine, as well as the plot of dimension-reduction visualization, was originally cited from the official website of 10X genonics: https://www.10xgenomics.com). **B.** UMAP plots of all single cells by original clustering and cell type re-clustering, according to the published lineage-specific marker genes, respectively. **C.** Violin plots demonstrating the expression of marker genes that correspond to each of the major cell types. **D.** UMAP plots demonstrating the most typical marker gene specific to each type of cell cluster. **E.** UMAP plot and histogram plot showing the occupation ratios of the major cell types in each individual sample, with HPV status and histological type summarized. **F.** UMAP (up) and histogram (down) plots to show the differences of distribution and proportion of each cell type between different histological types (ADC vs. SCC). Statistics were performed using R software with two-sided Wilcoxon test (*P* values for each group are listed below: T cell: *P=*0.661; Epithelial cell: *P*=0.039; Neutrophil: *P*=0.026; Macrophage: *P*=0.661; Fibroblast: *P*=0.661; Plasma: *P*=0.078; Endothelial cell: *P*=0.851; B cell: *P*=0.104; Mast cell: *P*=0.753). Statistics were shown as \**P* < 0.05; \*\**P* < 0.01; n.s. not significant.

According to our scRNA-seq data, ADC exhibited a significant higher proportion of neutrophils than SCC (20.05% vs. 4.23%). The proportion of T cells was also higher in ADC (41.13%) compared to SCC (35.48%). Conversely, SCC showed a nearly threefold enrichment of epithelial cell (47.43%) compared to ADC (16.48%) (**Fig. 1F**). When considering the HPV infection status, HPV-negative patients demonstrated decreased proportions of epithelial cells (7.78% vs. 28.38%) and neutrophils (16.66% vs. 24.68%) compared to HPV-positive patients. The proportion of T cells enriched in HPV-negative cases (45.77%) was higher than that in HPV-positive cases (32.42%). HPV-negative patients exhibited prominently higher proportions of plasma cells (10.13% vs. 1.99%) and B cells (2.84% vs. 0.77%) compared to HPV-positive patients, even though their proportions were lower than T cells, epithelial cells and neutrophils (**Supplementary Fig. 1A**). Furthermore, we conducted a comparison between early-stage and late-stage cases, revealing that late-stage patients have a higher proportion of epithelial cells (35.50%) than early-stage cases (28.82%) (**Supplementary Fig. 1B**). In summary, these data describes primarily a complex microenvironment in cervical cancer, especially in the ADC type, with varied composition of immune and tumor cells under different conditions.

### The sub-clusters of epithelial cells in ADC exhibit elevated stem-like features

The aforehand data has shown that the majority of the CC tumor is composed of epithelial cells. In order to explore the characteristics of these epithelial cells within the TIME of ADC, we further classified them into 12 sub-clusters based on the subsets of differently expressed genes (DEGs) in each group (**Fig. 2A**). Each sub-cluster was named based on the most prominently up-regulated gene in each panel of signature genes (**Fig. 2B-C**). Several epithelial cell clusters were strongly enriched in ADC than SCC, such as Epi_04_TFF2, Epi_06_TMC5, Epi_07_CAPS and Epi_08_SCGB3A1. Notably, Epi_09_SST, Epi_10_CYSTM1, Epi_11_REG1A and Epi_12_RRAD seemed to be exclusively enriched in ADC, not in SCC (**Fig. 2D**). The stratified enrichment of different cluster between ADC and SCC has captured our attention for further investigation. Additionally, Epi_09_SST, Epi_10_CYSTM1 and Epi_11_REG1A were solely enriched in HPV-positive cases, although no distinctive clusters had been found in HPV-negative patients (**Supplementary Fig. 2A**). When comparing samples from patients in the early stages with those who have lymph node metastasis, 3 clusters (Epi_02_IGLC2, Epi_05_CCL5 and Epi_12_RRAD) were slightly increased among the late-stage patients. Surprisingly, we found that the sub-cluster Epi_10_CYSTM1 was exclusively present in stage IIIC patients, indicating that it might be a potential target for identification of the biomarkers for late stage patients (**Supplementary Fig. 2B**).

**Figure 2.**
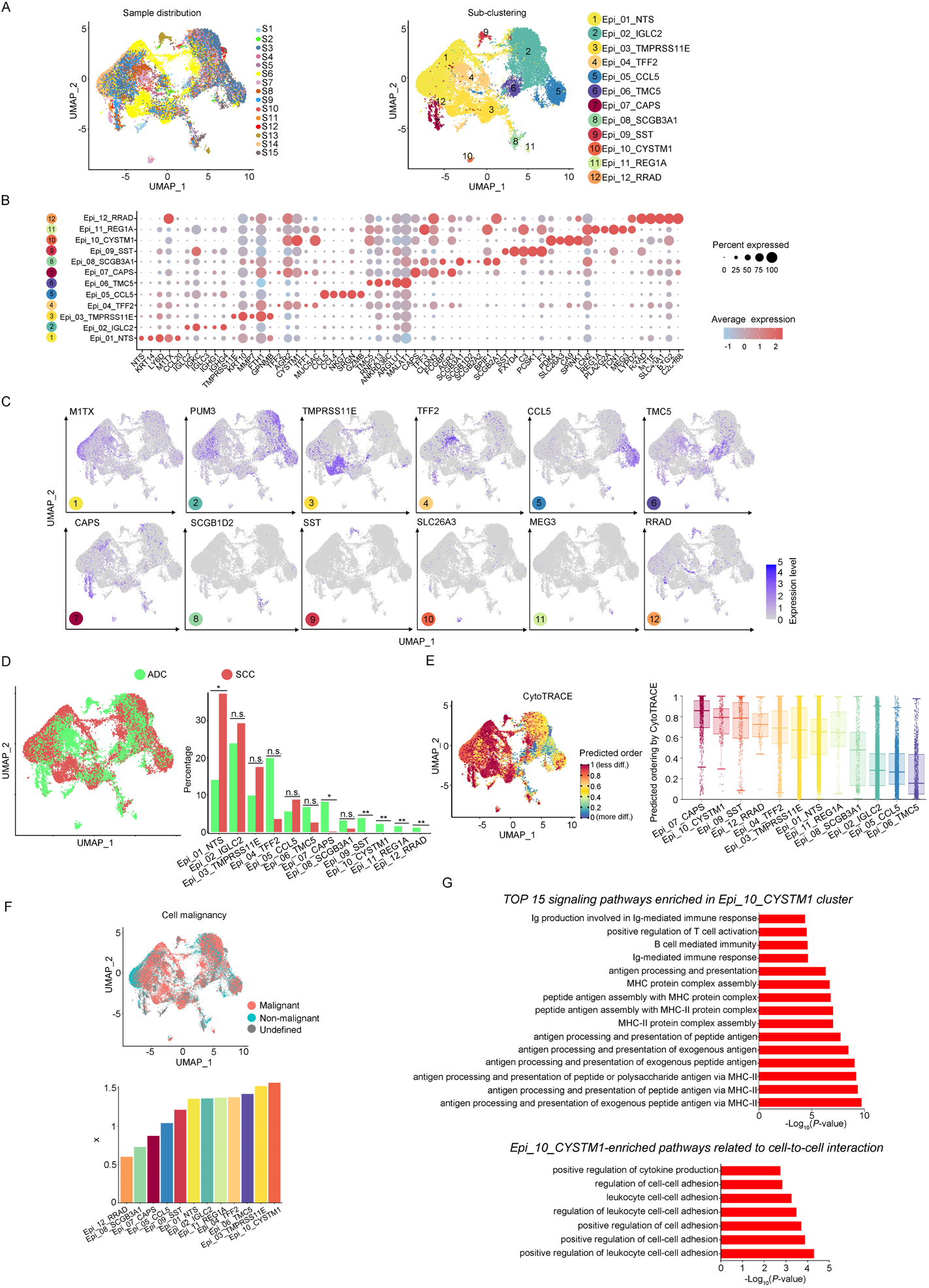
The scRNA-seq data reveals the malignant features of tumor epithelial cells. **A.** UMAP plots showing the distribution of epithelial cells in 15 samples and the sub-clustering into 12 clusters according to ectopic gene expressions. Each cluster of epithelial cells was named using the most highly enriched gene. **B.** Heatmap plot showing the annotation of each epithelial sub-cluster with top 5 DEGs. **C.** UMAP plots demonstrating the most specific gene as the marker for each sub-cluster. **D.** UMAP (left) and histogram (right) plots to compare the differences of distribution and proportion of each epithelial cell sub-cluster between different histological types (ADC vs. SCC). Statistics were performed using R software with two-sided Wilcoxon test (*P* values for each group are listed below: Epi_01_NTS: *P=*0.040; Epi_02_IGLC2: *P*=0.661; Epi_03_TMRPSS11E: *P*=0.661; Epi_04_TTF2: *P*=0.104; Epi_05_CCL5: *P*=0.950; Epi_06_TMC5: *P*=0.412; Epi_07_CAPS: *P*=0.040; Epi_08_SCGB3A1: *P*=0.412; Epi_09_SST: *P*<0.001; Epi_10_CYSTM1: *P*<0.001; Epi_11_REG1A: *P*<0.001; Epi_12_RRAD: *P*<0.001). Statistics were shown as \**P* < 0.05; \*\**P* < 0.01; n.s. not significant. **E.** UMAP plot of CytoTRACE showing the thermal imaging projection of predicted developmental order (left) and histogram plot showing the ranking of CytoTRACE scores (right). **F.** UMAP plot of malignancy analysis demonstrating the cell malignancy features (up) and ranking of scores (down). **G.** GOBP analyses showing the top 15 enriched signaling pathways of Epi_10_CYSTM1 that are more active in ADC than SCC (up), particularly pathways related to cell-to-cell interactions (down). The histogram data are transformed from -Log_10_(*P*-value) for visualization. ***Abbreviations:*** Ig: immunoglubolin; MHC: major histocompatibility complex.

We further evaluated the features of each epithelial cluster, especially their predicted roles in regulating tumorigenesis. To assess the differentiation potential of each cell cluster, we employed the Cellular Trajectory Reconstruction Analysis using gene Counts and Expression (CytoTRACE) method (**Fig. 2E**). The results showed that clusters of Epi_07_CAPS, Epi_09_SST, Epi_10_CYSTM1 and Epi_12_RRAD ranked the top four in CytoTRACE score, indicating the potential for pluripotency and differentiation, also referred to stem-like features. Interestingly, Epi_09_SST, Epi_10_CYSTM1 and Epi_12_RRAD were exclusively enriched in ADC. The proportion of cluster Epi_07_CAPS, which acquired the highest level of stemness, was higher in ADC, although Epi_07_CAPS was also identified in SCC (**Fig. 2D**). Subsequently, pseudotime analysis was conducted to elucidate the differentiation trajectories. The results showed that Epi_07_CAPS, Epi_09_SST and Epi_10_CYSTM1 were positioned towards the end of pseudotime developmental trajectory, which was in consistence with their high stem-like characteristics and indicated a higher degree of malignancy in ADC than SCC (**Supplementary Fig. 2C**). In addition, we performed the malignancy scoring analysis, which demonstrated that cluster Epi_10_CYSTM1 was predicted to exhibit the highest degree of malignancy compared to other clusters (**Fig. 2F**). Interestingly, cluster Epi_10_CYSTM1 displayed a diverse developmental profile and was exclusively identified in ADC. Conversely, Epi_07_CAPS and Epi_12_RRAD, which exhibited high stem-like characteristics, were ranking lower in terms of malignancy scoring when compared to other clusters. The observation of cluster Epi_10_CYSTM1 and its potential specificity to ADC raises the question of whether this cluster is associated with the aggressiveness of ADC.

Besides, we performed Gene Ontology Biological Process (GOBP) analysis on specific signaling pathways based on the gene expression pattern of cluster Epi_10_CYSTM1 (**Fig. 2G**). The results revealed that cluster Epi_10_CYSTM1 tended to be more active in regulating immunity-related pathways, which indicated this cluster might be possibly associated with dysfunction of immunity. In addition, cluster Epi_10_CYSTM1 was predicted to be more active in regulating cell-to-cell interaction (**Fig. 2G**). Therefore, further investigation is warranted to explore the intricate crosstalk networks between epithelial cells and other cell types in the TIME of ADC.

### ADC-specific epithelial cluster-derived gene SLC26A3 is a potential prognostic marker for lymph node metastasis

Based on the aggressive characteristics of cluster Epi_10_CYSTM1 from above bioinformatic predictions, we further examined the correlation between this cluster and the clinical features of CC. For validation, we alternatively examined cluster Epi_12_RRAD (**Supplementary Fig. 3A**), which was another malignant cluster predominantly found in ADC (**Figure. 2D**). We firstly checked the DEGs of these two clusters. In Epi_12_RRAD cluster, we identified Insulin-like growth factor 2 (IGF2) and Alcohol dehydrogenase 1C (ADH1C), which were exclusively expressed in Epi_12_RRAD, as two most specific genes of this cluster (**Supplementary Fig. 3B**). However, the immunohistochemistry (IHC) staining showed that IGF2 was almost negative across cases with different clinical stages (**Supplementary Fig. 3C**). On the other hand, ADH1C was strongly expressed in both early-stage and late-stage cases (**Supplementary Fig. 3C**). Likewise, the survival analysis of Epi_12_RRAD-enriched gene signatures did not demonstrate a significant difference between the two groups (**Supplementary Fig. 3D**). These results suggest that cluster Epi_12_RRAD may not be a potential target when relating to the clinical features of ADC.

In the cluster of Epi_10_CYSTM1, the DEG with the highest expression is CYSTM1 (**Fig. 3A**). However, CYSTM1 was also expressed in other cell clusters and showed low specificity (**Fig. 3B**). On the other hand, SLC26A3, ORM1 and ORM2 were specifically enriched in Epi_10_CYSTM1 and could be considered as potential candidate biomarkers for representation of this cluster. Nevertheless, ORM1/OMR2 were positively expressed but were not satisfied to distinguish the severity of CC cases, due to an irrelevance between expression intensity and clinical stages (**Supplementary Fig. 3E**). Interestingly, SLC26A3, as a representative marker of cluster Epi_10_CYSTM1, showed potential value to associated with late clinical stages for CC patients (**Supplementary Fig. 3E**). This is encouraging, since currently it lacks biomarkers to predict late stages of cervical cancer and the only way to detect LN metastasis beforehand is using radiological tools, which have technical limitations.

**Figure 3.**
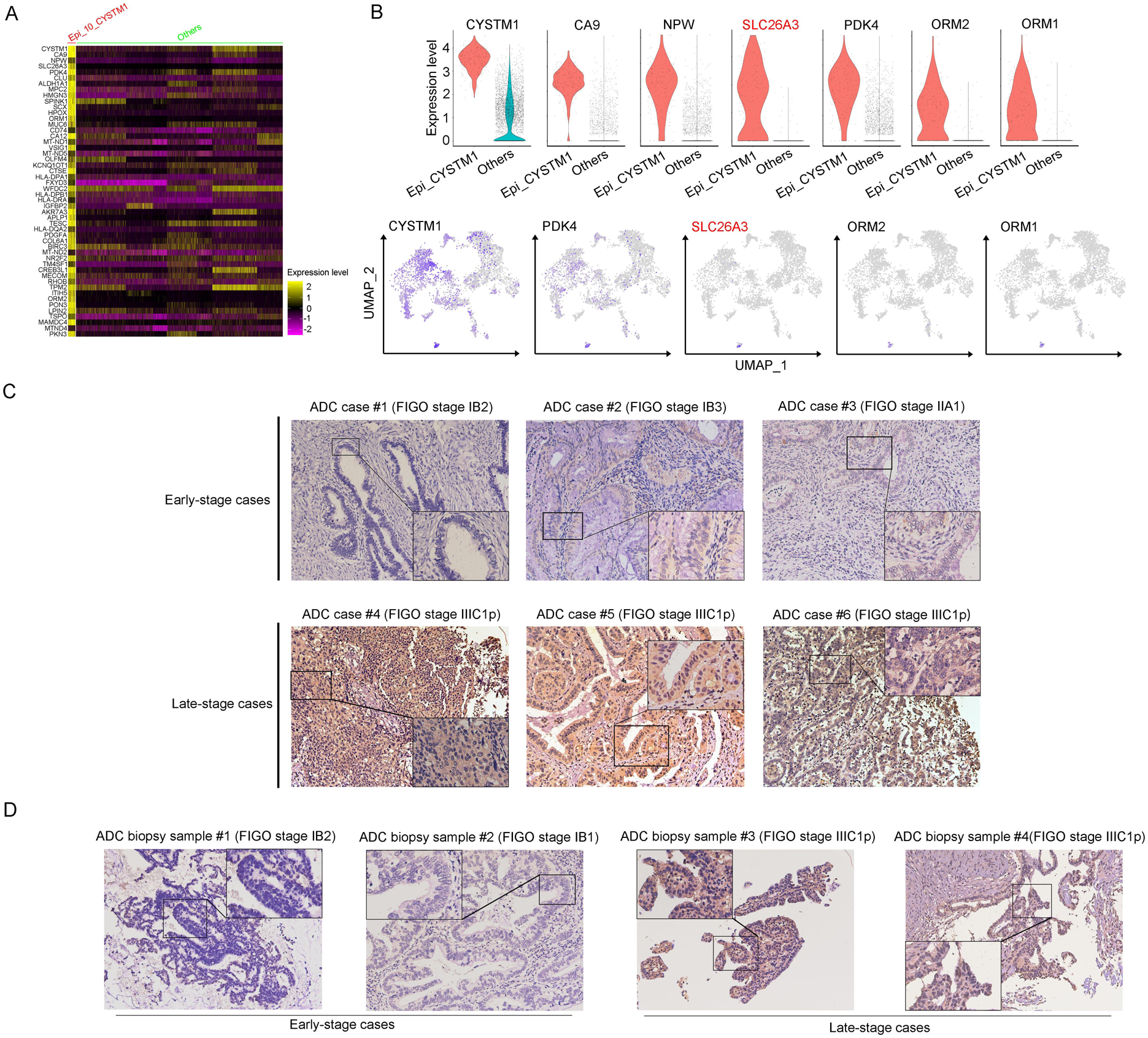
SLC26A3 is identified as a potential prognostic and diagnostic indicator for lymph node metastasis of CC patients. **A.** Heatmap showing the top 50 DEGs of Epi_10_CYSTM1 cluster in comparison to all the other sub-clusters (Red: Epi_10_CYSTM1 cluster; Green: other epithelial cell sub-clusters). **B.** Violin plots showing the expression difference between two groups (top), with UMAP plots (bottom) demonstrating the specificity of each candidate gene, filtering SLC26A3 as the most identical marker for this sub-cluster. **C.** IHC staining showing the protein expression of SLC26A3 in surgically-resected ADC samples which are classified as early stages (FIGO stage I-IIA) and late stages (FIGO stage IIIC1∼2p). Images from 6 individual cases were shown as representatives for each group. **D.** IHC staining showing the protein expression of SLC26A3 in biopsy ADC samples which are classified as early stages (FIGO stage I-IIA) and late stages (FIGO stage IIIC1∼2p). Images from 4 individual cases were shown as representatives for each group. The method of *H*-score is used and the scoring system is as follows: negative (0), weak (1), intermediate (2), and strong (3). Expression is quantified by the H-score method.

This is a problem because misdiagnosis of clinical staging affects the disease outcome and treatment approach for CC patients. In order to examine the fact of post-surgical upstaging, we merged data from two independent patient cohorts obtained from two clinical centers: Xiangya hospital (*Cohort 1*) and Hunan cancer hospital (*Cohort 2*). The criterion for objective selection was outlined in **Supplementary Fig. 3F**. The rates of upstaging for each center are 10.85% (23 out of 212 in *Cohort 1*) and 15.31% (214 out of 1398 in *Cohort 2*), respectively. These rates fall within the range reported by previous studies (Dabi et al., 2018; Thelissen et al., 2022). Interestingly, both cohorts of data revealed that the misdiagnose rate was significantly higher in HPV-negative patients than HPV-positive patients, especially among ADC patients. From the data of *Cohort 2*, it was observed that ADC patients were more likely to be misdiagnosed, which means that metastatic LNs were more difficult to detect in ADC patients via radiological tools. In *Cohort 1*, we have also observed the same tendency although without significant difference between different histological types, probably due to a smaller sample size (**Table. 1-2**).

As one of stage IIIC-specific (**Supplementary Fig. 2B**) cell clusters, Epi_10_CYSTM1, with its representative marker gene SLC26A3, presents potential diagnostic value to predict lymph node metastasis, which has been shown above (**Fig. 3B** and **Supplementary Fig. 3E**). Therefore, we try to validate SLC26A3’s association with staging of IIIC, via detecting the expression pattern on post-surgical ADC samples, and more practically on biopsy samples. The results from IHC staining showed that ADC patients with stage IIIC had a higher expression of SLC26A3 (**Fig. 3C, D** and **Table 3, 4**). In summary, our results propose that SLC26A3 might be considered as a diagnostic marker to predict lymph node metastasis in ADC patients.

**Table 1.**
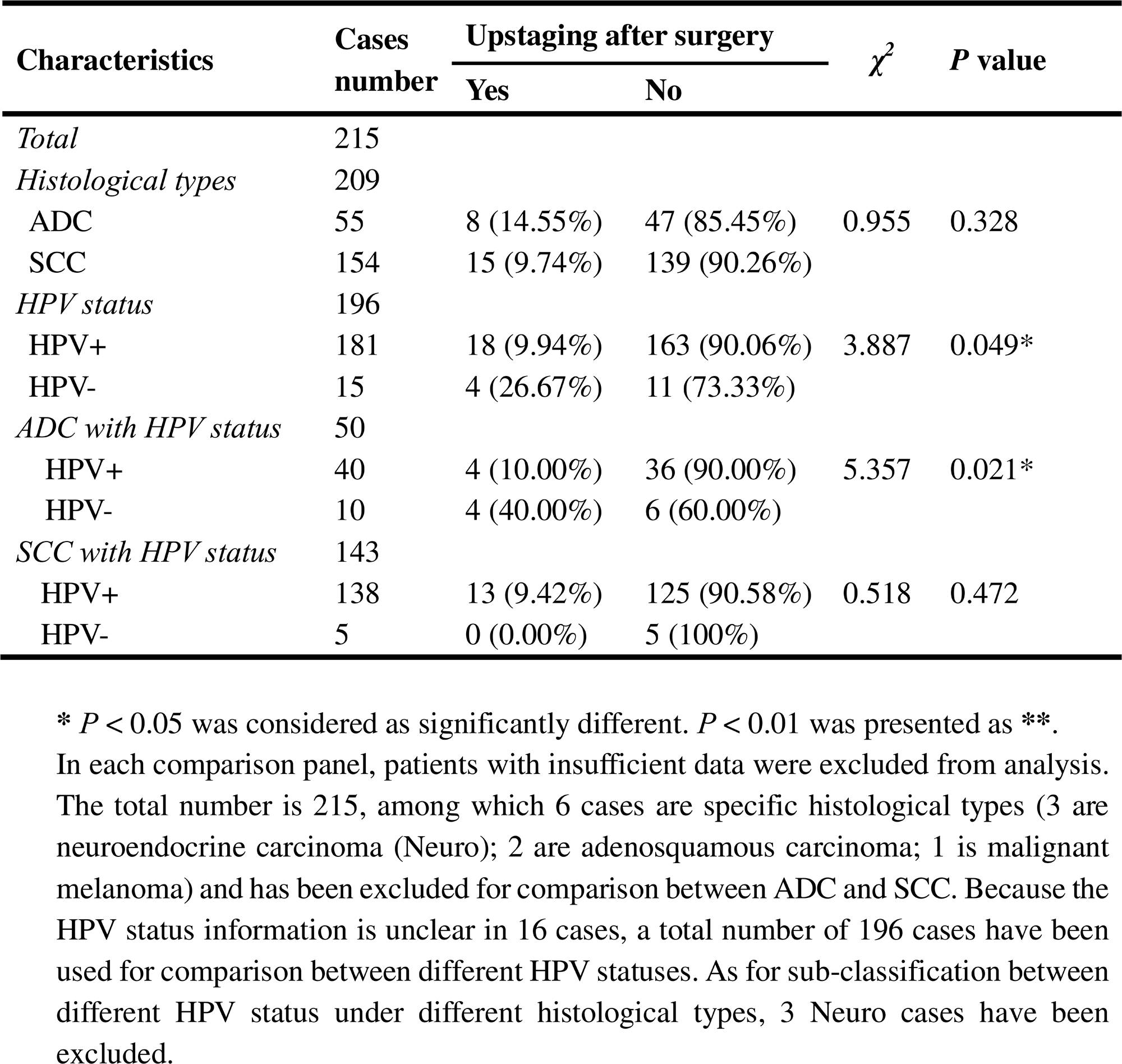
The association between post-surgical upstaging and clinical characteristics in patient cohort 1 (from Xiangya hospital)

**Table 2.**
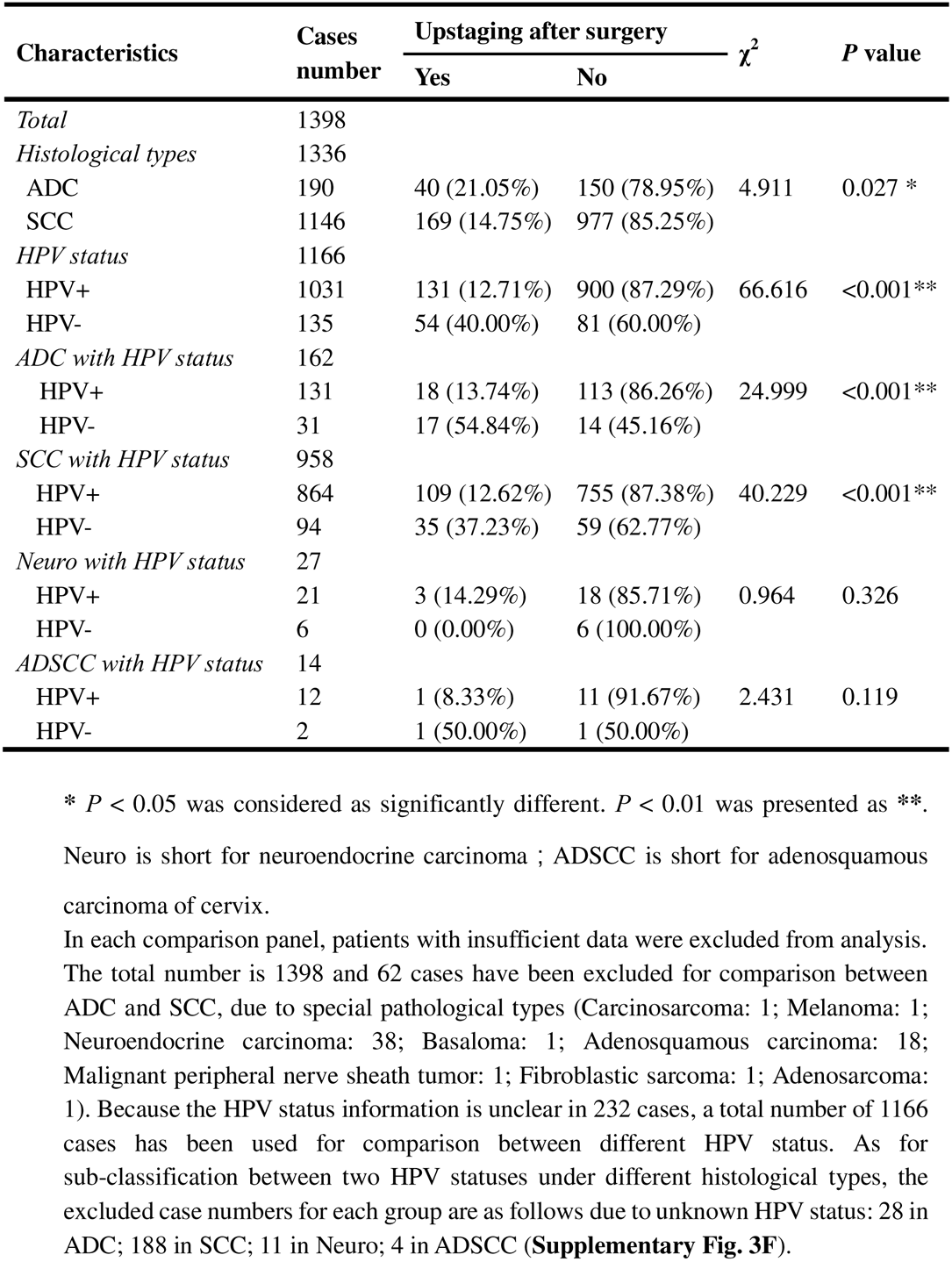
The association between post-surgical upstaging and clinical characteristics in patient cohort 2 (from Hunan cancer hospital)

**Table 3.**
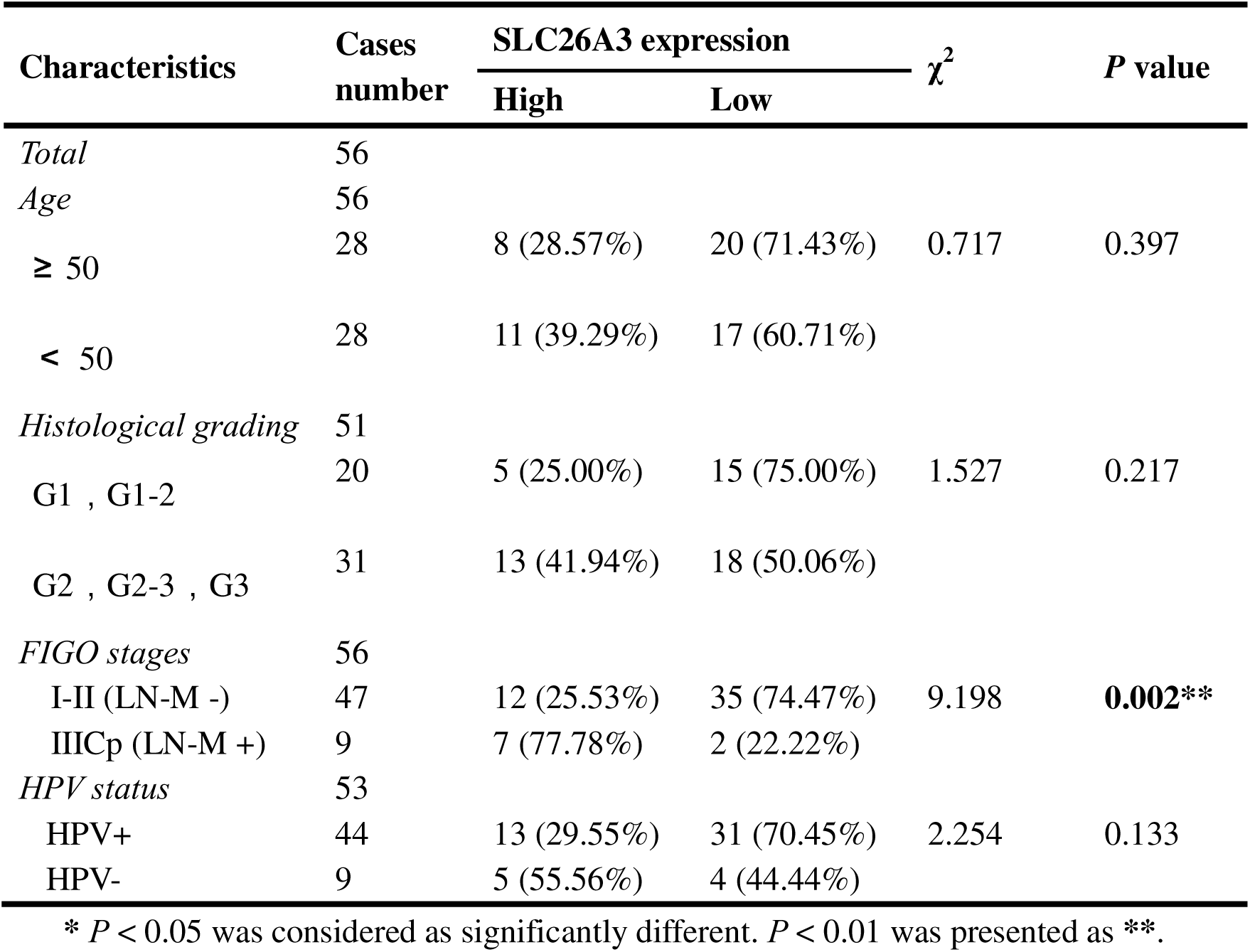
The association between clinical characteristics and SLC26A3 protein expression via IHC tested on post-surgical samples.

### Enrichment of regulatory T cells indicates an immunosuppressive status of ADC

T cells play a crucial role as the primary effectors of tumor immunity. Our data reveals that T cells constitute the predominant cell population within the TIME of cervical cancer (**Fig. 1F**). A total of 52,297 T cells were categorized into 8 main clusters (**Fig. 4A**). The gene expression patterns of T cells in CC were in accord with those observed in other types of cancers (Sorin et al., 2023; Xue et al., 2022). Based on canonical genes (C. Li et al., 2022), these cells were re-clustered as the following 4 types of T cells: exhausted T cells (marked with CD8, TIGIT, PDCD1, LAG3, HAVCR2), cytotoxic T cells (marked with CD8 and CD3), regulatory T cells (Treg, marked with CD4, FOXP3 and IL2RA), activated T cells (marked with CD8, C69 and CD3G) (**Fig. 4A-C**). The subsets of DEGs from each type of T cells have been shown in **Supplementary Fig. 4C**. When comparing the checkpoint pathway states among different T cell clusters, we observed relatively higher levels of LAG3 and PDCD1 (PD1) in exhausted T cells, while CTLA4 and TIG1 were more highly expressed in Tregs. These genes were identified to present immunosuppressive functions (Qiu et al., 2023) (**Fig. 4D**). On the other hand, genes such as ICOS and TNFRSF, which indicate the activation of immune checkpoint pathways, exhibited low expression levels in cytotoxic T cells and activated T cells (**Fig. 4D)**. These findings lead us to hypothesize that the establishment of an immunosuppressive TIME in ADC may involve the recruitment of regulatory T cells (Tregs) to the tumor area and the subsequent inactivation of a substantial proportion of cytotoxic T cells.

**Figure 4.**
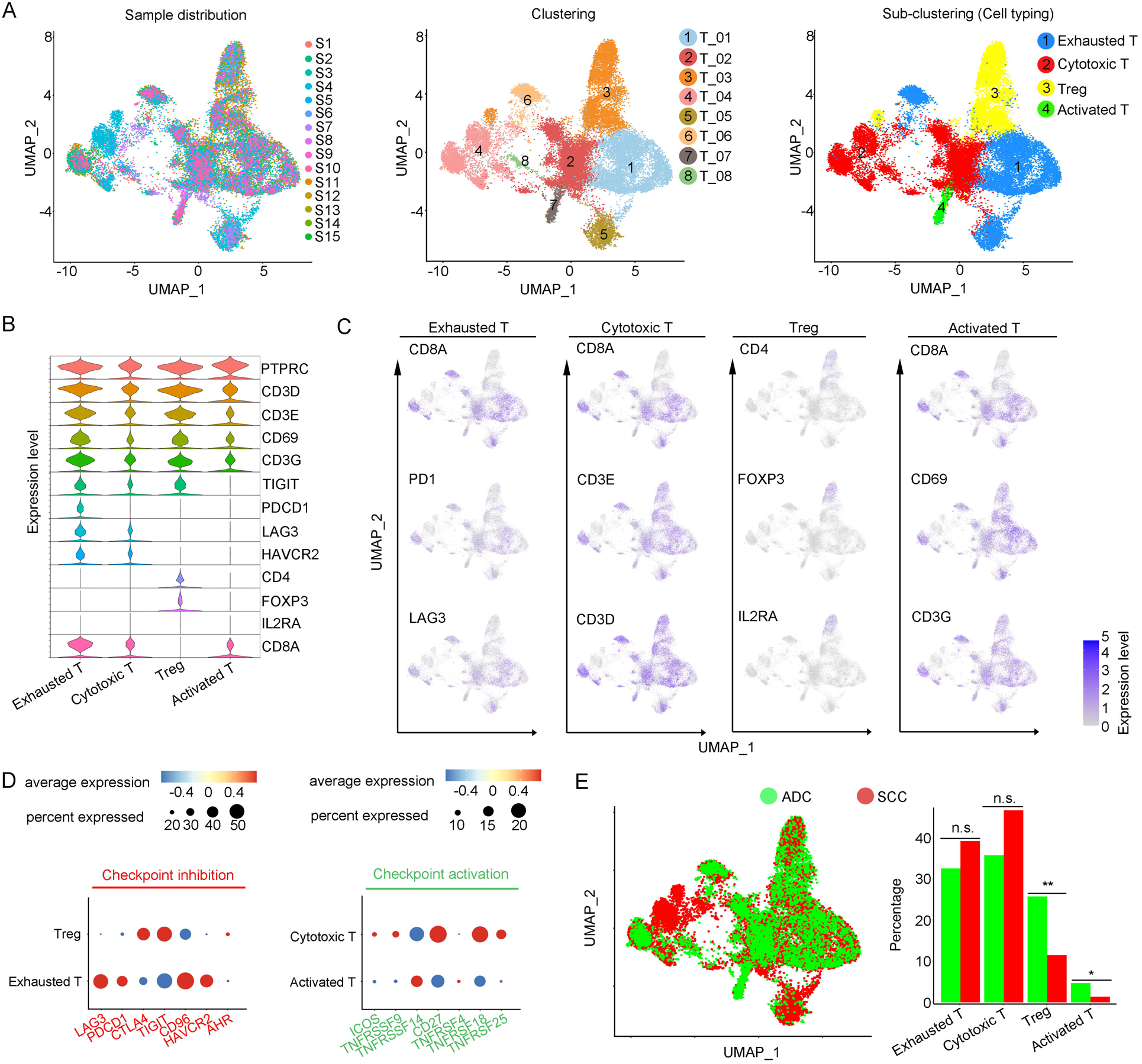
Cellular and molecular heterogeneity of T cells in ADC. **A.** UMAP plots showing the sample distribution (left), original cell clustering of T cells (middle, 8 clusters in total) and re-clustering of T cell subtypes (right, 4 major sub-clusters: exhausted T cell, cytotoxic T cell, regulatory T cell and activated T cell) according to acknowledged marker genes. **B.** Violin plot showing the expression of specific marker genes that annotate each sub-type of T cells. **C.** UMAP plots showing the widely recognized classification markers that denote each type of T cell sub-cluster. **D.** Dot heatmap plots that demonstrate the level of marker genes representing the signaling pathways of immune checkpoint activation and inhibition for grouped sub-clusters. **E.** Differences of distribution and proportion of each T cell sub-cluster between different histological types (ADC vs. SCC). Statistics were performed using R software with two-sided Wilcoxon test (*P* values for each group are listed below: Exhausted T: *P=*0.571; Cytotoxic T: *P*=0.078; Treg: *P*<0.01; Activated T: *P=*0.040). Statistics were shown as \**P* < 0.05; \*\**P* < 0.01; n.s. not significant. ***Abbreviation:*** Treg: regulatory T cell.

We further compared different status of CC with different attribution of T cell subtypes. It was demonstrated that ADC had a significantly higher proportion of Tregs compared to SCC (**Fig. 4E)**. Meanwhile, ADC patients who tested negative for HPV showed a higher enrichment of Tregs and a lower proportion of activated T cells compared to those with HPV-positive status (**Supplementary Fig. 4A**). These findings shed light on the underlying mechanism for immunotherapy insensitivity of ADC, especially in HPV-negative status.

### Tumor associated neutrophils (TANs) surrounding ADC tumor area may contribute to the formation of a malignant microenvironment

The large-scale enrichment of specific gene subsets in neutrophils in CC prompted us to further examine their roles. A total of 12,586 neutrophils were detected and the composition proportion in ADC (20.05%) was almost five times higher than that in SCC (4.23%) (**Fig. 1F**). Based on this observation, we hypothesize that ADC-specific neutrophil clusters may be related to poor prognosis and increased malignancy. All the cells were initially grouped into 6 clusters based on the gene expression modules (**Fig. 5A**). In the context of cancer, tumor associated neutrophils (TANs) can exhibit dual functions, either promoting tumor (pro-tumor TANs) or inhibiting tumor progression (anti-tumor TANs). Based on the consensus markers identified by *Jaillon et al.* (Jaillon et al., 2020) and *Xue et al.*(Xue et al., 2022), we performed further clustering of the neutrophils in our dataset, resulting in the identification of 3 sub-types of TANs: pro-tumor TANs (Neu_03, marked with CD66, CD11 and MT1X), anti-tumor TANs (Neu_02, marked with CD66, CD11, CCL5, KRT5 and ELF3), TANs with isg (interferon stimulated genes) and other undefined types (Neu_01, distinguished by CD66, CD11, IFITM2, S100A8 and LST1) (**Fig. 5B-C**). Generally, the Neu_01 sub-cluster, which consists of TANs with isg with the subset of specific genes, accounted for the largest proportion among all neutrophils. Notably, the presence of pro-tumor TANs was significantly higher in ADC compared to SCC (**Fig. 5D**), as well as in HPV-negative cases compared to HPV-positive cases (**Supplementary Fig. 5A**). On the other hand, the proportion of anti-tumor TANs in ADC was predominantly lower than that in SCC (**Fig. 5D**).

**Figure 5.**
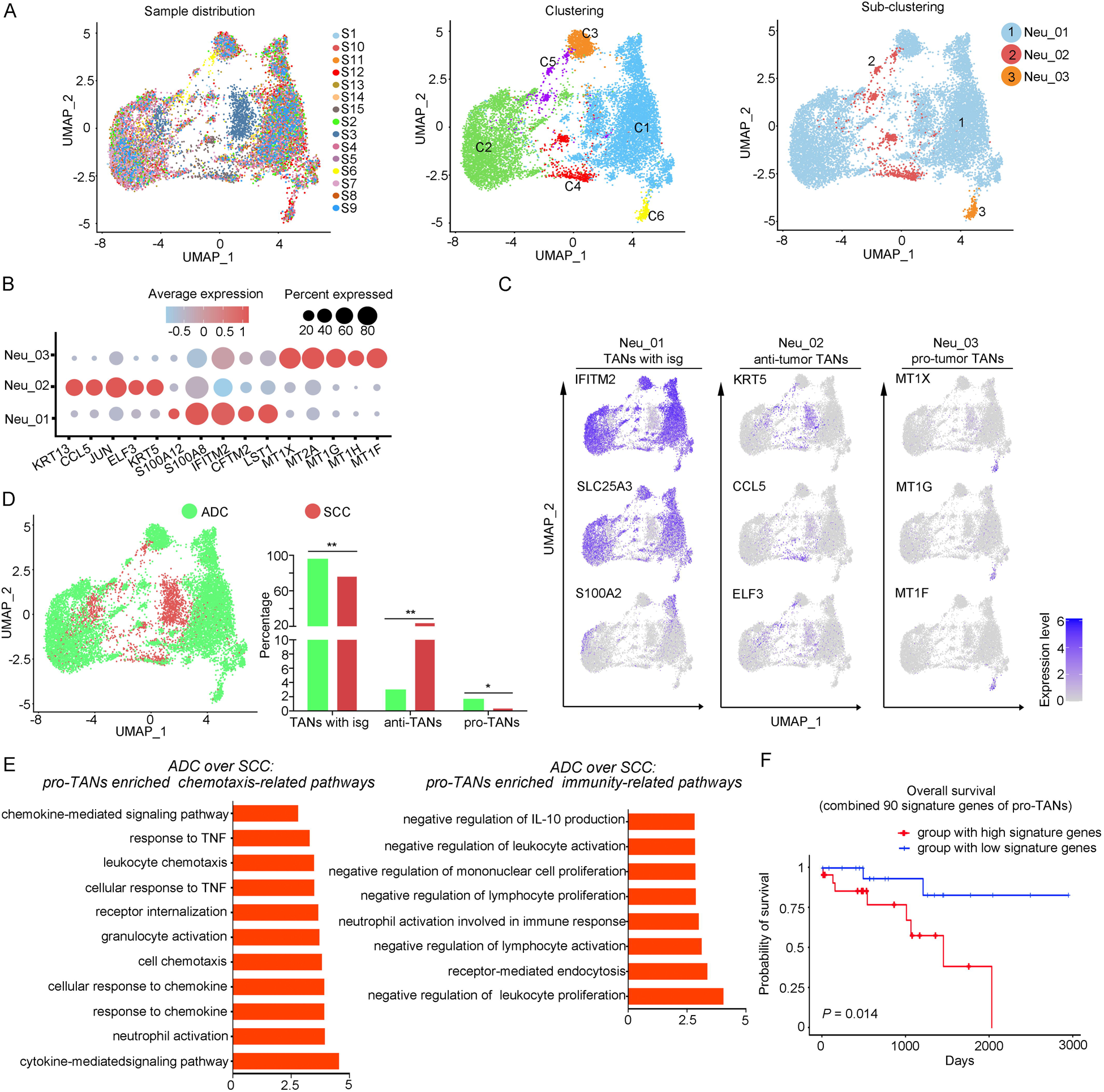
The heterogeneity of tumor associated neutrophils in ADC. **A.** UMAP plots showing the distribution of neutrophils in 15 samples (left), original clustering (middle) the re-clustering (right) into 3 sub-clusters according to gene markers of tumor-associated neutrophils (TANs). **B.** Dot heatmap showing the top 5 DEGs in each sub-cluster of TANs. **C.** UMAP plots for annotation of each sub-type of TANs with published marker genes presented: pro-tumor TANs, anti-tumor TANs and TANs with isg. **D.** UMAP and histogram plots to compare the differences of distribution and proportion of each sub-types of TANs between different histological types (ADC vs. SCC). Statistics were performed using R software with two-sided Wilcoxon test (*P* values for each group are listed below: TANs with isg: *P=*0.002; anti-TANs: *P*=0.002; pro-TANs: *P*=0.049). Statistics were shown as \**P* < 0.05; \*\**P* < 0.01; n.s. not significant. **E.** GOBP analyses of signaling pathways that are more active in ADC than SCC, in terms of the pro-tumor TANs cluster. The histogram data are transformed from -Log_10_ (*P*-value). **F.** Kaplan-Meier curve showing the overall survival rate of CC patients stratified by the top 90 genes-scaled signature of pro-TANs.

We further conducted the GOBP analysis on gene subsets derived from pro-tumor TANs to predict their potential functions. The findings indicated that pro-tumor TANs in ADC might be more active in regulating cell-to-cell interactions and chemotaxis. Specifically, they were involved in pathways such as chemokine-mediated or cytokine-mediated signaling, neutrophil activation, and receptor internalization (**Fig. 5E**). These results suggest that pro-tumor TANs may engage in communication with various cell types. Interestingly, pro-tumor TANs were predicted to acquire enhanced activation in mediating inhibitory pathways related to immunity (**Fig. 5E**). These pathways include the negative regulation of leukocyte proliferation and lymphocyte activation, as well as the suppression of IL-10 production, which is known to promote immune responses (Saraiva, Vieira, & O’Garra, 2020; Sarvaria, Madrigal, & Saudemont, 2017).

The role of pro-tumor TANs in association with disease prognosis was further investigated using TCGA database. The patients with higher expression of gene panels specific to pro-tumor TANs exhibited a poorer prognosis compared to those with lower expression (**Fig. 5F**). These findings suggest that the increased malignancy observed in ADC, particularly in HPV-negative cases, may also be related to the enrichment of pro-tumor TANs.

### Cellular heterogeneity of plasma/B cells in ADC

In our study, we identified 6 distinctive clusters of plasma/B cells based on the set of gene enrichment: Plasma/B_01(marked with IGHA2, IGHG2 and IGLC2), Plasma/B_02 (marked with HLA-DRA, HLA-DPB and CD37), Plasma/B_03 (marked with KRT17 and S100A2), Plasma/B_04 (marked with CCL5, CXCL13, IL-32 and CCL4), Plasma/B_05 (marked with CXCL8), Plasma/B_06 (marked with HMGB2) (**Fig. 6A-C**). Among these clusters, the most abundant one was Plasma/B_01_IGHA2 (**Fig. 6D** and **Supplementary Fig. 6A-B**). Interestingly, the proportion of Plasma/B_01_IGHA2 was found to be higher in ADC compared to SCC, and it was also slightly higher in HPV-negative cases than in HPV-positive cases (**Fig. 6D** and **Supplementary Fig. 6A**). Generally, B-lymphocytes are known to have tumor-inhibitory properties. However, the ectopic gene module of plasma/B cells in ADC suggested that some sub-clusters might present a tumor-promoting role. When we combined the top 50 signature genes in Plasma/B_01_IGHA2-enriched subset and performed the survival analysis using TCGA database, we observed that higher expression of the signature genes predicted a poorer prognosis of CC patients (**Fig. 6E**). Taken together, the cellular heterogeneity of plasma/B cells in the TIME of ADC is identified at the single-cell level.

**Figure 6.**
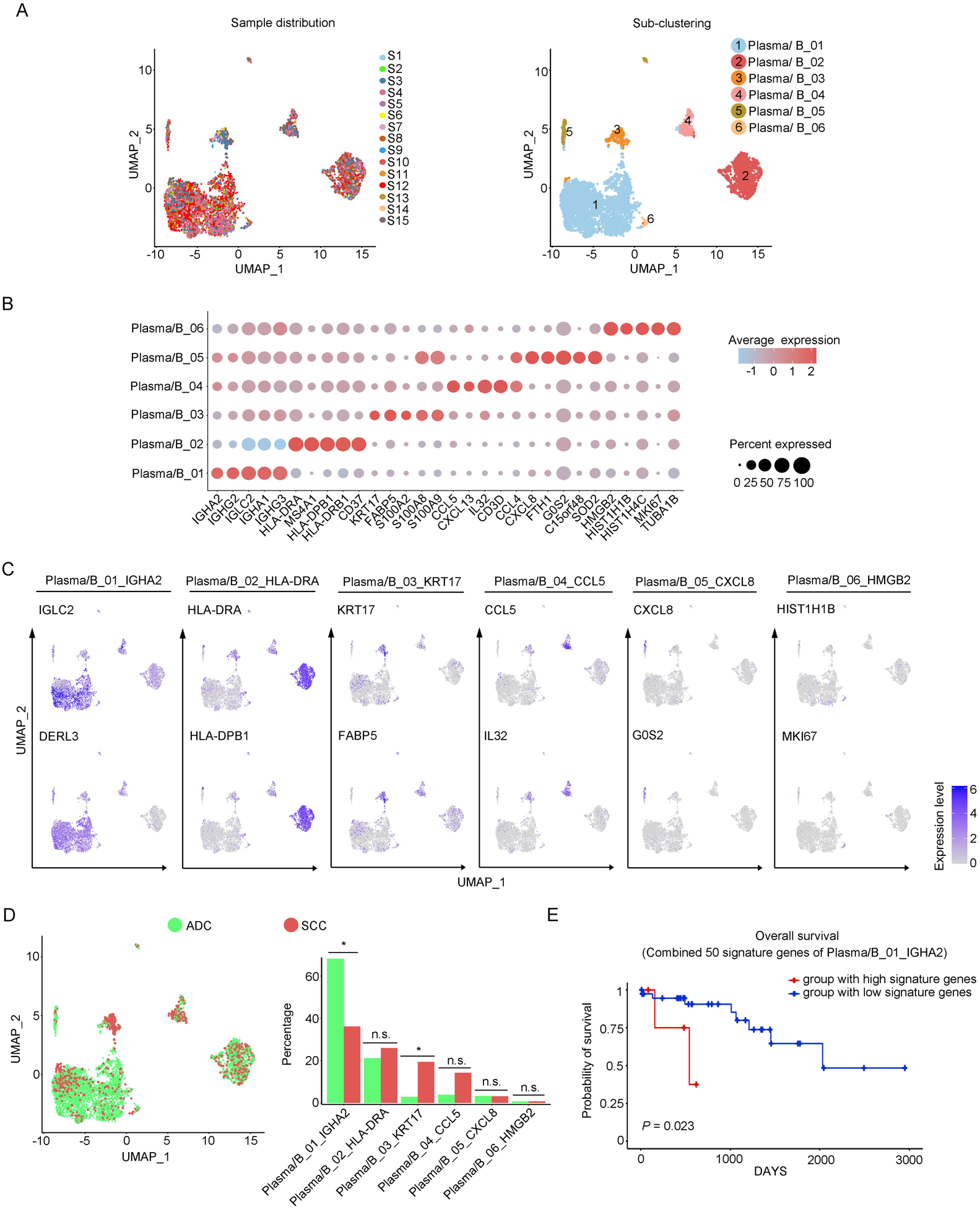
Phenotype diversity of plasma/B cells in ADC. **A.** UMAP plots showing the distribution of plasma/B cells in 15 samples (left) and the re-clustering into 6 clusters (right) according to ectopic expressions of genes. **B.** Plot heatmap showing the annotation of each sub-cluster with top 5 DEGs. **C.** UMAP plots demonstrating the most specific genes as the marker for each sub-cluster. **D.** UMAP and histogram plots to compare the differences of distribution and proportion of each plasma/B cell sub-cluster between different histological types. Statistics were performed using R software with two-sided Wilcoxon test (*P* values for each group are listed below: Plasma/B_01_IGHA2: *P=*0.040; Plasma/B_02_HLA-DAR: *P=*0.177; Plasma/B_03_KRT17: *P=*0.040; Plasma/B_04_CCL5: *P=*0.388; Plasma/B_05_CXCL8: *P=*0.169; Plasma/B_06_HMGB2: *P=*0.211). Statistics were shown as \**P* < 0.05; \*\**P* < 0.01; n.s. not significant. **E.** Kaplan-Meier curve showing the overall survival rate of CC patients stratified by the top 50 genes-scaled signature of Plasma/B_01_IGHA2.

### Crosstalk among tumor cells, Tregs and neutrophils establishes the immunosuppressive TIME in ADC

In the above description, we have outlined that certain ADC-enriched cell sub-clusters, such as Epi_10_CYSTM1, Tregs, and pro-tumor TANs, might play oncogenic roles in regulating various cellular activities, including maintenance of stemness and evasion of immune surveillance. To gain deeper insights into the interplay among these clusters and their influence on ADC progression, we harnessed the Cellchat tool (Jin et al., 2021). Our primary focus revolved around unraveling the mechanisms governing Treg recruitment to the tumor microenvironment and the maintenance of stemness within ADC-enriched cell sub-clusters.

Given that the 3 ADC-specific sub-clusters (Epi_07_CAPS, Epi_10_CYSTM1 and Epi_12_RRAD) of epithelial cells were predicted to exhibit high levels of stem-like properties, and particularly that Epi_10_CYSTM1 presented the highest degree of malignancy among them, we further analyzed their possible interactions with other types of cells. Firstly, in comparison to SCC, ADC-enriched epithelial sub-clusters exhibited a higher propensity for interaction with Tregs through ligand-receptor signaling pathways, including ALCAM (ALCAM>>CD6) and MHC-II (HLA-DR>>CD4) (**Fig. 7A-E**), which have been reported to be essential for Treg recruitment, expansion and stabilization to establish the immunosuppressive microenvironment (Chalmers et al., 2022; Ferragut, Vachetta, Troncoso, Rabinovich, & Elola, 2021; Freitas et al., 2019). To further uncover the underlying reason for the enhanced stemness observed in ADC epithelial cells, we focused on the signaling received by Epi_10_CYSTM1 cell cluster. Interestingly, the Cellchat analysis revealed that, compared to other pathways, such as CD46>>JAG1 and GZMA>>PARD3, the interaction of TGFβ>>TGFBR between Tregs and epithelial cells (particularly in cluster Epi_10_CYSTM1) was activated in ADC but not in SCC (**Fig. 7F-G**). TGFβ is a canonical secreted protein that induces carcinogenesis, such as cancer cell proliferation, invasion, self-renewal and EMT (epithelial-to-mesenchymal transition) (Derynck, Turley, & Akhurst, 2021; Massagué, 2008; Tauriello, Sancho, & Batlle, 2022). Thus, we speculate that the recruited Tregs might secrete TGFβ to stimulate the tumor epithelial cells of ADC, leading to increased stemness. Additionally, pro-tumor TANs were also recruited to the tumor area and interacted with Epi_10_CYSTM1, as well as Epi_07_CAPS and Epi_12_RRAD cells, via the ANNEXIN-to-FPR1/FPR2 signaling pathway (**Supplementary Fig. 7A-B**). Previous studies have reported that the ANNEXIN pathway can promote the chemotaxis of neutrophils (Araújo et al., 2021). Likewise, TANs expressing isg, which have been shown to promote tumorigenesis as mentioned earlier (**Supplementary Fig. 5C**), were also recruited to the tumor area through the ANNEXIN pathway. These findings suggest that the aggressive clusters of TANs may contribute to the formation of an ADC-specific TIME.

**Figure 7.**
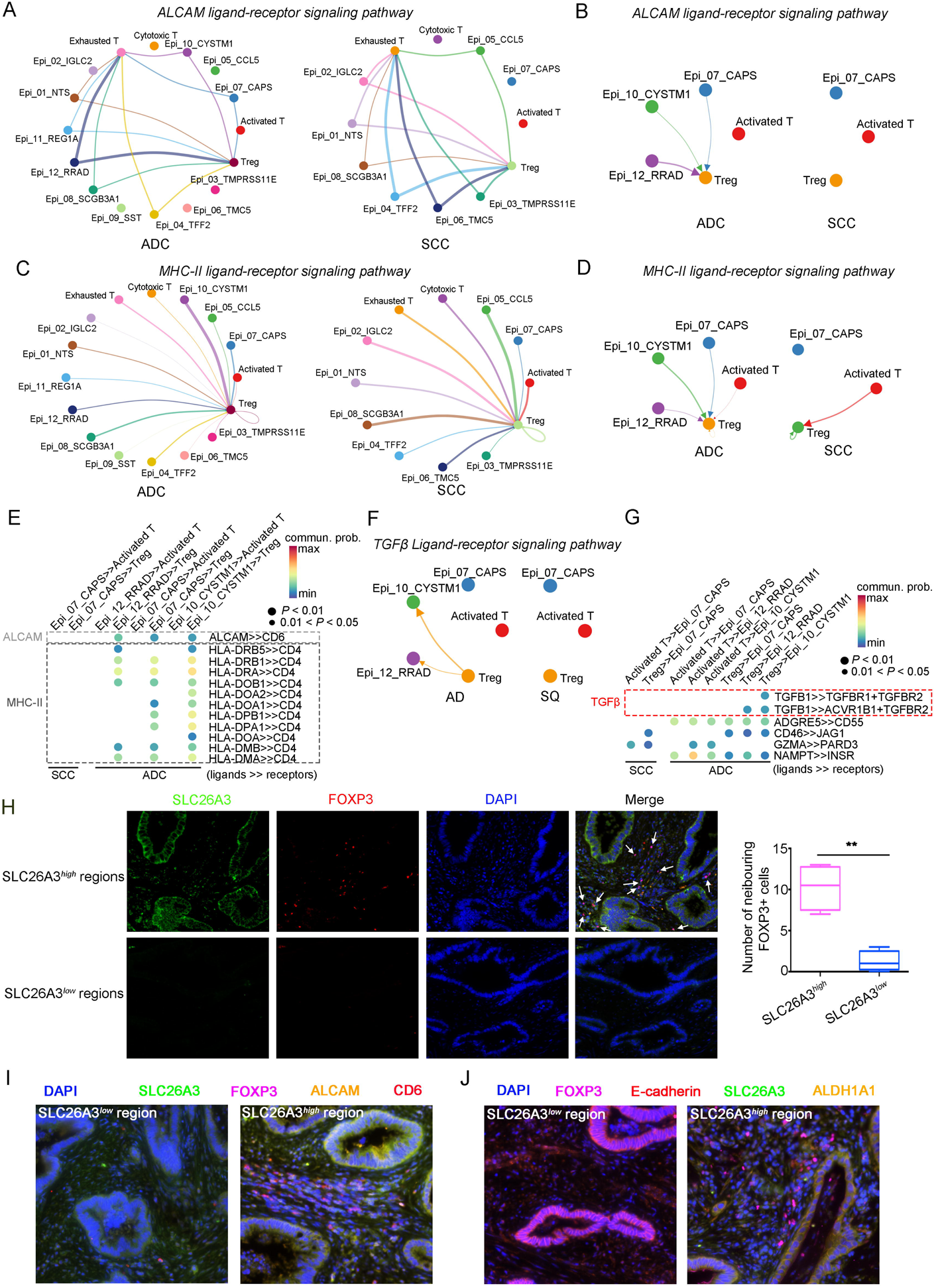
The cellular interaction modules of sub-clusters of T cells, neutrophils and tumor epithelial cells. **A.** Circle plots showing the interacting networks between epithelial cell sub-clusters and T cell sub-clusters via the pathway of ALCAM, by comparing ADC and SCC. **B.** Circle plots simplified from **A** to show the interactions among target cell clusters via ALCAM pathway. **C.** Circle plots showing the interacting networks between epithelial cell sub-clusters and T cell sub-clusters via the pathway of MHC-II, by comparing ADC and SCC. **D.** Circle plots simplified from **C** to show the interactions among target cell clusters via MHC-II pathway. From **A** to **D**, The direction of each arrow shows the regulation from outputting cells to incoming cells. The width of the each line shows the predicted weight and strength of regulation. **E.** Bubble plot showing the probability of ligand-to-receptor combination of each pathway between two different target sub-clusters of cells, by comparing ADC with SCC. **F.** Circle plots showing Tregs regulate epithelial cells via the TGF-β pathway, which is solely activated in ADC. **G.** Bubble plot showing the probability of ligand-to-receptor combination of TGF-β pathway between Tregs and epithelial cells, by comparing ADC with SCC. The pathways of ADGRE5, CD46, GZMA and NAMPT are used as negative controls**. H.** Dual IF staining confirming that in SLC26A3^high^ regions of CC tissues, more FOXP3^+^ cells are recruited than in SLC26A3^low^ regions (left). The numbers of recruited FOXP3^+^ cell are quantified using histogram plot (right). Three individual samples with ROI were calculated and *P*<0.01 were marked with **, showing significant difference. **I.** Multiplexed IF staining confirming the interaction between CD6 (on FOXP3^+^ cells) and ALCAM (on SLC26A3^high^ epithelial cells) in the ALCAM pathway. **J.** Multiplexed IF staining showing that the recruitment of FOXP3+ cells towards SLC26A3^high^ cells might induce EMT (marked with E-cadherin) and increase the stemness (marked with ALDH1A1) of tumor cells, via TGF-β pathway.

To test these hypotheses, we conducted immunofluorescence (IF) analysis and observed a greater recruitment of Tregs (marked by FOXP3) in close proximity to regions exhibiting high SLC26A3 expression (Epi_10_CYSTM1 cell cluster) (**Fig. 7H**). Furthermore, when compared to regions with low SLC26A3 expression, tumors displaying high SLC26A3 levels exhibited significantly increased expression of Aldehyde Dehydrogenase 1 Family Member A1 (ALDH1A1), a known marker for cancer stem cells in CC (Douville, Beaulieu, & Balicki, 2009) (**Supplementary Fig. 7C**). Notably, we identified SLC26A3^high^ tumor cells expressing ALCAM in close proximity to CD6+ Tregs, suggesting potential involvement of ALCAM>>CD6 signaling in Treg recruitment (**Fig. 7I**). Additionally, we observed reduced E-cadherin expression in tumor cells within SLC26A3^high^ regions, possibly induced by TGFβ secreted by adjacent Tregs (**Fig. 7J** and **Supplementary Fig. 7D**). Altogether, these findings confirm the intricate crosstalk between the Epi_10_CYSTM1 cell cluster and Tregs, which might help to establish an immunosuppressive TIME and sustain heightened stemness in tumor cells.

## Discussion

Cervical cancer is often regarded as less aggressive compared to ovarian cancer and endometrial cancer, primarily due to the effectiveness of HPV vaccines in prevention. When diagnosed at early stages, the 5-year survival rate for cervical cancer patients exceeds 90%, surpassing that of other gynecologic malignancies (Gennigens et al., 2022; Schiffman, Castle, Jeronimo, Rodriguez, & Wacholder, 2007). However, if cervical cancer metastasizes to the intraperitoneal or retroperitoneal lymph nodes, the survival rate will decrease to 30%∼60% (Cohen et al., 2019). Worse still, the 5-year survival rate for recurrent CC patients is less than 20% (H. Li, Wu, & Cheng, 2016). Specifically, when considering the ADC type alone, the prognosis is even worse. Unfortunately, our understanding for ADC is limited. Most previous studies have concentrated on SCC type of CC, and obtained heterogenetic information derived from bulk RNA-seq method. Therefore, it is worthwhile to shift our attention towards an in-depth investigation on the disparities between ADC and SCC at the single-cell resolution.

More recently, researchers have started to utilize scRNA-seq approach to investigate the tumor microenvironment of CC. For instance, Cao *et al* compared 5 cases of SCC tumor tissues with paired adjacent normal tissues and described the immune landscape in which specific clusters of T and B cells exhibited immune-exhausting or activating processes (Cao et al., 2023). Another study conducted by *Qiu et al* aimed to elucidate the distinct molecular patterns of immune reactions between ADC and SCC in the context of different HPV infection status at the single-cell level (Qiu et al., 2023). However, due to limited samples tested in these studies, our knowledge on the TIME in CC is yet inadequate. In the current study, we have generated a genomic atlas of ADC, which may depict the landscape of TIME built by both ADC tumor cells and neighboring cells. Apart from the predominant presence of epithelial cells, the cell clusters in ADC are majorly composed of T cells, tumor-associated neutrophils (TANs), and plasma/B cells. The gene subsets identified within ADC-abundant cell sub-clusters, including Epi_10_CYSTM1, Tregs, pro-tumor TANs and P/B_01_IGHA2, exert potential roles in promoting both the carcinogenesis and immunosuppression of ADC. In the TIME of ADC, the unique module of cellular interaction has been preliminarily indicated that the immunosuppressive Tregs are recruited to tumor cells via chemo-attraction. Finally, our hypothesis from scRNA-seq data is further confirmed using clinical samples, providing evidence that SLC26A3 may be a potential biomarker for ADC (**Supplementary Fig. 8**).

It has been reported that immunotherapies are less effective on ADC patients, especially those who are not infected with HPV (Attademo et al., 2020; Ferrall, Lin, Roden, Hung, & Wu, 2021). As the two major components of ADC cell types in our study, T cells and epithelial cells have been implied to be the dominant regulators in the TIME due to their reciprocal interaction dynamics. We identified several ADC-specific epithelial cell clusters that may exhibit stronger stemness and aggressiveness than other clusters. Among these clusters, Epi_10_CYSTM1 seemed to have the highest level of malignancy, but the underlying mechanism behind this behavior remains unknown. The Cellchat analysis revealed that Tregs might quite actively crosstalk to Epi_10_CYSTM1. To confirm this, immunostaining for co-localization demonstrated an increased aggregation of Tregs around tumor cells exhibiting overexpression of SLC26A3. It has also been widely reported that tumor cells acquire higher stemness present more aggressive features of resistance to immunotherapy (Dianat-Moghadam et al., 2022). This study indicated that TGFβ produced by Tregs might induce the stemness of Epi_10_CYSTM1 cells by activating downstream oncogenic pathways, inducing EMT on tumor epithelial cells. Because TGFβ is a soluble protein and is hard to detect on solid tissue samples, here we only validated markers of EMT and stemness to indirectly indicate the functional existence of TGFβ, which is predicted from informatic hypothesis (Nixon, Gao, Wang, & Li, 2023). Interestingly, Epi_10_CYSTM1 cells express ALCAM, which is a membranal ligand, and bind to its receptor CD6 on Tregs, activating chemotaxis and facilitating Treg recruitment (**Fig. 7I**). The infiltration of a large number of Tregs in ADC may significantly accelerate immune evasion (**Supplementary Fig. 8**). Therefore, the presence of a higher number of Tregs infiltrating the tumor area of ADC may partially explain the increased resistance to immunotherapy.

Another interesting phenomenon observed in our sing-cell sequencing data is the involvement of TANs in the regulatory interactions within the TIME of ADC. It has been clearly investigated that pro-tumor TANs, characterized by CD66, CD11, CD170 and PD-L1, exert the oncogenic roles by promoting tumor cell proliferation, angiogenesis, inducing genetic instability, and most importantly causing immunosuppression (Jaillon et al., 2020). In this study, pro-tumor TANs are relatively enriched in ADC tissues compared to SCC, especially in HPV-negative ADC cases. This suggests that ADC-specific epithelial cell sub-clusters may actively engage in the recruitment of pro-tumor TANs.

The anti-tumor function of B cells has been identified previously (Bod et al., 2023; Helmink et al., 2020). However, our study has presented a specific cluster of plasma/B cells with tumor-promoting functions. The overexpression of gene signatures from Plasma/B_01_IGH2 cells is associated with poor regression in CC patients (**Fig. 6F**). Interestingly, Plasma/B_01-IGHA2 was found to be closely related to the PC Cluster (Plasma cell Cluster) in Cao’s study due to the shared gene set (Cao et al., 2023), such as IGHA1, IGHA2, IGHG2, IGHG4, *etc.* However, their roles were contradictory. According to that study, PC cluster was derived from the local expansion of TIME and showed potential effectiveness in anti-tumor immunity. In our case, Plasma/B_01-IGHA2 was indicated to have a reverse effect and played a tumor-promoting role in ADC. This difference in function is likely attributed to the distinct cell types and their interactions in ADC, instead of SCC. However, the underneath dynamics of B cells in ADC still needs further investigation.

Owing to post-surgical upstaging, some CC patients with stage IIIC are misdiagnosed as early-stage and have undergone radical surgeries. For these patients, the most appropriate strategy of treatment should be radiotherapy alone. However, it is challenging to avoid this issue because it is difficult to identify metastatic lymph node only through CT or MRI. To solve this problem, a reliable biomarker is required to diagnose lymph node metastasis in conjunction with radiographic tool. In this study, SLC26A3 has shown its potential as a diagnostic biomarker to predict lymph node metastasis in ADC patients. Combining two large cohorts of clinical data, we found that CC patients with ADC types were more vulnerable to be misdiagnosed for clinical staging. Derived from scRNA-seq data, SLC26A3 could be employed as a representative gene for the Epi_10_CYSTM1 cluster, which was found to be enriched in stage IIIC cases of ADC. The IHC staining further confirmed that ADC patients at stage IIIC exhibited a higher expression of SLC26A3 compared to early stages (FIGO stage I∼ II). Furthermore, we tested on biopsy samples and it implied that SLC26A3 might serve as a potential predictor of LN metastasis. Our study for the first time provides evidence that ADC patients are at a higher risk of developing lymph node metastasis, which can be challenging to detect solely via radiological imaging techniques. However, by incorporating lymph node imaging with biomarkers such as SLC26A3, it may probably reduce the misdiagnosis rate among FIGO IIIC patients.

In this study, our objective is to unravel the genomic characteristics of ADC in the hope of identifying gene signatures that can elucidate its aggressive nature. SLC26A3 is selected from the DEGs enriched in cluster Epi_10_CYSTM1, and is identified to specifically represent this ADC-specific cluster. However, based on our data, we have inevitably encountered two questions: (1) the existence of cluster Epi_10_CYSTM1; (2) the specificity of SLC26A3. As for the first question, Epi_10_CYSTM1 is distinctive from the other 11 epithelial cell clusters, with specific DEGs (**Fig. 2A-C**). Although Epi_10_CYSTM1 is dominantly derived from sample S7, however, its proportion is relatively high when compared with other clusters such as Epi_08_SCGB3A1, Epi_09_SST and Epi_11_REG1A (**Fig 2D** and **Supplementary Fig. 2A-B**). Moreover, we have detected a positive expression of this cluster on tissue samples, whether by using ORM1/2 or SLC26A3 antibodies (two candidate markers of cluster Epi_10_CYSTM1) via IHC. As a result, we have identified the existence of this cell cluster. As for the second question, SLC26A3 has demonstrated a higher specificity compared to other candidates, such as CYSTM1, CA9, ORM1 and ORM2, following the process of identifying markers (**Fig. 3A-B** and **Supplementary Fig. 3E**). Although the IHC results indicate that SLC26A3 is positively expressed in stage IIIC cases, however, the expression is confined to specific regions (**Supplementary Fig. 9**), which aligns with the scRNA-seq findings for SLC26A3. Notably, according to **Table 3** and **Table 4**, most of the clinical cases have a lower expression of SLC26A3. Taken together, SLC26A3 is a specific representative of cluster Epi_10_CYSTM1.

**Table 4.**
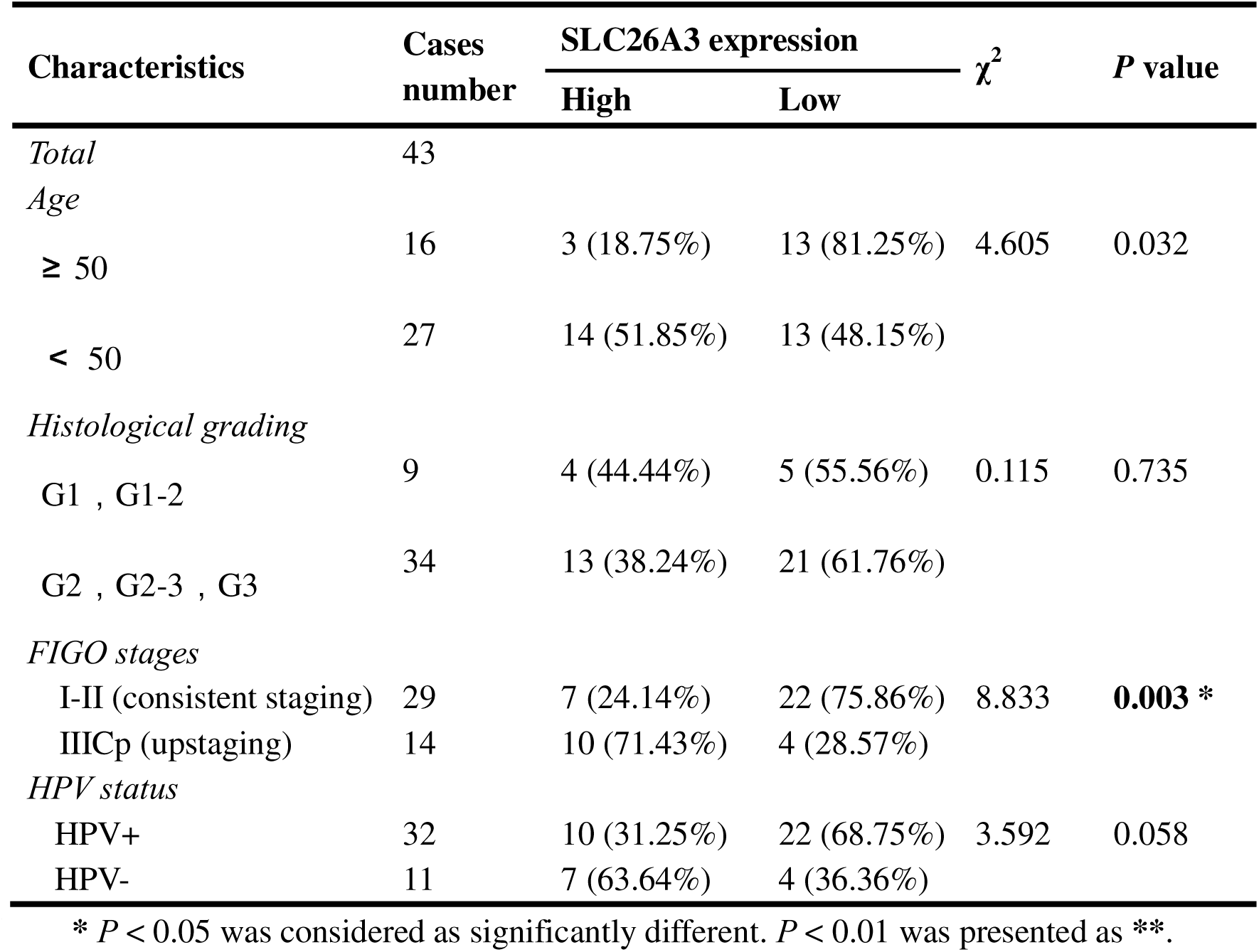
The association between clinical characteristics and SLC26A3 protein expression via IHC tested on biopsy small specimens.

In summary, we characterized the genomic landscape of the TIME in ADC at a single-cell resolution. Our study revealed the presence of specific epithelial cell clusters in ADC that exhibited more aggressive features. The enriched gene signatures from these clusters may serve as potential therapeutic targets to improve the efficacy of immunotherapy. The interactions between tumor epithelial cells and other types of cells, such as T cells and neutrophils, appropriately depict a highly immunosuppressive microenvironment of ADC. Moreover, our investigation extended to the issue of post-surgical upstaging at the single-cell level, with the identification of biomarkers from specific tumor cell clusters showing potential as diagnostic indicators for the precise diagnosis of cervical cancer patients.

## Supporting information

Supplementary Data 1-annotated copies of script

## Additional information

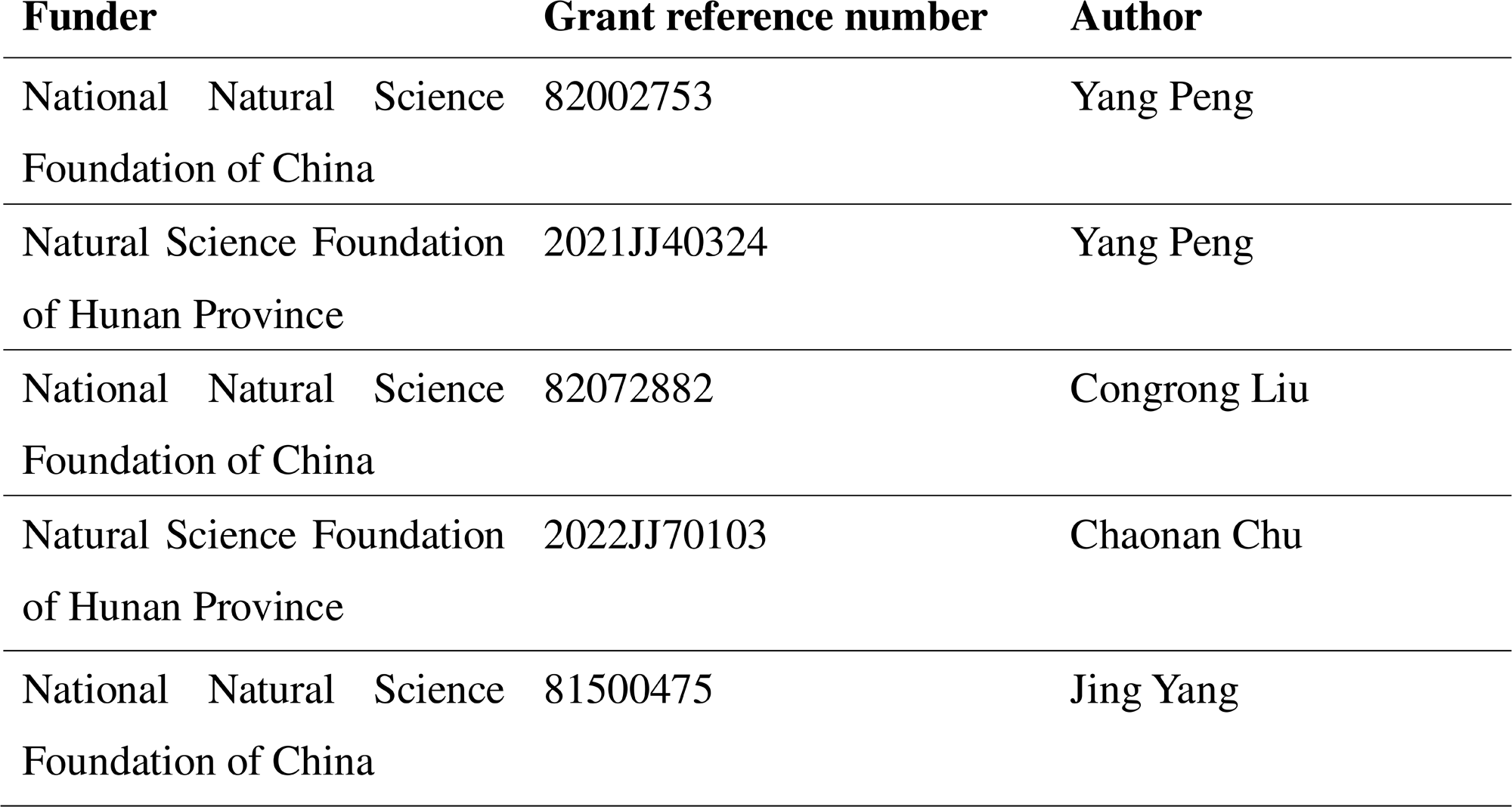

## Author contributions

Yang Peng, Conceptualization, Methodology, Formal analysis, Investigation, Data curation, Visualization, Writing – original draft preparation, Project administration, Funding acquisition; Jing Yang, Methodology, Investigation, Data curation, Writing – review & editing, Formal analysis; Jixing Ao, Methodology, Validation, Software, Resources, Investigation, Data curation; Jia Shen, Software, Validation, Investigation, Data curation, Visualization; Yilin Li, Methodology, Validation, Investigation, Resources, Data curation, Visualization; Xiang He, Software, Investigation, Data curation, Visualization; Dihong Tang, Methodology, Formal analysis, Writing – review & editing; Chaonan Chu, Formal analysis, Writing – review & editing; Congrong Liu, Conceptualization, Methodology, Formal analysis, Writing – review & editing, Supervision, Project administration, Funding acquisition; Liang Weng, Conceptualization, Methodology, Software, Validation, Formal analysis, Investigation, Writing – original draft preparation, Writing – review & editing, Supervision, Project administration.

## Ethics approval and consent to participate

Specimen collection and pathological data used in this study were approved by the Ethics Committee of Hunan Cancer Hospital (No. KYJJ-2020-036). Written informed consent was obtained from patients for use of samples. This study was also in accordance with the Declaration of Helsinki and none of any procedures conducted interfered with the treatment plan of patients.

## Data availability

In this study, the single-cell RNA sequencing data generated is available in the Genome Sequence Archive for Human (GSA-Human, from China National Center for Bioinformation) by accession number: PRJCA023028 (https://ngdc.cncb.ac.cn/gsa-human/). These original sequencing data can be obtained only for non-profitable use, under request from the corresponding authors.

