## Supplementary figures and images for "Single-cell profiling reveals the intratumor heterogeneity and immunosuppressive microenvironment in cervical adenocarcinoma"

### Supplementary Figure.1

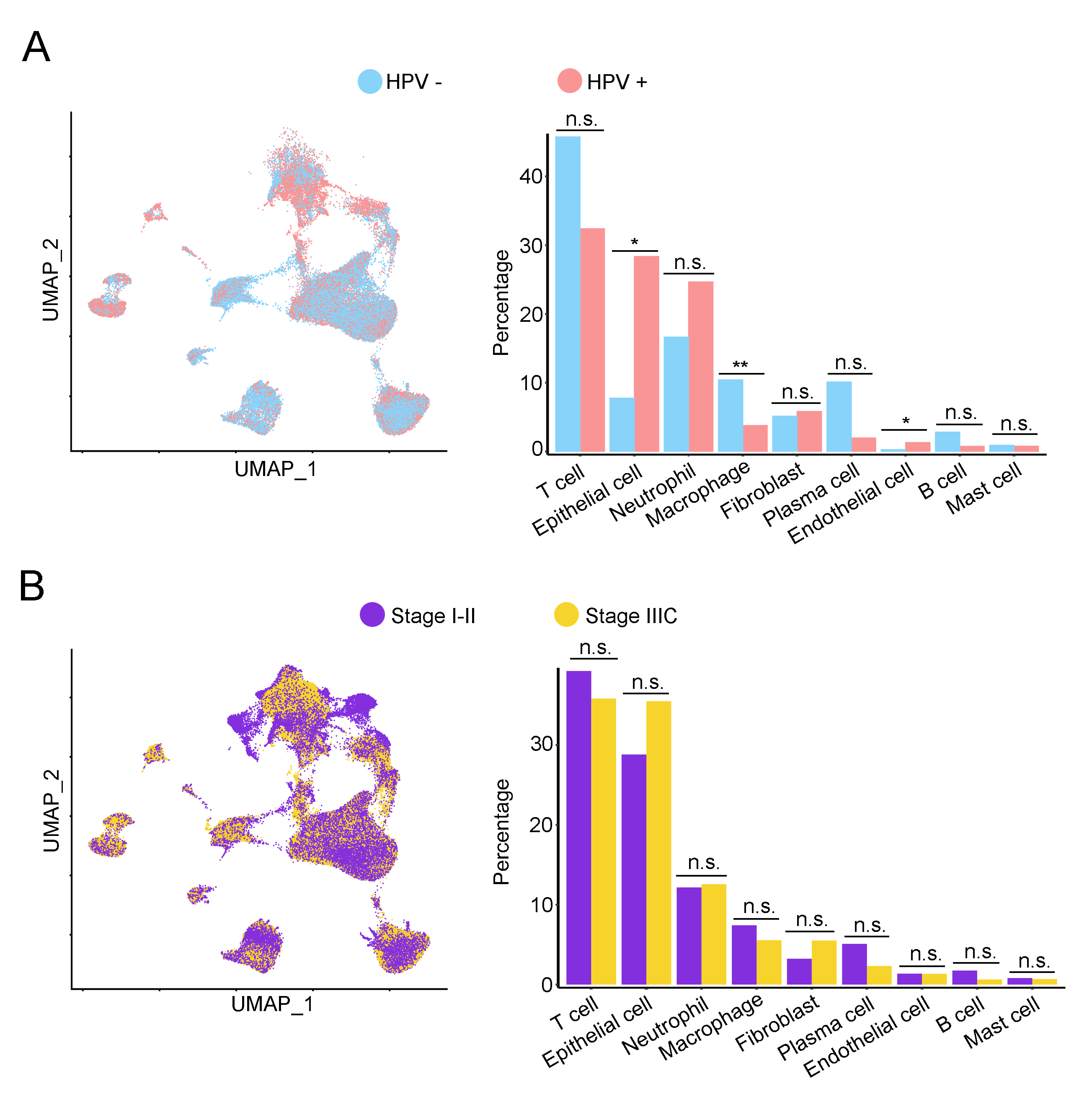

### Supplementary Figure.2

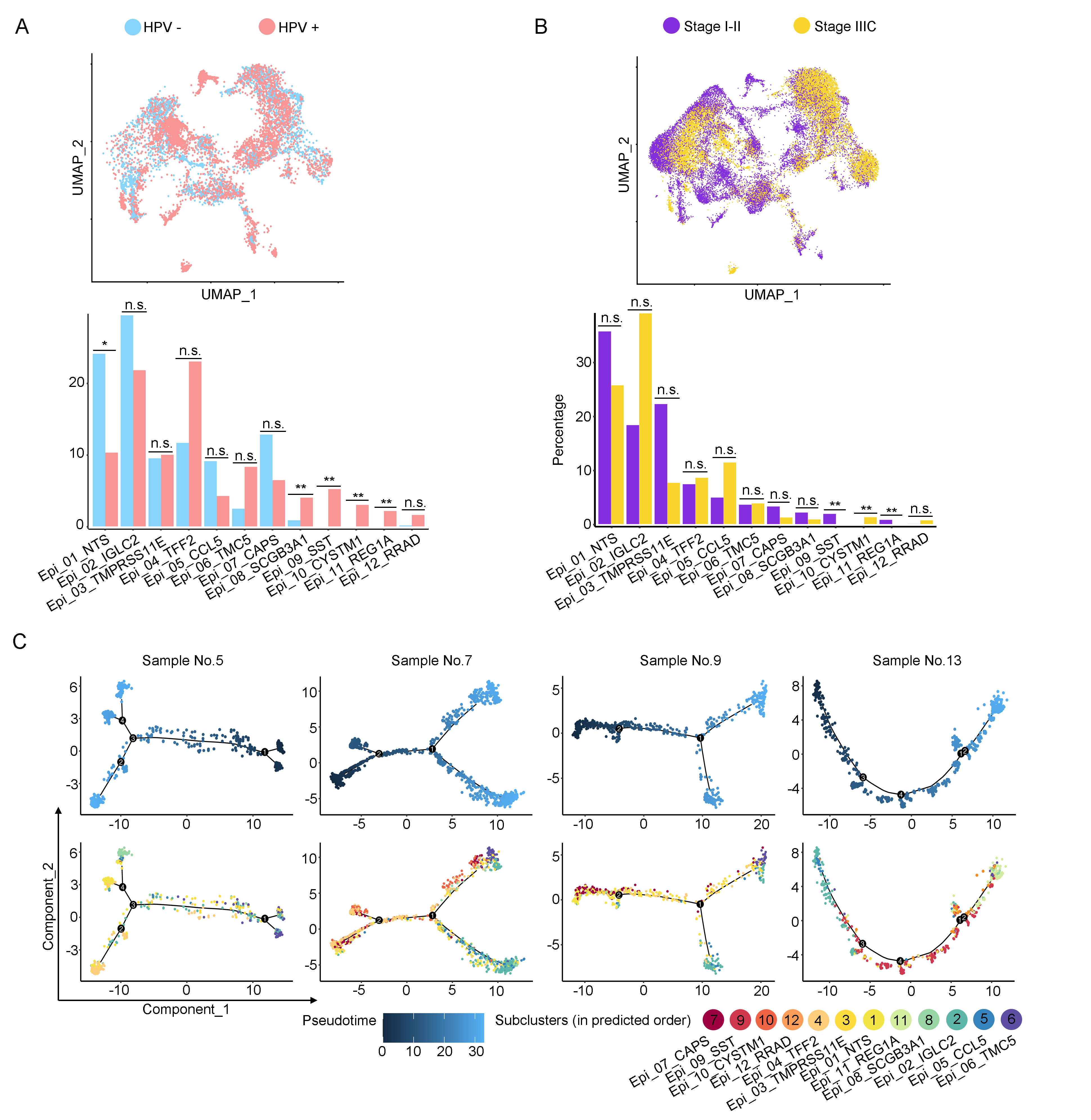

### Supplementary Figure.3

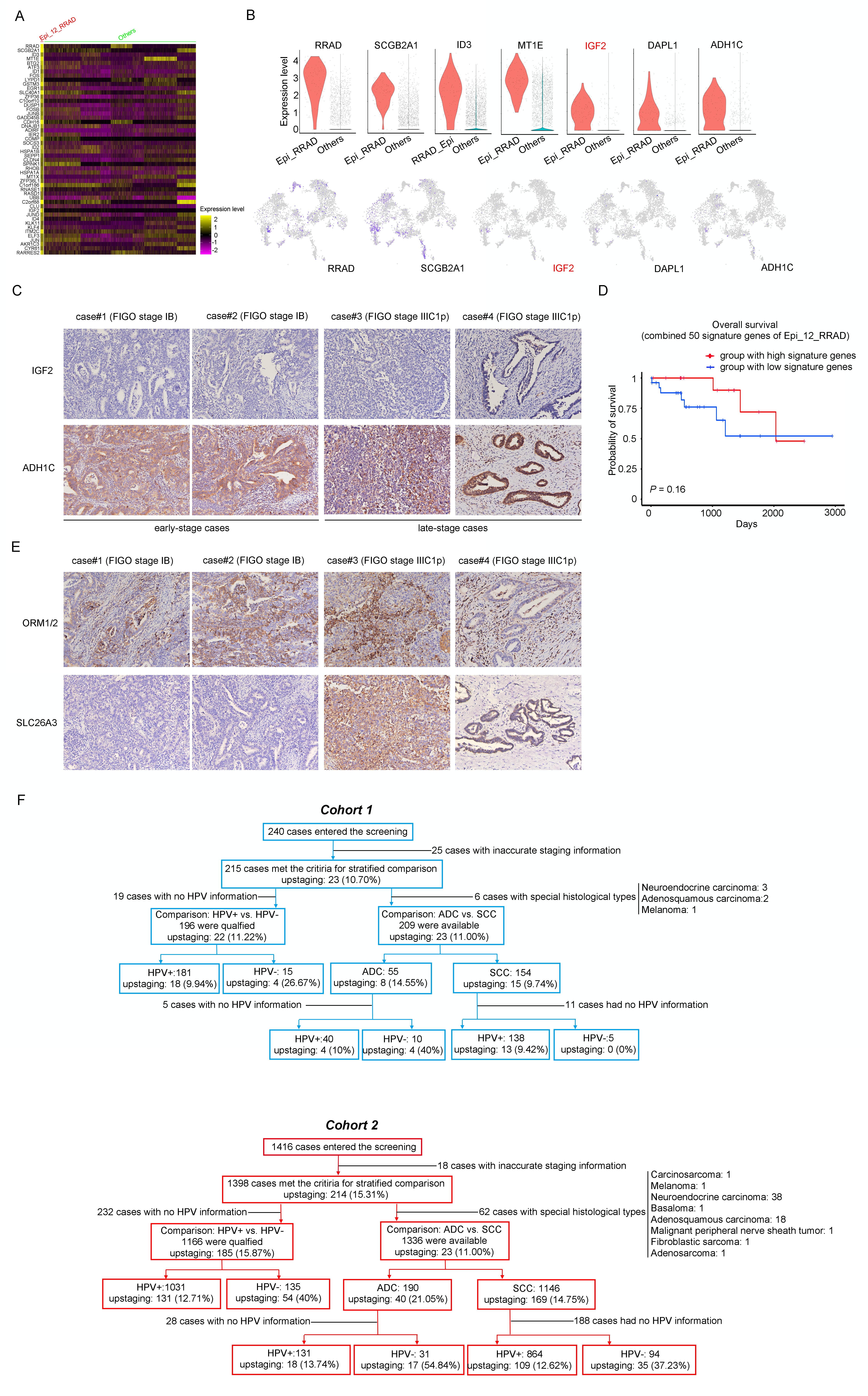

### Supplementary Figure.4

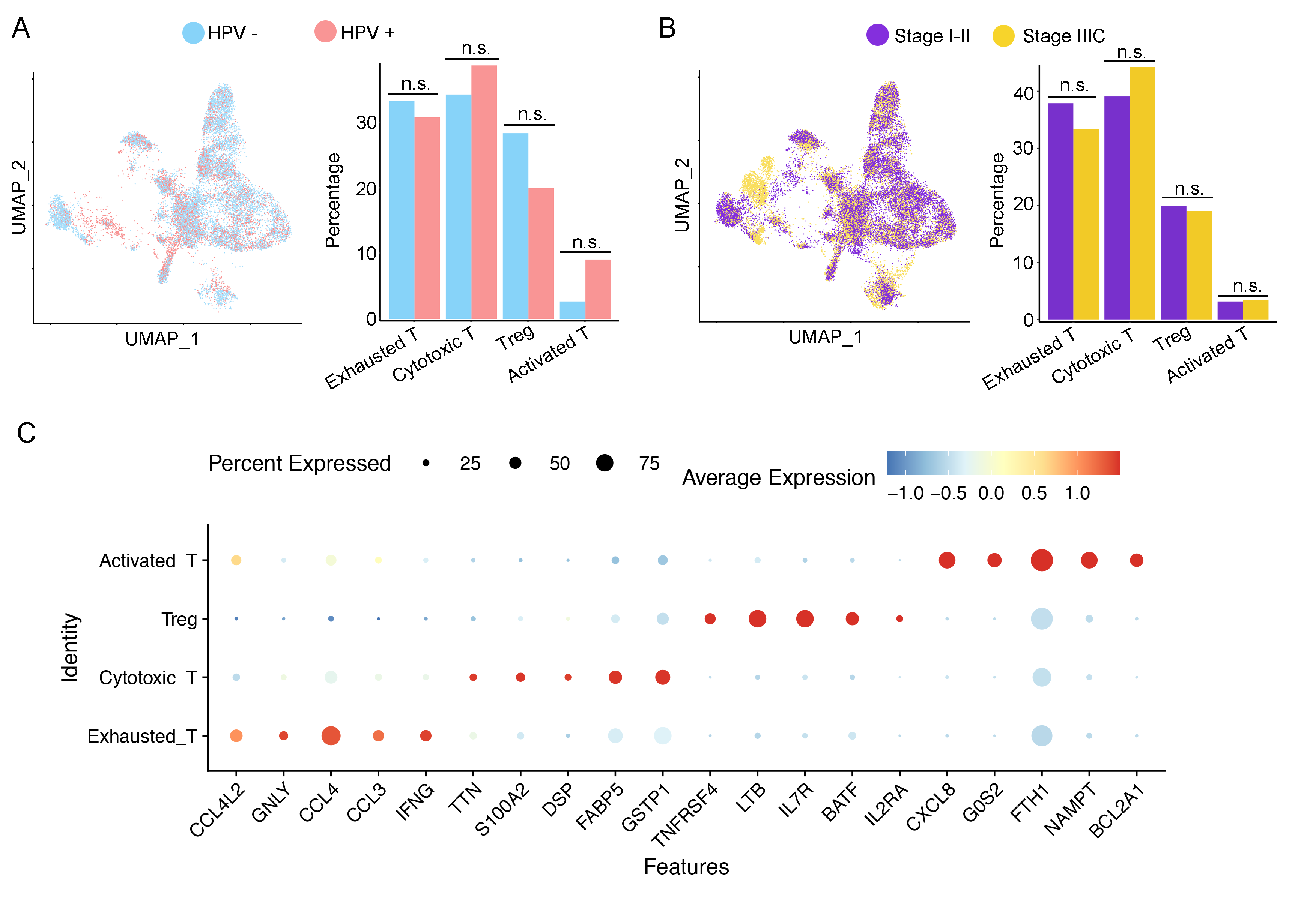

### Supplementary Figure.5

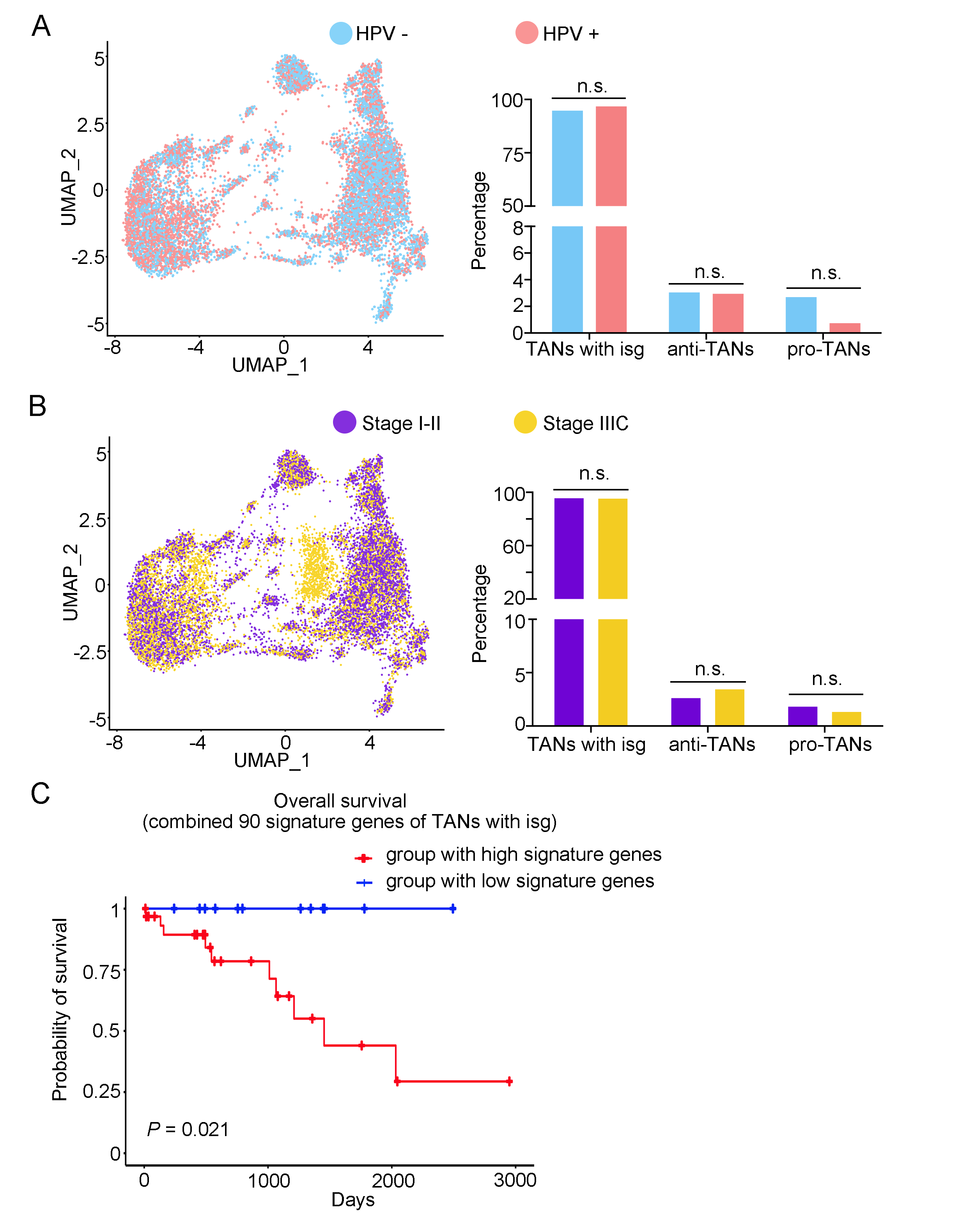

### Supplementary Figure.6

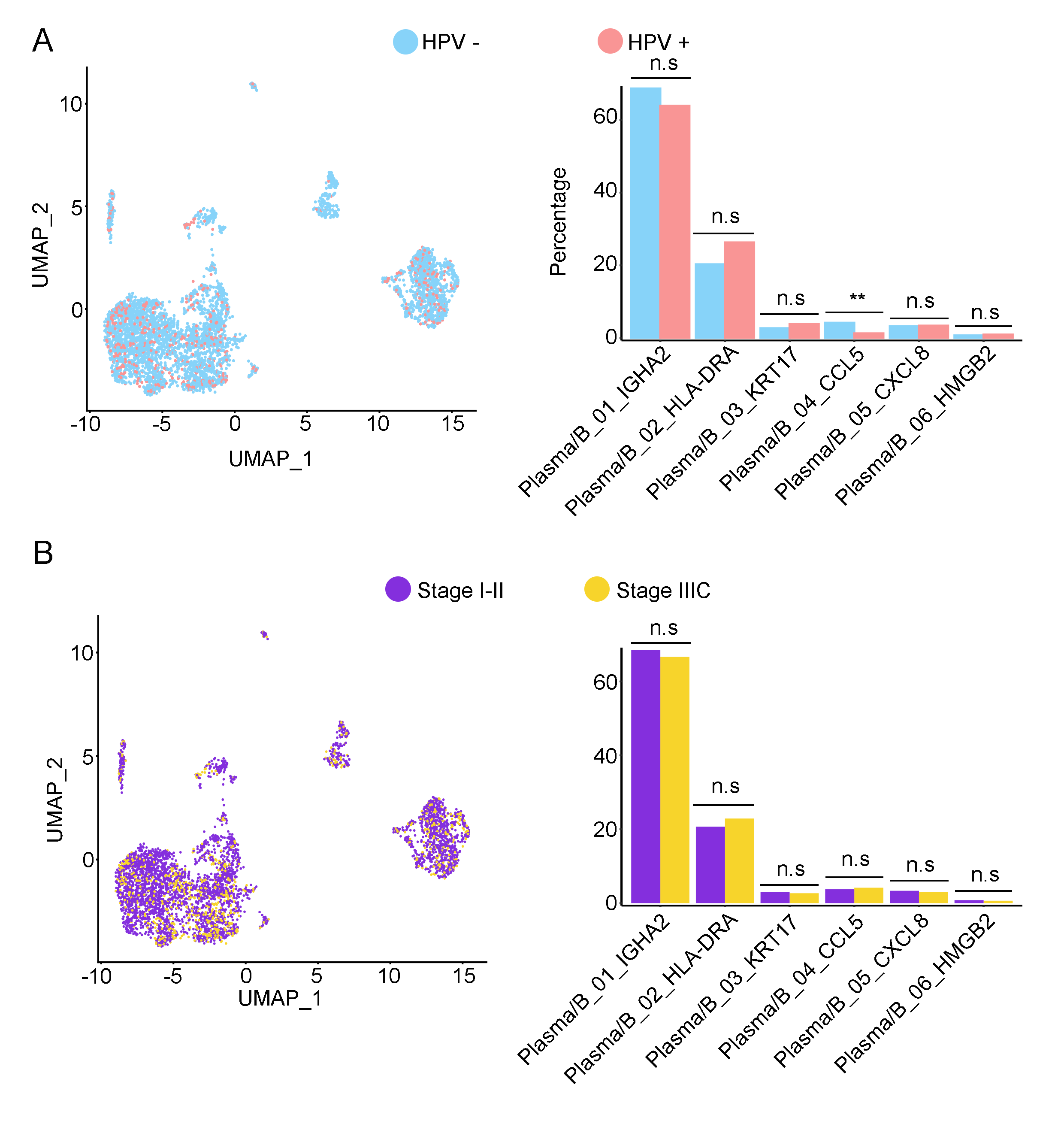

### Supplementary Figure.7

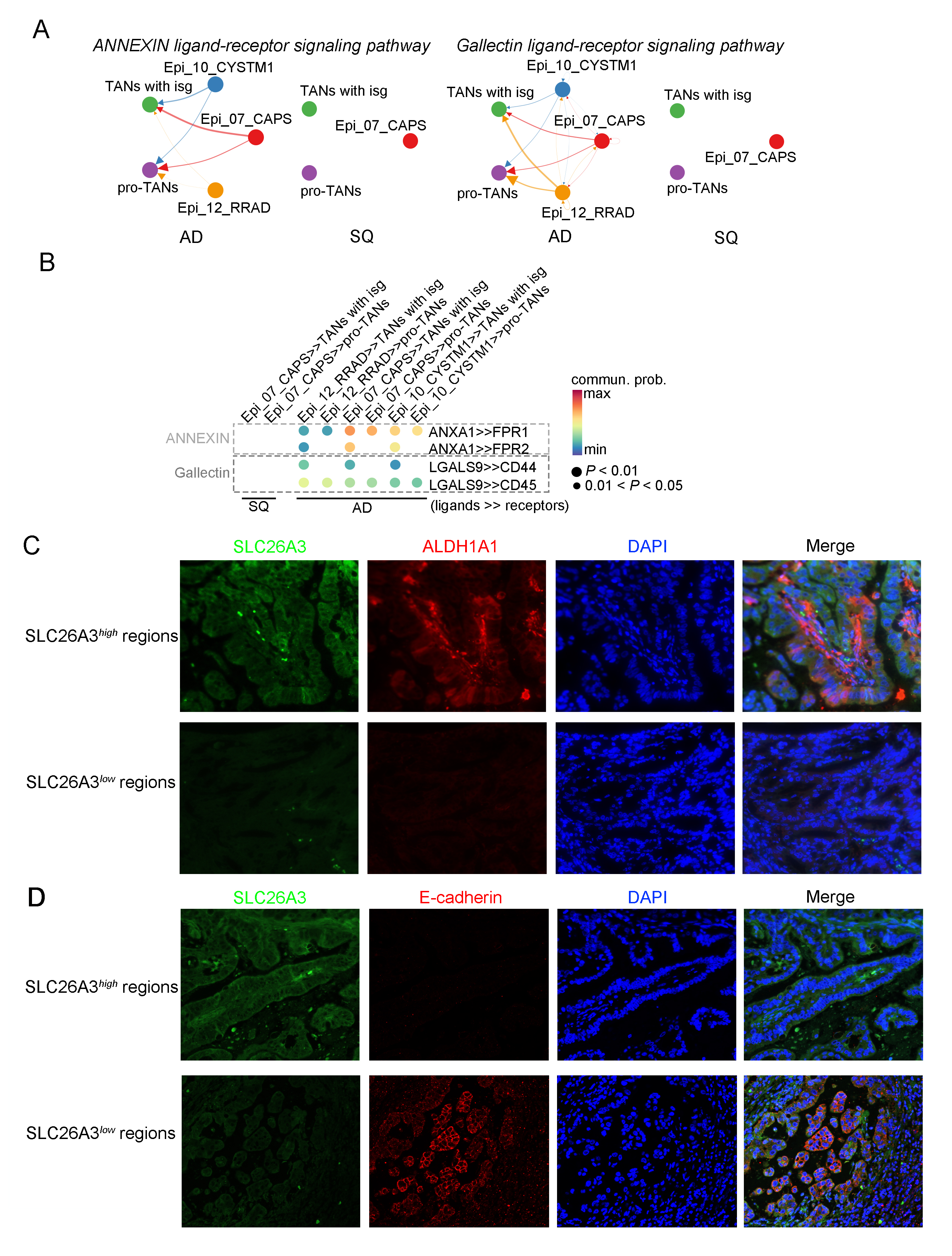

### Supplementary Figure.8

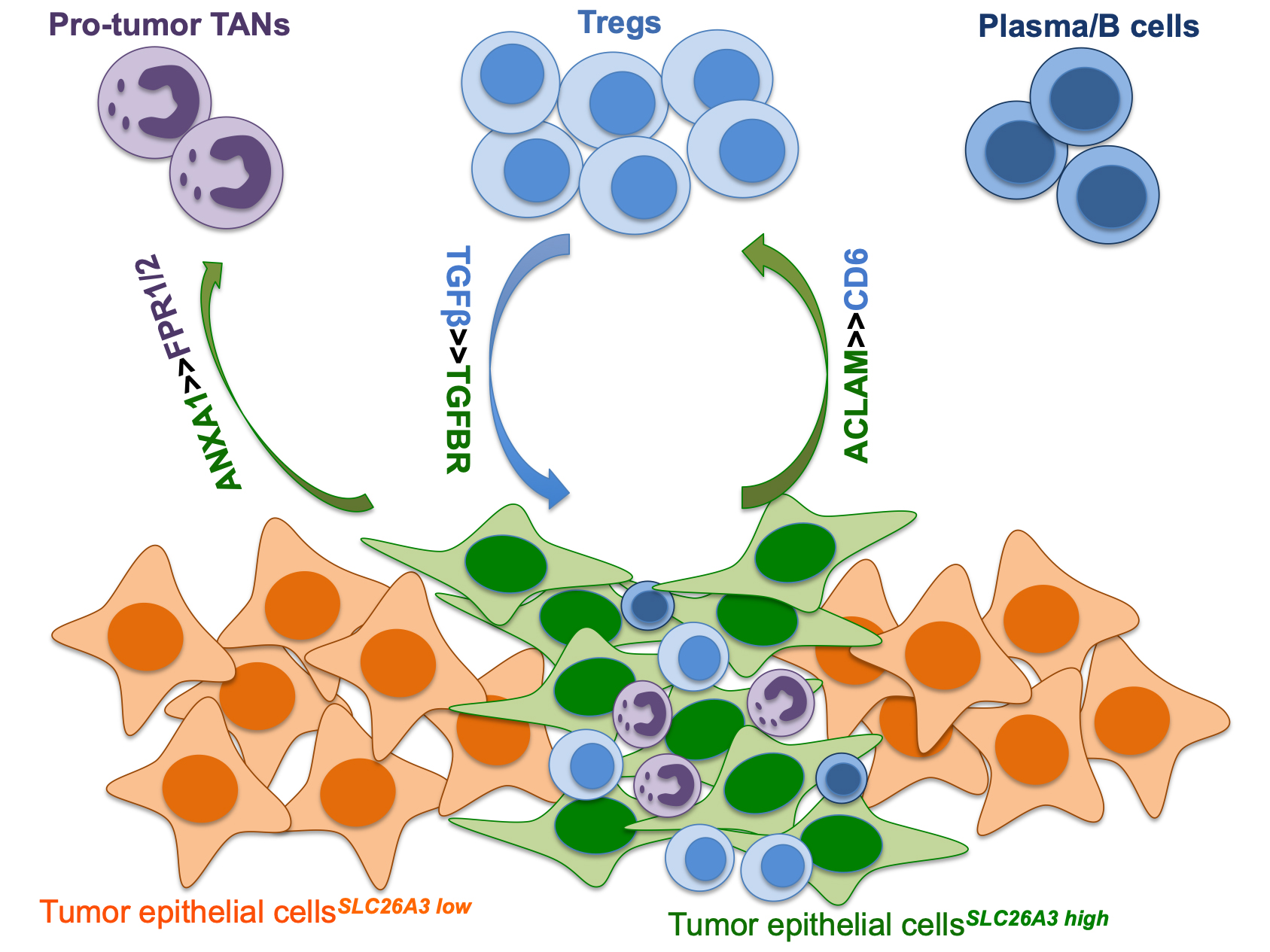

### Supplementary Figure.9

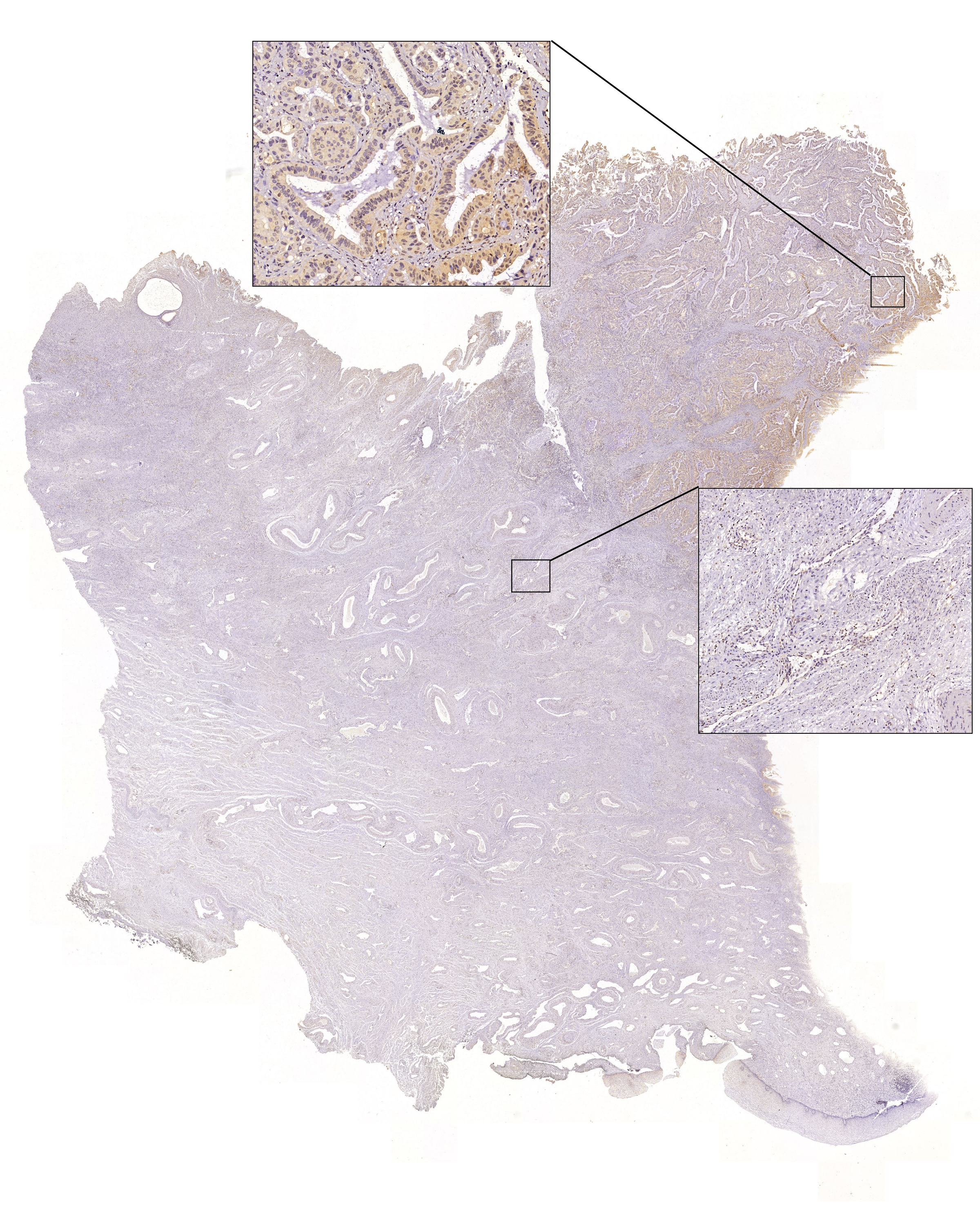
