## Supplementary Figure Legends for "Single-cell profiling reveals the intratumor heterogeneity and immunosuppressive microenvironment in cervical adenocarcinoma"

**Supplementary figure 1. The single-cell genomic atlas of cervical cancer. A.** UMAP and histogram plots to show the differences of distribution (left) and proportion (right) of each cell type between different HPV infection status (HPV+ vs. HPV-). Statistics were performed using R software with two-sided Wilcoxon test (*P* values for each group are listed below: T cell: *P=*0.178; Epithelial cell: *P*=0.017; Neutrophil: *P*=0.792; Macrophage: *P*=0.004; Fibroblast: *P*=1; Plasma: *P*=0.126; Endothelial cell: *P*=0.017; B cell: *P*=0.247; Mast cell: *P*=1). **B.** The UMAP and histogram plots were used to compare the differences of distribution (left) and proportion (right) of each cell type between different stages (stage I-II vs. stage IIIC). Statistics were performed using and R software with two-sided Wilcoxon test. (*P* values for each group are listed below: T cell: *P=*0.514; Epithelial cell: *P*=0.953; Neutrophil: *P*=0.953; Macrophage: *P*=0.953; Fibroblast: *P*=075; Plasma: *P*=0.859; Endothelial cell: *P*=0.440; B cell: *P*=0.953; Mast cell: *P*=0.594). All statistics were shown as **P* < 0.05; ***P* < 0.01；n.s. not significant.

**Supplementary figure 2. The scRNA-seq data reveals the malignant features of tumor epithelial cells. A** and **B.** UMAP and histogram plots to compare the differences of distribution (up) and proportion (down) of each epithelial cell sub-cluster between different HPV infection status (HPV+ vs. HPV-, *P* values for each group are listed below: Epi_01_NTS: *P=*0.017; Epi_02_IGLC2: *P*=0.792; Epi_03_TMRPSS11E: *P*=0.792; Epi_04_TTF2: *P*=0.662; Epi_05_CCL5: *P*=0.247; Epi_06_TMC5: *P*=0.082; Epi_07_CAPS: *P*=0.429; Epi_08_SCGB3A1: *P*=0.009; Epi_09_SST: *P*<0.001; Epi_10_CYSTM1: *P*<0.001; Epi_11_REG1A: *P*<0.001; Epi_12_RRAD: *P*=0.662) and different stages (stage I-II vs. stage IIIC, *P* values for each group are listed below: Epi_01_NTS: *P=*0.953; Epi_02_IGLC2: *P*=0.679; Epi_03_TMRPSS11E: *P*=0.440; Epi_04_TTF2: *P*=0.100; Epi_05_CCL5: *P*=0.371; Epi_06_TMC5: *P*=0.440; Epi_07_CAPS: *P*=0.514; Epi_08_SCGB3A1: *P*=0.440; Epi_09_SST: *P*<0.001; Epi_10_CYSTM1: *P*<0.001; Epi_11_REG1A: *P*<0.001; Epi_12_RRAD: *P*=0.161). **C.** Pseudotime trajectory plots from 4 individual cases to represent the predicted developmental potential of epithelial cells. Statistics were performed using R software with two-sided Wilcoxon test. All statistics were shown as **P* < 0.05; ***P* < 0.01；n.s. not significant.

**Supplementary figure 3. SLC26A3 is identified as a potential prognostic and diagnostic indicator for lymph node metastasis of CC patients. A.** Heatmap showing the top 50 DEGs in cluster Epi_12_RRAD, comparing to all the other sub-clusters. **B.** Violin plots showing the each gene’s expression differences between two groups (top), with UMAP plots (bottom) demonstrating the specificity of each candidate gene, filtering IGF2 as the most identical marker for this sub-cluster. **C.** IHC staining for validation of the protein expression of IGF2 and ADH1C in ADC samples which are classified as early stages (FIGO stage I-IIA) and late stages (FIGO stage IIIC1~2p). **D.** Kaplan-Meier curves showing the overall survival rate of CC patients stratified by the top 50 genes-scaled signature of Epi_12_RRAD. **E.** IHC staining for validation of the protein expression of ORM1/ORM2 and SLC26A3 in ADC samples which are classified as early stages and late stages. As for **C** and **E**, the intensity of each protein marker is shown on 4 individual samples and the scoring is as follows: negative (0), weak (1), intermediate (2), and strong (3). Expression is quantified by the H-score method. **F.** The selecting procedure for stratified comparison (considering HPV status, histological type, *etc*.) in *Cohort 1* (top, in blue) and *Cohort 2* (bottom, in red), respectively. In each comparison module, the case number and rate of post-surgical upstaging are presented.

**Supplementary figure 4. Cellular and molecular heterogeneity of T cells in ADC. A** and **B.** UMAP and histogram plots to show the distribution (left) and proportion (right) of each T cell sub-cluster between different HPV infection status (HPV+ vs. HPV-, *P* values for each group are listed below: Exhausted T: *P=*0.537; Cytotoxic T: *P*=0.662; Treg: *P=*0.329; Activated T: *P=*0.792) and different stages (stage I-II vs. stage IIIC, *P* values for each group are listed below: Exhausted T: *P=*0.768; Cytotoxic T: *P*=0.768; Treg: *P=*0.514; Activated T: *P=*1.000). **C.** Dot heatmap plot of row-scaled expression of overexpressed genes for each sub-cluster of T cells. Statistics were performed using R software with two-sided Wilcoxon test. All statistics were shown as **P* < 0.05; ***P* < 0.01；n.s. not significant.

**Supplementary figure 5. The heterogeneity of tumor associated neutrophils in ADC. A** and **B.** UMAP and histogram plots to compare the differences of distribution (left) and proportion (right) of each sub-type of TANs between different HPV infection status (HPV+ vs. HPV-, *P* values for each group are listed below: TANs with isg: *P=*0.427; anti-TANs: *P*=0.931; pro-TANs: *P*=0.178) and different stages (stage I-II vs. stage IIIC, *P* values for each group are listed below: TANs with isg: *P=*1.000; anti-TANs: *P*=0.514; pro-TANs: *P*=0.951). **C.** Kaplan-Meier curve to show the overall survival rate of CC patients stratified by the top 90 genes-scaled signature of the sub-cluster of TANs with isg. Statistics were performed using R software with two-sided Wilcoxon test. All statistics were shown as **P* < 0.05; ***P* < 0.01；n.s. not significant.

**Supplementary figure 6. Phenotype diversity of plasma/ B cells in ADC. A.** UMAP and histogram plots to show the differences of distribution (left) and proportion (right) of each plasma/B cell sub-cluster between different HPV infection status (HPV+ vs. HPV-, *P* values for each group are listed below: Plasm/B_01_IGHA2: *P=*0.931; Plasm/B_02_HLA-DAR: *P=*0.662; Plasm/B_03_KRT17: *P=*0.931; Plasm/B_04_CCL5: *P=*0.007; Plasm/B_05_CXCL8: *P=*1.000; Plasm/B_06_HMGB2: *P=*0.410). **B.** UMAP and histogram plots to compare the differences of distribution and proportion of each plasma/B cell sub-cluster between different clinical stages (stage I-II vs. stage IIIC, *P* values for each group are listed below: Plasma/B_01_IGHA2: *P=*0.768; Plasma/B_02_HLA-DAR: *P=*0.768; Plasma/B_03_KRT17: *P=*1.000; Plasma/B_04_CCL5: *P=*0.950; Plasma/B_05_CXCL8: *P=*0.667; Plasma/B_06_HMGB2: *P=*0.294). Statistics were performed using R software with two-sided Wilcoxon test. All statistics were shown as **P* < 0.05; ***P* < 0.01；n.s. not significant.

**Supplementary figure 7. The cellular interaction modules of sub-clusters from T cells, neutrophils and tumor epithelial cells. A.** Circle plots showing the interacting networks between epithelial cell sub-clusters and neutrophil sub-clusters via pathways such as ANNEXIN and Gallectin, by comparing ADC and SCC. The direction of one arrow shows the regulation from outputting cells to incoming cells. The width of the each line shows the predicted weight and strength of regulation. **B.** Bubble plot showing the probability of ligand-to-receptor combination of each pathway (corresponding to **A**) between two different sub-types of cells, by comparing ADC with SCC. **C.** Dual IF staining showing that in SLC26A3^high^ regions the stemness of tumor epithelial cells is actively induced than that in SLC26A3^low^ regions. **D.** Dual IF staining showing that in SLC26A3^high^ regions the EMT of tumor epithelial cells is actively induced than in SLC26A3^low^ regions.

**Supplementary figure 8. Graphical presentation of the TIME in ADC, with cellular crosstalk among tumor cells, Tregs, neutrophils and plasma cells.**

**Supplementary figure 9. Full-scan image of the slide from ADC case #5 in Figure 3C.** It is the image of IHC staining via using SLC26A3 antibody to test the expression of this protein on surgically-resected ADC sample #5. The positive (left square) and negative (right square) regions are presented separately.
