## Supplementary Data 1-annotated copies of script for "Single-cell profiling reveals the intratumor heterogeneity and immunosuppressive microenvironment in cervical adenocarcinoma": step1_human_HPV2_harmony.sh

```
#!/bin/sh
#SBATCH -A pi_luyao
#SBATCH -p fatQ
#SBATCH -q fatq
set -exv
date
echo 'initiation of task'
~/software/r-4.1.1/R-4.1.1/bin/Rscript ~/seurat_human_harmony.R -r HPV2_raw_seurat.rds -o HPV2_seurat_deal.rds
echo 'completion of task'
date
```
