## Supplementary Data 1-annotated copies of script for "Single-cell profiling reveals the intratumor heterogeneity and immunosuppressive microenvironment in cervical adenocarcinoma": step1_seurat_human_harmony.R

```
#!/usr/bin/Rscript
suppressMessages(library(optparse))
option_list<-list(
  make_option(c("-r","--rawdata"),type = "character", help  = "Seurat object"),
  make_option(c("-o","--out"),type = "character", help  = "Output the RDS file name")
)
args <- parse_args(OptionParser(option_list=option_list,usage = "Usage: %prog [options] \nDescription: This script is CCA merging the single cell raw matrix data under the current path!"))

message("version: 1.0.0")
message("author: 刘品博-Liu_Pinbo")
message("e-Mail：")
Sys.sleep(0.5)

t1<-Sys.time()
message('*Loading the environment...')
suppressMessages(library("tidyverse"))
suppressMessages(library('ggplot2'))
suppressMessages(library("scales"))
suppressMessages(library("gridExtra"))
options(tibble.width = Inf)
suppressMessages(library("Matrix"))
suppressMessages(library("Seurat"))
suppressMessages(library('harmony'))
suppressMessages(library('cowplot'))
suppressMessages(library('future'))
set.seed(2022)


all_fat_seurat<-readRDS(args$rawdata)

after_CCA<-function(x,t=10,mem=50){
  t_after_CCA<-Sys.time()
  ###质控
  #message('**We are counting mitochondrial gene expression...')
  #x[["percent.mt"]] <- PercentageFeatureSet(x, pattern = "^MT-")
  message('**Filtering in progress...')
  x <- subset(x, subset = nFeature_RNA > 200 & nFeature_RNA < 6000 & percent.mt < 25)
  ###标准化
  message('**Normalization is under way...')
  x <- NormalizeData(x, normalization.method = "LogNormalize", scale.factor = 10000)
  if (t==1) {
    plan("sequential")
  }else{
    plan("multiprocess", workers = t)
    options(future.globals.maxSize = as.numeric(mem)*1000 * 1024^2)
  }
  ###确定高变基因
  x <- FindVariableFeatures(x, selection.method = "vst", nfeatures = 2000)
  ###归一化
  message('**It is being standardized...')
  x <- ScaleData(x, features = rownames(x),vars.to.regress = "percent.mt")
  ###PCA降维
  message('**Pca dimension reduction is in progress')
  plan("sequential")
  x <- RunPCA(x, features = VariableFeatures(object = x),verbose = FALSE)
  ###批次效应处理
  message('**Harmony correction in progress...')
  f<-'ElbowPlot_harmony.pdf'
  while (!is.null(dev.list()))  dev.off()
  pdf(f)
  x <- RunHarmony(x,"orig.ident", plot_convergence = TRUE)
  while (!is.null(dev.list()))  dev.off()
  ###降维+聚类+分簇
  message('**Clustering and visualization in progress...')
  dim.use<-30
  x <- x %>% 
    RunUMAP(reduction = "harmony", dims = 1:dim.use) %>% 
    FindNeighbors(reduction = "harmony", dims = 1:dim.use) %>% 
    FindClusters(resolution = 0.2)
  x <- x %>% 
    RunTSNE(reduction = "harmony", dims = 1:dim.use)
  f<-'tsne_cluster.png'
  while (!is.null(dev.list()))  dev.off()
  png(f,width = 1800,height = 900,res = 90)
  p1<-DimPlot(x, reduction = "tsne", group.by = "orig.ident", pt.size=0.01)
  p2<-DimPlot(x, reduction = "tsne", pt.size=0.01)
  print(plot_grid(p1, p2, labels = c('sample','cluster'),align = "v",ncol = 2))
  while (!is.null(dev.list()))  dev.off()
  f<-'umap_cluster.png'
  while (!is.null(dev.list()))  dev.off()
  png(f,width = 1800,height = 900,res = 90)
  p1<-DimPlot(x, reduction = "umap", group.by = "orig.ident", pt.size=0.01)
  p2<-DimPlot(x, reduction = "umap", pt.size=0.01)
  print(plot_grid(p1, p2, labels = c('sample','cluster'),align = "v",ncol = 2))
  while (!is.null(dev.list()))  dev.off()
  print(Sys.time()-t_after_CCA)
  return(x)
}

x<-after_CCA(all_fat_seurat)
saveRDS(x,args$out)
print(Sys.time()-t1)
```
