## Supplementary Data 1-annotated copies of script for "Single-cell profiling reveals the intratumor heterogeneity and immunosuppressive microenvironment in cervical adenocarcinoma": step2_Fig1_cell_type_ann2.R

```
##col-set====
set.seed(2023)
library(Seurat)
library(future)
library(ggplot2)
library(tidyverse)
##淡黄：#FEBF80; 深红：#DD5856; 深黄：#F38D28；淡红：#F69B9A; 棕色：#B5982F；灰色：#7D6D6A；草绿：#8DCA7D；
#淡蓝：#9FCCE6；深蓝：#4E7CA6
Science_col<-unique(c("#9FCCE6","#DD5856","#F38D28","#F69B9A","#B5982F","#FEBF80","#DD5856","#F38D28","#F69B9A","#B5982F","#7D6D6A","#8DCA7D","#9FCCE6","#4E7CA6","#D57096",
                      "#AD7AA5","#FDBFD2","#DBA3C8","#CC5C15","#D42F7E",'#E5C06A','#E28D8F',"#80B880","#EDB1C4","#5378A1","#847C7D","#4E928D","#E7863C","#B9AAAD",
                      "#D45760","#F1AD7C","#66639E","#92B646","#157BB7","#EDBD41","#229873","#CC5C15","#D42F7E","#DBA513","#638CC9","#6F6AA7","#A1C7DC","#359837","#ED7A1C","#8A298C",
                      "#286EA1","#DEEDD0","#F49897",'#61B5BA','#D96951','#40507D','#9D5B34','#5D3C8A','#997635','#729D44',
                      '#666666','#C2362B','#EABB7B','#C1AFCE'))


#大群MarkerVlnplot====
seurat_deal<-readRDS("E:/work_station/2023/3_6_HPV2/01_data/HPV2_seurat_deal.rds")
out<-"E:/work_station/2023/3_6_HPV2/02_all"
Group_1<-'AD'
Group_2<-'SQ'
Group_3<-NULL
Group_4<-NULL

# #降维图
# while (!is.null(dev.list()))  dev.off()
# png(filename = paste0(out,"/","all_umap.png"),width = 1100,height = 900)
# print(DimPlot(seurat_deal,label = TRUE,label.size = 10,pt.size = 0.8,raster=FALSE,cols = Science_col)+ggtitle('all'))
# while (!is.null(dev.list()))  dev.off()
# pdf(file = paste0(out,"/","all_umap.pdf"),width = 11,height = 9)
# print(DimPlot(seurat_deal,label = TRUE,label.size = 10,pt.size = 0.8,raster=FALSE,cols = Science_col)+ggtitle('all'))
# while (!is.null(dev.list()))  dev.off()
# while (!is.null(dev.list()))  dev.off()
# png(filename = paste0(out,"/","all_tsne.png"),width = 1100,height = 900)
# print(DimPlot(seurat_deal,reduction = "tsne",label = TRUE,label.size = 10,pt.size = 0.8,raster=FALSE,cols = Science_col)+ggtitle('all'))
# while (!is.null(dev.list()))  dev.off()
# pdf(file = paste0(out,"/","all_tsne.pdf"),width = 11,height = 9)
# print(DimPlot(seurat_deal,reduction = "tsne",label = TRUE,label.size = 10,pt.size = 0.8,raster=FALSE,cols = Science_col)+ggtitle('all'))
# while (!is.null(dev.list()))  dev.off()
# 
# 
# human_common_marker_list<-list(immune=c("PTPRC"),
#                                Tcell=c("CD3E","CD3D","CD3G"),
#                                Bcell=c("CD19"),
#                                Endotheliocyte=c('VWF',"PECAM1"),
#                                Mastcell=c('KIT','FCER1A'),
#                                Neutrophil=c('S100A8', 'S100A9'),
#                                macrophage=c('ADGRE1','CCR5','CD63'),
#                                Monocyte=c('CD14','CD16'),
#                                Plasma=c('JCHAIN','XBP1'),
#                                Fibroblast=c('COL3A1','DCN','COL1A1'))
# if (!(dir.exists(paste(sep = "/", out, "common_marker")))) {
#   dir.create(paste(sep = "/", out, "common_marker"))
# }
# library(homologene)
# for (i in 1:length(human_common_marker_list)) {
#   P<-VlnPlot(seurat_deal, features = human_common_marker_list[[i]], pt.size = 0)
#   save_picture(P, out = paste(sep = "/",out,"common_marker"), Pname = names(human_common_marker_list)[i], width_len = 1100, height_len = 900)
# }


#大群注释====
seurat_deal<-readRDS("E:/work_station/2023/3_6_HPV2/01_data/HPV2_seurat_deal.rds")
out<-"E:/work_station/2023/3_6_HPV2/03_ann"
Group_1<-'AD'
Group_2<-'SQ'
Group_3<-NULL
Group_4<-NULL

# seurat_ann<-ann_and_plot(seurat_deal, out,
#                          ann_file = 'E:/work_station/2023/3_6_HPV2/02_all/HPV2_ann.txt',
#                          meta_names = "cell_type", label_size = 10, pt.size = 1)
# 
# 


ann<-read.table(file = "E:/work_station/2023/3_6_HPV2/02_all/HPV2_ann.txt",
                header = F,check.names = F)
if(all.equal(as.character(levels(seurat_deal)),as.character(ann$V1)) == "TRUE"){
  new_id<-ann$V2
  names(new_id)<-levels(seurat_deal)
  new_id
  seurat_deal <- RenameIdents(seurat_deal,new_id)
  all.equal(rownames(seurat_deal@meta.data),names(seurat_deal@active.ident))
  meta_names = "cell_type"
  seurat_deal@meta.data[,meta_names]<-as.factor(seurat_deal@active.ident)
  levels(seurat_deal)
}
```
