## Supplementary Data 1-annotated copies of script for "Single-cell profiling reveals the intratumor heterogeneity and immunosuppressive microenvironment in cervical adenocarcinoma": EPI_cluster_ann.R

```
###上皮细胞分群注释

#大群注释====
seurat_deal<-readRDS("E:/work_station/2023/3_6_HPV2/01_data/HPV2_seurat_deal.rds")
out<-"E:/work_station/2023/3_6_HPV2/03_ann"
Group_1<-'AD'
Group_2<-'SQ'
Group_3<-NULL
Group_4<-NULL

seurat_ann<-ann_and_plot(seurat_deal, out,
                         ann_file = 'E:/work_station/2023/3_6_HPV2/02_all/HPV2_ann.txt',
                         meta_names = "cell_type", label_size = 10, pt.size = 1)

#Epi分群====
#human_after_CCA:亚群标准分析
human_after_CCA<-function(x,resolution,t=1,mem=1){
  suppressMessages(library("tidyverse"))
  suppressMessages(library('ggplot2'))
  suppressMessages(library("scales"))
  suppressMessages(library("gridExtra"))
  suppressMessages(library("Matrix"))
  suppressMessages(library("Seurat"))
  suppressMessages(library('harmony'))
  suppressMessages(library('cowplot'))
  suppressMessages(library('future'))
  if (!(exists("resolution"))) {
    stop("Error: resolution is null")
  }
  t_after_CCA<-Sys.time()
  set.seed(2022)
  message("**Statistical Period information")
  s.genes <- read.table(file = "~/细胞周期标志物S期.txt",sep="\t",header=T)
  g2m.genes <- read.table(file = "~/细胞周期标志物G2M期.txt",sep="\t",header=T)
  message('物种：human')
  g2m.genes<-g2m.genes$G2M_gene
  s.genes<-s.genes$S_gene
  x<- CellCycleScoring(object = x, g2m.features = g2m.genes, s.features = s.genes)
  message('**Normalization is under way...')
  x <- NormalizeData(x, normalization.method = "LogNormalize", scale.factor = 10000)
  x <- FindVariableFeatures(x, selection.method = "vst", nfeatures = 2000)
  message('**It is being standardized...')
  library(future)
  if (t==1) {
    plan("sequential")
  }else{
    plan("multiprocess", workers = t)
    options(future.globals.maxSize = as.numeric(mem)*1000 * 1024^2)
  }
  x <- ScaleData(x, features = rownames(x),vars.to.regress = c("percent.mt","S.Score","G2M.Score"))
  message('**Pca dimension reduction is in progress')
  x <- RunPCA(x, features = VariableFeatures(object = x),verbose = FALSE)
  message('**Harmony correction in progress...')
  x <- RunHarmony(x,"orig.ident", plot_convergence = TRUE)
  message('**Clustering and visualization in progress...')
  dim.use<-30
  #####降维+聚类+分群
  x <- x %>% 
    RunUMAP(reduction = "harmony", dims = 1:dim.use) %>% 
    FindNeighbors(reduction = "harmony", dims = 1:dim.use) %>% 
    FindClusters(resolution = resolution)
  x <- x %>% 
    RunTSNE(reduction = "harmony", dims = 1:dim.use)
  
  plan("sequential")
  print(Sys.time()-t_after_CCA)
  return(x)
}
temp_subseurat<-subset(seurat_ann, idents = "Epithelial_cell")
temp_subseurat_deal<-human_after_CCA(temp_subseurat, resolution = 0.1, t = 4, mem = 30)


###EPI注释：先进行差异分析，将分群的差异基因按照对应的avg_log2FC升序排序，取排序第一的差异基因作为该群的名字
temp_DE<-FindAllMarkers(temp_subseurat_deal,logfc.threshold = 0,only.pos = T) ###差异分析

new_id<-1:length(levels(temp_subseurat_deal))
names(new_id)<-levels(temp_subseurat_deal)
for (i in 1:length(levels(temp_subseurat_deal))) {
  new_id[i]<-paste(subset(temp_DE, cluster==names(new_id)[i]) %>% arrange(-avg_log2FC) %>% .[1,"gene"],
                   names(subgroup)[[groupone]],sep = "+_")
}

new_id
temp_subseurat_deal <- RenameIdents(temp_subseurat_deal,new_id)
meta_names = "sub_celltype"
temp_subseurat_deal@meta.data[,meta_names]<-as.factor(temp_subseurat_deal@active.ident)
```
