## Supplementary Data 1-annotated copies of script for "Single-cell profiling reveals the intratumor heterogeneity and immunosuppressive microenvironment in cervical adenocarcinoma": step2_σÅîτ╗åΦâ₧σÄ╗ΘÖñ.R

```
setwd("E:/work_station/2023/3_6_HPV2/20240905_supplement/Tcell")
library(DoubletFinder)
library(tidyverse)
library(Seurat)
library(patchwork)
rm(list=ls())

seurat_data<-readRDS("E:/work_station/2023/3_6_HPV2/05_subset/RDS/T_cell_ann.rds")
levels(seurat_data)

###双细胞去除
ls("package:DoubletFinder")
# library(rstatix)
# trace(paramSweep,edit = T)##修改：seu@assays$RNA$counts改为seu@assays$RNA@counts
paramSweep_v3<-function (seu, PCs = 1:10, sct = FALSE, num.cores = 1) 
{
  require(Seurat)
  require(fields)
  require(parallel)
  pK <- c(5e-04, 0.001, 0.005, seq(0.01, 0.3, by = 0.01))
  pN <- seq(0.05, 0.3, by = 0.05)
  min.cells <- round(nrow(seu@meta.data)/(1 - 0.05) - nrow(seu@meta.data))
  pK.test <- round(pK * min.cells)
  pK <- pK[which(pK.test >= 1)]
  orig.commands <- seu@commands
  if (nrow(seu@meta.data) > 10000) {
    real.cells <- rownames(seu@meta.data)[sample(1:nrow(seu@meta.data), 
                                                 10000, replace = FALSE)]
    data <- seu@assays$RNA@counts[, real.cells]
    n.real.cells <- ncol(data)
  }
  if (nrow(seu@meta.data) <= 10000) {
    real.cells <- rownames(seu@meta.data)
    data <- seu@assays$RNA@counts
    n.real.cells <- ncol(data)
  }
  if (num.cores > 1) {
    require(parallel)
    cl <- makeCluster(num.cores)
    output2 <- mclapply(as.list(1:length(pN)), FUN = parallel_paramSweep, 
                        n.real.cells, real.cells, pK, pN, data, orig.commands, 
                        PCs, sct, mc.cores = num.cores)
    stopCluster(cl)
  }
  else {
    output2 <- lapply(as.list(1:length(pN)), FUN = parallel_paramSweep, 
                      n.real.cells, real.cells, pK, pN, data, orig.commands, 
                      PCs, sct)
  }
  sweep.res.list <- list()
  list.ind <- 0
  for (i in 1:length(output2)) {
    for (j in 1:length(output2[[i]])) {
      list.ind <- list.ind + 1
      sweep.res.list[[list.ind]] <- output2[[i]][[j]]
    }
  }
  name.vec <- NULL
  for (j in 1:length(pN)) {
    name.vec <- c(name.vec, paste("pN", pN[j], "pK", pK, 
                                  sep = "_"))
  }
  names(sweep.res.list) <- name.vec
  return(sweep.res.list)
}


## (1)pK Identification： 确定参数，寻找最优pK值 ----------------------------------------------------------
#这是一个测试最佳参数的过程，运行速度慢
sweep.res.list_keloid <- paramSweep_v3(seurat_data, PCs = 1:20, sct = FALSE)
#head(sweep.res.list_keloid)
sweep.stats_keloid <- summarizeSweep(sweep.res.list_keloid, GT = FALSE)
bcmvn_keloid <- find.pK(sweep.stats_keloid) #可以看到最佳参数的点
## 所以最佳的参数是：
mpK<-as.numeric(as.vector(bcmvn_keloid$pK[which.max(bcmvn_keloid$BCmetric)]))


## (2) Homotypic Doublet Proportion Estimate:双细胞比例计算 -------------------------------------
annotations <- seurat_data@meta.data$seurat_clusters
homotypic.prop <- modelHomotypic(annotations)  
DoubletRate = ncol(seurat_data)*8*1e-6 #按每增加1000个细胞，双细胞比率增加千分之8来计算
DoubletRate = 0.309016
#估计同源双细胞比例，根据modelHomotypic()中的参数人为混合双细胞。这里是从seurat_clusters中来混双细胞 
nExp_poi <- round(DoubletRate*length(seurat_data$seurat_clusters))  #最好提供celltype，而不是seurat_clusters。
# 计算双细胞比例
nExp_poi.adj <- round(nExp_poi*(1-homotypic.prop))


## (3)最后，使用确定好的参数鉴定Doublets. Run DoubletFinder with varying classification stringencies ----------------------------------------------------------------
doubletFinder_v3<-function (seu, PCs, pN = 0.25, pK, nExp, reuse.pANN = FALSE, 
                            sct = FALSE, annotations = NULL) 
{
  require(Seurat)
  require(fields)
  require(KernSmooth)
  if (reuse.pANN != FALSE) {
    pANN.old <- seu@meta.data[, reuse.pANN]
    classifications <- rep("Singlet", length(pANN.old))
    classifications[order(pANN.old, decreasing = TRUE)[1:nExp]] <- "Doublet"
    seu@meta.data[, paste("DF.classifications", pN, pK, 
                          nExp, sep = "_")] <- classifications
    return(seu)
  }
  if (reuse.pANN == FALSE) {
    real.cells <- rownames(seu@meta.data)
    data <- seu@assays$RNA@counts[, real.cells]
    n_real.cells <- length(real.cells)
    n_doublets <- round(n_real.cells/(1 - pN) - n_real.cells)
    print(paste("Creating", n_doublets, "artificial doublets...", 
                sep = " "))
    real.cells1 <- sample(real.cells, n_doublets, replace = TRUE)
    real.cells2 <- sample(real.cells, n_doublets, replace = TRUE)
    doublets <- (data[, real.cells1] + data[, real.cells2])/2
    colnames(doublets) <- paste("X", 1:n_doublets, sep = "")
    data_wdoublets <- cbind(data, doublets)
    if (!is.null(annotations)) {
      stopifnot(typeof(annotations) == "character")
      stopifnot(length(annotations) == length(Cells(seu)))
      stopifnot(!any(is.na(annotations)))
      annotations <- factor(annotations)
      names(annotations) <- Cells(seu)
      doublet_types1 <- annotations[real.cells1]
      doublet_types2 <- annotations[real.cells2]
    }
    orig.commands <- seu@commands
    if (sct == FALSE) {
      print("Creating Seurat object...")
      seu_wdoublets <- CreateSeuratObject(counts = data_wdoublets)
      print("Normalizing Seurat object...")
      seu_wdoublets <- NormalizeData(seu_wdoublets, normalization.method = orig.commands$NormalizeData.RNA@params$normalization.method, 
                                     scale.factor = orig.commands$NormalizeData.RNA@params$scale.factor, 
                                     margin = orig.commands$NormalizeData.RNA@params$margin)
      print("Finding variable genes...")
      seu_wdoublets <- FindVariableFeatures(seu_wdoublets, 
                                            selection.method = orig.commands$FindVariableFeatures.RNA$selection.method, 
                                            loess.span = orig.commands$FindVariableFeatures.RNA$loess.span, 
                                            clip.max = orig.commands$FindVariableFeatures.RNA$clip.max, 
                                            mean.function = orig.commands$FindVariableFeatures.RNA$mean.function, 
                                            dispersion.function = orig.commands$FindVariableFeatures.RNA$dispersion.function, 
                                            num.bin = orig.commands$FindVariableFeatures.RNA$num.bin, 
                                            binning.method = orig.commands$FindVariableFeatures.RNA$binning.method, 
                                            nfeatures = orig.commands$FindVariableFeatures.RNA$nfeatures, 
                                            mean.cutoff = orig.commands$FindVariableFeatures.RNA$mean.cutoff, 
                                            dispersion.cutoff = orig.commands$FindVariableFeatures.RNA$dispersion.cutoff)
      print("Scaling data...")
      seu_wdoublets <- ScaleData(seu_wdoublets, features = orig.commands$ScaleData.RNA$features, 
                                 model.use = orig.commands$ScaleData.RNA$model.use, 
                                 do.scale = orig.commands$ScaleData.RNA$do.scale, 
                                 do.center = orig.commands$ScaleData.RNA$do.center, 
                                 scale.max = orig.commands$ScaleData.RNA$scale.max, 
                                 block.size = orig.commands$ScaleData.RNA$block.size, 
                                 min.cells.to.block = orig.commands$ScaleData.RNA$min.cells.to.block)
      print("Running PCA...")
      seu_wdoublets <- RunPCA(seu_wdoublets, features = orig.commands$ScaleData.RNA$features, 
                              npcs = length(PCs), rev.pca = orig.commands$RunPCA.RNA$rev.pca, 
                              weight.by.var = orig.commands$RunPCA.RNA$weight.by.var, 
                              verbose = FALSE)
      pca.coord <- seu_wdoublets@reductions$pca@cell.embeddings[, 
                                                                PCs]
      cell.names <- rownames(seu_wdoublets@meta.data)
      nCells <- length(cell.names)
      rm(seu_wdoublets)
      gc()
    }
    if (sct == TRUE) {
      require(sctransform)
      print("Creating Seurat object...")
      seu_wdoublets <- CreateSeuratObject(counts = data_wdoublets)
      print("Running SCTransform...")
      seu_wdoublets <- SCTransform(seu_wdoublets)
      print("Running PCA...")
      seu_wdoublets <- RunPCA(seu_wdoublets, npcs = length(PCs))
      pca.coord <- seu_wdoublets@reductions$pca@cell.embeddings[, 
                                                                PCs]
      cell.names <- rownames(seu_wdoublets@meta.data)
      nCells <- length(cell.names)
      rm(seu_wdoublets)
      gc()
    }
    print("Calculating PC distance matrix...")
    dist.mat <- fields::rdist(pca.coord)
    print("Computing pANN...")
    pANN <- as.data.frame(matrix(0L, nrow = n_real.cells, 
                                 ncol = 1))
    if (!is.null(annotations)) {
      neighbor_types <- as.data.frame(matrix(0L, nrow = n_real.cells, 
                                             ncol = length(levels(doublet_types1))))
    }
    rownames(pANN) <- real.cells
    colnames(pANN) <- "pANN"
    k <- round(nCells * pK)
    for (i in 1:n_real.cells) {
      neighbors <- order(dist.mat[, i])
      neighbors <- neighbors[2:(k + 1)]
      pANN$pANN[i] <- length(which(neighbors > n_real.cells))/k
      if (!is.null(annotations)) {
        for (ct in unique(annotations)) {
          neighbors_that_are_doublets = neighbors[neighbors > 
                                                    n_real.cells]
          if (length(neighbors_that_are_doublets) > 
              0) {
            neighbor_types[i, ] <- table(doublet_types1[neighbors_that_are_doublets - 
                                                          n_real.cells]) + table(doublet_types2[neighbors_that_are_doublets - 
                                                                                                  n_real.cells])
            neighbor_types[i, ] <- neighbor_types[i, 
            ]/sum(neighbor_types[i, ])
          }
          else {
            neighbor_types[i, ] <- NA
          }
        }
      }
    }
    print("Classifying doublets..")
    classifications <- rep("Singlet", n_real.cells)
    classifications[order(pANN$pANN[1:n_real.cells], decreasing = TRUE)[1:nExp]] <- "Doublet"
    seu@meta.data[, paste("pANN", pN, pK, nExp, sep = "_")] <- pANN[rownames(seu@meta.data), 
                                                                    1]
    seu@meta.data[, paste("DF.classifications", pN, pK, 
                          nExp, sep = "_")] <- classifications
    if (!is.null(annotations)) {
      colnames(neighbor_types) = levels(doublet_types1)
      for (ct in levels(doublet_types1)) {
        seu@meta.data[, paste("DF.doublet.contributors", 
                              pN, pK, nExp, ct, sep = "_")] <- neighbor_types[, 
                                                                              ct]
      }
    }
    return(seu)
  }
}

# seurat_data <- doubletFinder_v3(seurat_data, PCs = 1:10, pN = 0.25, pK = mpK, 
#                                 nExp = nExp_poi, reuse.pANN = FALSE, sct = F)
seurat_data <- doubletFinder_v3(seurat_data, PCs = 1:10, pN = 0.25, pK = mpK, 
                                nExp = nExp_poi.adj, reuse.pANN = FALSE, sct = F)
# 使用nExp = nExp_poi和nExp = nExp_poi.adj,分别进行doublets鉴定，以便后续确定哪些细胞是Doublet-High Confidience


## Plot results ---------------------------------------------------------------------------

# seurat_data@meta.data[,"DF_hi.lo"] <- seurat_data@meta.data$DF.classifications_0.25_0.005_9664
# seurat_data@meta.data$DF_hi.lo[which(seurat_data@meta.data$DF_hi.lo == "Doublet" & seurat_data@meta.data$DF.classifications_0.25_0.01_198 == "Singlet")] <- "Doublet-Low Confidience"
# seurat_data@meta.data$DF_hi.lo[which(seurat_data@meta.data$DF_hi.lo == "Doublet")] <- "Doublet-High Confidience"
# table(seurat_data@meta.data$DF_hi.lo)
# Doublet-High Confidience  Doublet-Low Confidience                  Singlet 
# 198                       24                     5041 

## 结果展示，分类结果在pbmc@meta.data中
# png("./output/2_cell_annotation/2_doubletFinder.png",2500,1800,res=300)
# DimPlot(seurat_data, reduction = "umap", group.by ="DF_hi.lo",cols =c("black","red","gold"))
# dev.off()

png("./doubletFinder.png",2000,1800,res=300)
DimPlot(seurat_data, reduction = "umap", group.by ="DF.classifications_0.25_0.005_9664")
dev.off()

# seurat_deal<-subset(seurat_data,DF.classifications_0.25_0.005_9664 == "Singlet")
# DimPlot(seurat_deal, reduction = "umap", group.by ="DF.classifications_0.25_0.005_9664")
saveRDS(seurat_data,file = "E:/work_station/2023/3_6_HPV2/20240905_supplement/Tcell/RDS/Tcell_seurat.rds")
```
