## Supplementary Data 1-annotated copies of script for "Single-cell profiling reveals the intratumor heterogeneity and immunosuppressive microenvironment in cervical adenocarcinoma": step3_Tcell_ann.R

```
setwd("E:/work_station/2023/3_6_HPV2/20240905_supplement/Tcell")
rm(list = ls())
suppressMessages(library("Seurat"))
suppressMessages(library('ggplot2'))
suppressMessages(library("RColorBrewer"))
suppressMessages(library('xlsx'))
suppressMessages(library("ggsci"))


output_dir3<-"20240905_ann"
species<-"human"
#human_common_marker_data<-"G:/human_commom_marker.xlsx"
# groups<-"group.txt"
seurat_data<-readRDS("E:/work_station/2023/3_6_HPV2/20240905_supplement/Tcell/RDS/Tcell_seurat.rds")
seurat_data<-subset(seurat_data,DF.classifications_0.25_0.005_9664 == "Singlet")
levels(seurat_data)


if(dir.exists(output_dir3) == FALSE){
  dir.create(output_dir3)
}

#################################################注释
out_result<-paste0(output_dir3,"/result")
if(!dir.exists(out_result)){
  dir.create(out_result,recursive = T)
}

seurat_data<-subset(seurat_data,`RNA_snn_res.0.1` %in% as.character(c(0,1,2,3,4,5,6)))
levels(seurat_data)

ann<-read.table(file = "20240905_ann/ann/Tcell_ann_final.txt",
                header = F,check.names = F)

new_id<-ann$V2
names(new_id)<-levels(seurat_data)
new_id
seurat_data <- RenameIdents(seurat_data,new_id)
all.equal(rownames(seurat_data@meta.data),names(seurat_data@active.ident))
meta_names = "subcell_type2"
seurat_data@meta.data[,meta_names]<-as.factor(seurat_data@active.ident)
levels(seurat_data)

levels_cell_type<-c("Exhausted_T","Cytotoxic_T","Treg","Activated_T")
seurat_data$subcell_type<-factor(seurat_data$subcell_type2,levels = rev(levels_cell_type))
Idents(object = seurat_data) <- "subcell_type2"
levels(seurat_data)
```
