## Supplementary Data 1-annotated copies of script for "Single-cell profiling reveals the intratumor heterogeneity and immunosuppressive microenvironment in cervical adenocarcinoma": Tcell_ann_plot.R

```
setwd("E:/work_station/2023/3_6_HPV2/20240905_supplement/Tcell")
rm(list = ls())
suppressMessages(library("Seurat"))
suppressMessages(library('ggplot2'))
suppressMessages(library("RColorBrewer"))
suppressMessages(library('xlsx'))
suppressMessages(library("ggsci"))


output_dir3<-"20240905_ann"
species<-"human"
#human_common_marker_data<-"G:/human_commom_marker.xlsx"
# groups<-"group.txt"
seurat_data<-readRDS("RDS/Tcell_seurat.rds")
seurat_data<-subset(seurat_data,DF.classifications_0.25_0.005_9664 == "Singlet")
levels(seurat_data)


if(dir.exists(output_dir3) == FALSE){
  dir.create(output_dir3)
}

## 图片R对象存储
message('save_picture')
save_picture<-function(picture, out, Pname, width_len = 1100, height_len = 900){
  library(ggplot2)
  while (!is.null(dev.list()))  dev.off()
  png(filename = paste(sep = "/", out, paste0(Pname,".png")),width = width_len, height = height_len)
  print(picture)
  while (!is.null(dev.list()))  dev.off()
  while (!is.null(dev.list()))  dev.off()
  pdf(file = paste(sep = "/", out, paste0(Pname,".pdf")),width = as.numeric(width_len/100), height = as.numeric(height_len/100))
  print(picture)
  while (!is.null(dev.list()))  dev.off()
}


Science_col<-unique(c("#9FCCE6","#DD5856","#F38D28","#F69B9A","#B5982F","#FEBF80",
                      "#DD5856","#F38D28","#F69B9A","#B5982F","#7D6D6A","#8DCA7D",
                      "#9FCCE6","#4E7CA6","#D57096","#AD7AA5","#FDBFD2","#DBA3C8",
                      "#CC5C15","#D42F7E",'#E5C06A','#E28D8F',"#80B880","#EDB1C4",
                      "#5378A1","#847C7D","#4E928D","#E7863C","#B9AAAD","#D45760",
                      "#F1AD7C","#66639E","#92B646","#157BB7","#EDBD41","#229873",
                      "#CC5C15","#D42F7E","#DBA513","#638CC9","#6F6AA7","#A1C7DC",
                      "#359837","#ED7A1C","#8A298C","#286EA1","#DEEDD0","#F49897",
                      '#61B5BA','#D96951','#40507D','#9D5B34','#5D3C8A','#997635',
                      '#729D44','#666666','#C2362B','#EABB7B','#C1AFCE'))


#################################################注释
out_result<-paste0(output_dir3,"/result")
if(!dir.exists(out_result)){
  dir.create(out_result,recursive = T)
}


###A==========================
seurat_data<-SetIdent(seurat_data,value = "orig.ident")
levels(seurat_data)
p<-DimPlot(seurat_data,reduction = "umap",pt.size = 1)
p
save_picture(p, paste(out_result,"A",sep = "/") , Pname = "1_Tcell_sample_all_umap",width_len = 1000, height_len = 1000)


seurat_data<-SetIdent(seurat_data,value = "RNA_snn_res.0.1")
levels(seurat_data)
p<-DimPlot(seurat_data,reduction = "umap",pt.size = 1,cols = Science_col,label = T)
p
save_picture(p, paste(out_result,"A",sep = "/") , Pname = "2_Tcell_cluster_all_umap",width_len = 1000, height_len = 1000)


seurat_data<-subset(seurat_data,`RNA_snn_res.0.1` %in% as.character(c(0,1,2,3,4,5,6)))
levels(seurat_data)

ann<-read.table(file = "20240905_ann/ann/Tcell_ann_final.txt",
                header = F,check.names = F)
if(all.equal(as.character(levels(seurat_data)),as.character(ann$V1)) == "TRUE"){
  new_id<-ann$V2
  names(new_id)<-levels(seurat_data)
  new_id
  seurat_data <- RenameIdents(seurat_data,new_id)
  all.equal(rownames(seurat_data@meta.data),names(seurat_data@active.ident))
  meta_names = "subcell_type2"
  seurat_data@meta.data[,meta_names]<-as.factor(seurat_data@active.ident)
  levels(seurat_data)
}
levels_cell_type<-c("Exhausted_T","Cytotoxic_T","Treg","Activated_T")
# rev(levels_cell_type)
seurat_data$subcell_type<-factor(seurat_data$subcell_type2,levels = rev(levels_cell_type))
Idents(object = seurat_data) <- "subcell_type2"
levels(seurat_data)

p<-DimPlot(seurat_data,reduction = "umap",pt.size = 1,cols = c("DodgerBlue","red","yellow","green"))
p
save_picture(p, paste(out_result,"A",sep = "/") , Pname = "3_Tcell_ann_umap",width_len = 1000, height_len = 1000)

###B==========================================
ann_marker_data<-"20240905_ann/marker/Tcell_marker_final.xlsx"
file.copy(ann_marker_data,out_result)
ann_marker_list <- lapply(as.data.frame(read.xlsx(ann_marker_data,sheetIndex=1)), function(x) x[complete.cases(x)])
ann_gene_list<-list()
ann_markers_all<-c()
library("stringr")
for (i in 1:length(ann_marker_list)) {
  if(species == "human"){
    common_markers<-toupper(ann_marker_list[[i]])
  }else if(species == "mouse"){
    common_markers<-str_to_title(ann_marker_list[[i]])
  }
  gene<-unique(common_markers[common_markers %in% rownames(seurat_data@assays$RNA)])
  if (length(gene) > 0){
    ann_gene_list[[names(ann_marker_list)[i]]]<-unique(gene)
    ann_markers_all<-unique(c(ann_markers_all,gene))
  }
}
print(ann_gene_list)
print(ann_markers_all)


#注释结果可视化
Idents(object = seurat_data) <- "subcell_type2"
levels(seurat_data)

##marker堆叠小提琴图                       
p2<-VlnPlot(seurat_data, features = ann_markers_all,stack = T,combine = T,flip = T)+
  theme(axis.text.x = element_text(angle = 45,hjust = 1,size = 15))
p2
save_picture(p2, paste(out_result,"B",sep = "/") , Pname = "vlnplot_stack_marker",width_len = 1000, height_len = 1500)


###C=====================================================
Idents(object = seurat_data) <- "subcell_type2"
levels(seurat_data)

cell_type<-c("Exhausted_T","Cytotoxic_T","Treg","Activated_T")

##Exhausted_T
marker<-c("CD8A","PDCD1","LAG3")
p1<-FeaturePlot(seurat_data,features = marker,combine = FALSE)
p1
fix.sc <- scale_color_gradientn( colours = c('lightgrey', 'blue'),  limits = c(0, 5))
p2 <- lapply(p1, function (x) x + fix.sc)
# CombinePlots(p2)
output<-paste(out_result,"C","Exhausted_T",sep = "/")
if(!dir.exists(output)){
  dir.create(output,recursive = T)
}
for (i in 1:length(marker)) {
  save_picture(p2[[i]], output , Pname = marker[i],width_len = 1000, height_len = 900)
}

##Cytotoxic_T
marker<-c("CD8A","CD3E","CD3D")
p1<-FeaturePlot(seurat_data,features = marker,combine = FALSE)
p1
fix.sc <- scale_color_gradientn( colours = c('lightgrey', 'blue'),  limits = c(0, 5))
p2 <- lapply(p1, function (x) x + fix.sc)
# CombinePlots(p2)
output<-paste(out_result,"C","Cytotoxic_T",sep = "/")
if(!dir.exists(output)){
  dir.create(output,recursive = T)
}
for (i in 1:length(marker)) {
  save_picture(p2[[i]], output , Pname = marker[i],width_len = 1000, height_len = 900)
}

##Treg
marker<-c("CD4","FOXP3","IL2RA")
p1<-FeaturePlot(seurat_data,features = marker,combine = FALSE)
p1
fix.sc <- scale_color_gradientn( colours = c('lightgrey', 'blue'),  limits = c(0, 5))
p2 <- lapply(p1, function (x) x + fix.sc)
# CombinePlots(p2)
output<-paste(out_result,"C","Treg",sep = "/")
if(!dir.exists(output)){
  dir.create(output,recursive = T)
}
for (i in 1:length(marker)) {
  save_picture(p2[[i]], output , Pname = marker[i],width_len = 1000, height_len = 900)
}


##Activated_T
marker<-c("CD8A","CD69","CD3G")
p1<-FeaturePlot(seurat_data,features = marker,combine = FALSE)
p1
fix.sc <- scale_color_gradientn( colours = c('lightgrey', 'blue'),  limits = c(0, 5))
p2 <- lapply(p1, function (x) x + fix.sc)
# CombinePlots(p2)
output<-paste(out_result,"C","Activated_T",sep = "/")
if(!dir.exists(output)){
  dir.create(output,recursive = T)
}
for (i in 1:length(marker)) {
  save_picture(p2[[i]], output , Pname = marker[i],width_len = 1000, height_len = 900)
}
###D=====================================================
out<-paste(out_result,"D",sep = "/")

Idents(object = seurat_data) <- "subcell_type2"
levels(seurat_data)

#DEGs
library(future)
plan("multiprocess", workers = 10)
options(future.globals.maxSize = 10*1000 * 1024^2)
seurat_DE<-FindAllMarkers(seurat_data,only.pos = TRUE,logfc.threshold = 0)
plan("sequential")
write.csv(seurat_DE,file = paste(sep = "/",out,'all_DEGs.csv'))

seurat_DE<-read.csv(paste(sep = "/",out,'all_DEGs.csv'),row.names = 1)
seurat_DE<-subset(seurat_DE,p_val_adj < 0.05)
top5 <- seurat_DE %>% group_by(cluster) %>% top_n(5, avg_log2FC)
top5_gene<-top5$gene
top5_gene

marker_deg<-c("CCL4L2","GNLY","CCL4","CCL3","IFNG",
              "TTN","S100A2","DSP","FABP5","GSTP1",
              "TNFRSF4","LTB" ,"IL7R","BATF","IL2RA",
              "CXCL8","G0S2","FTH1","NAMPT","BCL2A1")

p<-DotPlot(seurat_data,features = marker_deg,cols = "RdYlBu")+
  theme(axis.text.x = element_text(angle = 45,hjust = 1))+
  theme(legend.position = "top",legend.key.width =  unit(30, "pt"))
p
save_picture(p, out , Pname = "DEG_marker",
             width_len = 1000, height_len = 400)


###E=============================================================================
out<-paste(out_result,"E",sep = "/")

Idents(object = seurat_data) <- "subcell_type2"
levels(seurat_data)

sub_seurat<-subset(seurat_data,subcell_type2 %in% c("Exhausted_T","Treg"))
levels(sub_seurat)

marker<-c("LAG3","PDCD1","CTLA4","TIGIT","CD96","HAVCR2","AHR")
p<-DotPlot(sub_seurat,features = marker,cols = "RdYlBu")+
  theme(axis.text.x = element_text(angle = 45,hjust = 1))+
  theme(legend.position = "top",
        legend.key.width =  unit(20, "pt"),
        legend.key.size = unit(10,"pt"))
p
save_picture(p, out , Pname = "1_marker",
             width_len = 1000, height_len = 400)


sub_seurat<-subset(seurat_data,subcell_type2 %in% c("Cytotoxic_T","Activated_T"))
levels(sub_seurat)
marker<-c("ICOS","TNFRSF9","TNFRSF14","CD27","TNFRSF4","TNFRSF18","TNFRSF25")
p<-DotPlot(sub_seurat,features = marker,cols = "RdYlBu")+
  theme(axis.text.x = element_text(angle = 45,hjust = 1))+
  theme(legend.position = "top",
        legend.key.width =  unit(20, "pt"),
        legend.key.size = unit(10,"pt"))
p
save_picture(p, out , Pname = "2_marker",
             width_len = 1000, height_len = 400)


###F==========================================================================
out<-paste(out_result,"F",sep = "/")

Idents(object = seurat_data) <- "Group"
levels(seurat_data)

p<-DimPlot(seurat_data,reduction = "umap",pt.size = 2,cols = c("green","red"))
p
save_picture(p, out , Pname = "1_Group_UMAP",
             width_len = 1000, height_len = 900)

Idents(object = seurat_data) <- "subcell_type2"
levels(seurat_data)


###细胞比例柱状图+堆叠图
###数据准备
cell_type_name<-c("subcell_type2")###选择要分析的亚群分组列

level<-NULL  ###设置亚群排列顺序，不设置则系统按字典排序
Group_data<-unique(seurat_data$Group)
if(length(Group_data)==2){
  Group_1<-Group_data[1]
  Group_2<-Group_data[2]
  Group_3<-NULL
  Group_4<-NULL
}else if(length(Group_data)==3){
  Group_1<-Group_data[1]
  Group_2<-Group_data[2]
  Group_3<-Group_data[3]
  Group_4<-NULL
}else if(length(Group_data)==4){
  Group_1<-Group_data[1]
  Group_2<-Group_data[2]
  Group_3<-Group_data[3]
  Group_4<-Group_data[4]
}
#分期绘图
par1 = Group_1
par2 = Group_2
par3 = Group_3
par4 = Group_4
ggtj<-function(x,cell_types){
  gplot<-aggregate(x,by=list(x[,cell_types]),length) ###将meta.data按照cell_type进行分组并计算meta.data每列频率数量
  a<-rownames(gplot)
  gplot<-subset(gplot,select=c('Group.1',cell_types))##提取分组及细胞亚群数量两列
  colnames(gplot)<-c("cell_type","counts")
  gplot$percent<-as.numeric(round(gplot$counts/nrow(x),4)*100)##计算百分比
  return(gplot)
}
G1<-ggtj(subset(seurat_data@meta.data,Group==par1),cell_types=cell_type_name)
G1$group<-par1
if(!(is.null(par2) | length(which(par2 == 'NULL'))>0)){
  G2<-ggtj(subset(seurat_data@meta.data,Group==par2),cell_types=cell_type_name)
  G2$group<-par2
  if (!(is.null(par3) | length(which(par3 == 'NULL'))>0)) {
    G3<-ggtj(subset(seurat_data@meta.data,Group==par3),cell_types=cell_type_name)
    G3$group<-par3
    if (!(is.null(par4)  | length(which(par4 == 'NULL'))>0)) {
      G4<-ggtj(subset(seurat_data@meta.data,Group==par4),cell_types=cell_type_name)
      G4$group<-par4
      G<-rbind(G1,G2) %>% rbind(.,G3) %>% rbind(.,G4)
    }else{
      G<-rbind(G1,G2) %>% rbind(.,G3)
    }
  }else{
    G<-rbind(G1,G2)
  }
}else{
  G<-G1
}

###设置亚群排序
if (!(is.null(level))) {
  G$cell_type<-factor(G$cell_type, levels = level)
}else{
  G$cell_type<-factor(G$cell_type,levels = levels(seurat_data@active.ident))
}

write.csv(G,file = paste0(out,"/","cellpre.csv"),quote = F,row.names = F)

###各大群样本分组细胞比例柱状图cellpre_change
P<-ggplot(G,aes(x=cell_type,y=percent,fill=group,group=group))+
  geom_bar(stat = "identity",position = "dodge",size=1.3)+
  theme_classic()+
  theme(axis.text.x = element_text(margin=margin(2,0,0,0, "mm"),size = 20,angle = 45,hjust = 1))+
  scale_y_continuous(expand = c(0.01,0))+
  # geom_text(aes(x=cell_type,y=percent,label=paste(percent,"%",sep = "")),size = 3,position = position_dodge(0.9))+
  scale_fill_manual(values = c("green","red"))
P
save_picture(P,out,Pname = "2_cellpre_change",width_len = 1000, height_len = 900)


#组间差异分析：仍然是使用这个比例数据进行分析，不过却是在各个样本中进行比例比较
out_result<-out
scedata<-seurat_data
table(scedata$orig.ident)#查看各组细胞数
table(Idents(scedata))#查看各种类型细胞数目
table(Idents(scedata), scedata$orig.ident)#各组不同细胞群细胞数
Cellratio <- prop.table(table(Idents(scedata), 
                              scedata$orig.ident), margin = 2)#计算各组样本不同细胞群比例
Cellratio <- data.frame(Cellratio)
colnames(Cellratio)<-c("cell_type","Sample","percent")
Cellratio$group<-NA
sample <- c("CC1","CC10","CC11","CC12","CC13","CC14","CC15","CC2","CC3","CC4","CC5","CC6","CC7","CC8","CC9")
group <- c("AD","AD","AD","AD","AD","SQ","SQ","AD","SQ","AD","AD","SQ","AD","AD","AD")
for (i in 1:length(sample)) {
  Cellratio$group[which(Cellratio$Sample == sample[i])]<-group[i]
}


###作图展示
library(ggplot2)
library(dplyr)
library(ggpubr)
library(cowplot)

Cellratio$group<-as.factor(Cellratio$group)
Cellratio$cell_type<-as.factor(Cellratio$cell_type)
write.csv(Cellratio,file = paste0(out_result,"/Sample_celltype_percent.csv"))

G_plot<-G
G_plot$t_test<-"t.test"
G_plot$p_value<-NA
p_result<-c()
for (i in levels(seurat_data)) {
  Cellratio_plot<-subset(Cellratio,cell_type == i)
  group1<-Cellratio_plot$percent[which(Cellratio_plot$group == "AD")]
  group2<-Cellratio_plot$percent[which(Cellratio_plot$group == "SQ")]
  diff_result<-t.test(group1,group2)
  p<-diff_result$p.value
  G_plot$p_value[which(G_plot$cell_type == i)]<-paste0("p.value: ",round(p,4))
  p_result<-c(p_result,p)
}

G_plot$wilcox<-"wilcox.test"
G_plot$p_value_wilcox<-NA
p_result<-c()
for (i in levels(seurat_data)) {
  Cellratio_plot<-subset(Cellratio,cell_type == i)
  group1<-Cellratio_plot$percent[which(Cellratio_plot$group == "AD")]
  group2<-Cellratio_plot$percent[which(Cellratio_plot$group == "SQ")]
  diff_result<-wilcox.test(group1,group2)
  p<-diff_result$p.value
  G_plot$p_value_wilcox[which(G_plot$cell_type == i)]<-paste0("p.value: ",round(p,4))
  p_result<-c(p_result,p)
}


###柱状图
pp1 = ggplot(G_plot,aes(x=group,y=percent,fill = group))+
  geom_bar(stat = "identity", position = "dodge", width = 1) +
  ylab("percent") +
  # geom_errorbar(aes(ymin = len - 1.96 * sd, ymax = len + 1.96 * sd), width = 0.2, position = position_dodge(0.7)) +
  facet_grid(~cell_type+t_test+p_value) +
  theme_cowplot() +
  scale_y_continuous(expand = c(0,0))+
  # geom_signif(test="wilcox.test", comparisons = list(c("AD","SQ")), map_signif_level = T)
  scale_fill_manual(values = c("green","red")) +
  theme(axis.text = element_text(size = 10),axis.title = element_text(size = 10),legend.text = element_text(size = 10),
        legend.title = element_text(size = 10),plot.title = element_text(size = 10,face = 'plain'),legend.position = 'none')

pp1
save_picture(pp1, out_result , Pname = "cellbar_t_test",width_len = 550, height_len = 1000)

pp2 = ggplot(G_plot,aes(x=group,y=percent,fill = group))+
  geom_bar(stat = "identity", position = "dodge", width = 1) +
  ylab("percent") +
  # geom_errorbar(aes(ymin = len - 1.96 * sd, ymax = len + 1.96 * sd), width = 0.2, position = position_dodge(0.7)) +
  facet_grid(~cell_type+wilcox+p_value_wilcox) +
  theme_cowplot() +
  scale_y_continuous(expand = c(0,0))+
  # geom_signif(test="wilcox.test", comparisons = list(c("AD","SQ")), map_signif_level = T)
  scale_fill_manual(values = c("green","red")) +
  theme(axis.text = element_text(size = 10),axis.title = element_text(size = 10),legend.text = element_text(size = 10),
        legend.title = element_text(size = 10),plot.title = element_text(size = 10,face = 'plain'),legend.position = 'none')

pp2
save_picture(pp2, out_result , Pname = "cellbar_wilcox",width_len = 550, height_len = 1000)


##HPV_Group
out_result<-paste(out,"other_barplot","HPV",sep = "/")
if(!dir.exists(out_result)){
  dir.create(out_result,recursive = T)
}
#按HPV+和HPV-分群图====
library(plyr)
seurat_data@meta.data$Group_HPV <- mapvalues(seurat_data@meta.data$orig.ident,
                                            from = c('CC1','CC2','CC4','CC5',
                                                     'CC7','CC8','CC9','CC10','CC11','CC12',
                                                     'CC13'), 
                                            to=c('P','P','N','P','P','P','N','N','N','N','P'))
seurat_data@meta.data$Group_HPV <- ifelse(seurat_data$Group_HPV != 'N' & seurat_data$Group_HPV != 'P','SQ',seurat_data$Group_HPV)
DimPlot(seurat_data,group.by = 'Group_HPV',label = TRUE,label.size = 10,pt.size = 0.8,raster=FALSE,cols = c(Science_col[1:2],'gray',''))+ggtitle('sample')
DimPlot(seurat_data,reduction = "tsne",group.by = 'Group_HPV',label = TRUE,label.size = 10,pt.size = 0.8,raster=FALSE,cols = c(Science_col[1:2],'gray',''))+ggtitle('sample')

# #按病人FIGO分期的分群图====
# library(openxlsx)
# add_sample_data<-read.xlsx('E:/work_station/2023/3_6_HPV2/样本信息（简化）.xlsx')
# add_sample_data$sample<-gsub('-','',add_sample_data$sample)
# library(plyr)
# seurat_data@meta.data$FIGO_group <- mapvalues(seurat_data@meta.data$orig.ident,
#                                              from = add_sample_data$sample, 
#                                              to=add_sample_data$FIGO_group)
# 
# DimPlot(seurat_data,group.by = 'FIGO_group',label = TRUE,label.size = 10,pt.size = 0.8,raster=FALSE,cols = c(Science_col[1:2],'gray',''))+ggtitle('sample')
# DimPlot(seurat_data,reduction = "tsne",group.by = 'FIGO_group',label = TRUE,label.size = 10,pt.size = 0.8,raster=FALSE,cols = c(Science_col[1:2],'gray',''))+ggtitle('sample')
# 
# 


seurat_ann2<-subset(seurat_data,Group_HPV %in% c("P",'N'))
p<-DimPlot(seurat_ann2,group.by = 'Group_HPV',
           label = TRUE,label.size = 10,
           pt.size = 0.8,raster=FALSE,
           cols = Science_col)+
  ggtitle('sample')
p
save_picture(p, out_result , Pname = "1_Group_UMAP",
             width_len = 1000, height_len = 900)


p<-DimPlot(seurat_ann2,reduction = "tsne",group.by = 'Group_HPV',
           label = TRUE,label.size = 10,
           pt.size = 0.8,raster=FALSE,
           cols = Science_col)+
  ggtitle('sample')
p
save_picture(p, out_result , Pname = "1_Group_TSNE",
             width_len = 1000, height_len = 900)


p<-DimPlot(seurat_ann2,group.by = 'Group_HPV',
           label = FALSE,label.size = 10,
           pt.size = 0.8,raster=FALSE,
           cols = Science_col)+
  ggtitle('sample')
p
save_picture(p, out_result , Pname = "2_Group_UMAP",
             width_len = 1000, height_len = 900)


p<-DimPlot(seurat_ann2,reduction = "tsne",group.by = 'Group_HPV',
           label = FALSE,label.size = 10,
           pt.size = 0.8,raster=FALSE,
           cols = Science_col)+
  ggtitle('sample')
p
save_picture(p, out_result , Pname = "2_Group_TSNE",
             width_len = 1000, height_len = 900)

cell_type_name<-c("subcell_type2")###选择要分析的亚群分组列

Group_1<-'P'
Group_2<-'N'
Group_3<-NULL
Group_4<-NULL
level<-NULL  ###设置亚群排列顺序，不设置则系统按字典排序
#分期绘图
par1 = Group_1
par2 = Group_2
par3 = Group_3
par4 = Group_4
ggtj<-function(x,cell_types){
  gplot<-aggregate(x,by=list(x[,cell_types]),length) ###将meta.data按照cell_type进行分组并计算meta.data每列频率数量
  a<-rownames(gplot)
  gplot<-subset(gplot,select=c('Group.1',cell_types))##提取分组及细胞亚群数量两列
  colnames(gplot)<-c("cell_type","counts")
  gplot$percent<-as.numeric(round(gplot$counts/nrow(x),4)*100)##计算百分比
  return(gplot)
}
G1<-ggtj(subset(seurat_ann2@meta.data,Group_HPV ==par1),cell_types=cell_type_name)
G1$group<-par1
if(!(is.null(par2) | length(which(par2 == 'NULL'))>0)){
  G2<-ggtj(subset(seurat_ann2@meta.data,Group_HPV==par2),cell_types=cell_type_name)
  G2$group<-par2
  if (!(is.null(par3) | length(which(par3 == 'NULL'))>0)) {
    G3<-ggtj(subset(seurat_ann2@meta.data,Group_HPV==par3),cell_types=cell_type_name)
    G3$group<-par3
    if (!(is.null(par4)  | length(which(par4 == 'NULL'))>0)) {
      G4<-ggtj(subset(seurat_ann2@meta.data,Group_HPV==par4),cell_types=cell_type_name)
      G4$group<-par4
      G<-rbind(G1,G2) %>% rbind(.,G3) %>% rbind(.,G4)
    }else{
      G<-rbind(G1,G2) %>% rbind(.,G3)
    }
  }else{
    G<-rbind(G1,G2)
  }
}else{
  G<-G1
}

###设置亚群排序
if (!(is.null(level))) {
  G$cell_type<-factor(G$cell_type, levels = level)
}else{
  G$cell_type<-factor(G$cell_type,levels = levels(seurat_data@active.ident))
}

type_num<-length(levels(seurat_ann2))
for (i in unique(G$group)) {
  if(length(which(G$group == i)) == type_num){
    G<-G
  }else{
    cell_type_no<-levels(seurat_ann2)[!(levels(seurat_ann2) %in% G$cell_type[which(G$group == i)])]
    temp<-matrix(NA,nrow = length(cell_type_no),ncol = 4)
    temp[,1]<-cell_type_no
    temp[,2]<-0
    temp[,3]<-0
    temp[,4]<-i
    temp<-as.data.frame(temp)
    colnames(temp)<-colnames(G)
    G<-rbind(G,temp)
  }
}

G$counts<-as.numeric(G$counts)
G$percent<-as.numeric(G$percent)
write.csv(G,file = paste0(out_result,"/","cellpre.csv"),quote = F,row.names = F)

###各大群样本分组细胞比例柱状图cellpre_change
library(ggplot2)
library(ggpubr)
P<-ggplot(G,aes(x=cell_type,y=percent,fill=group,group=group))+
  geom_bar(stat = "identity",position = "dodge",size=1.3)+
  theme_classic()+
  theme(axis.text.x = element_text(margin=margin(2,0,0,0, "mm"),size = 15,angle = 90,hjust = 1))+
  scale_y_continuous(expand = c(0.01,0))+
  # geom_text(aes(x=cell_type,y=percent,label=paste(percent,"%",sep = "")),size = 3,position = position_dodge(0.9))+
  scale_fill_manual(values = Science_col)
P
save_picture(P,out_result,Pname = "cellpre_change",width_len = 500, height_len = 800)

#组间差异分析：仍然是使用这个比例数据进行分析，不过却是在各个样本中进行比例比较
scedata<-seurat_ann2
table(scedata$orig.ident)#查看各组细胞数
table(Idents(scedata))#查看各种类型细胞数目
table(Idents(scedata), scedata$orig.ident)#各组不同细胞群细胞数
Cellratio <- prop.table(table(Idents(scedata), 
                              scedata$orig.ident), margin = 2)#计算各组样本不同细胞群比例
Cellratio <- data.frame(Cellratio)
colnames(Cellratio)<-c("cell_type","Sample","percent")
Cellratio$group<-NA
sample <- c('CC1','CC2','CC4','CC5',
            'CC7','CC8','CC9','CC10','CC11','CC12',
            'CC13')
group <- c('P','P','N','P','P','P','N','N','N','N','P')
for (i in 1:length(sample)) {
  Cellratio$group[which(Cellratio$Sample == sample[i])]<-group[i]
}


###作图展示
library(ggplot2)
library(dplyr)
library(ggpubr)
library(cowplot)

Cellratio$group<-as.factor(Cellratio$group)
Cellratio$cell_type<-as.factor(Cellratio$cell_type)
write.csv(Cellratio,file = paste0(out_result,"/Sample_celltype_percent.csv"))


G_plot<-G
G_plot$t_test<-"t.test"
G_plot$p_value<-NA
p_result<-c()
for (i in levels(seurat_data)) {
  Cellratio_plot<-subset(Cellratio,cell_type == i)
  group1<-Cellratio_plot$percent[which(Cellratio_plot$group == "P")]
  group2<-Cellratio_plot$percent[which(Cellratio_plot$group == "N")]
  if(length(group1) == 0|length(group2) == 0){
    p<-NA
    G_plot$p_value[which(G_plot$cell_type == i)]<-"p.value: NA"
    p_result<-c(p_result,p)
  }else{
    diff_result<-t.test(group1,group2)
    p<-diff_result$p.value
    G_plot$p_value[which(G_plot$cell_type == i)]<-paste0("p.value: ",round(p,4))
    p_result<-c(p_result,p)
  }
}

G_plot$wilcox<-"wilcox.test"
G_plot$p_value_wilcox<-NA
p_result<-c()
for (i in levels(seurat_data)) {
  Cellratio_plot<-subset(Cellratio,cell_type == i)
  group1<-Cellratio_plot$percent[which(Cellratio_plot$group == "P")]
  group2<-Cellratio_plot$percent[which(Cellratio_plot$group == "N")]
  if(length(group1) == 0|length(group2) == 0){
    p<-NA
    G_plot$p_value_wilcox[which(G_plot$cell_type == i)]<-"p.value: NA"
    p_result<-c(p_result,p)
  }else{
    diff_result<-wilcox.test(group1,group2)
    p<-diff_result$p.value
    G_plot$p_value_wilcox[which(G_plot$cell_type == i)]<-paste0("p.value: ",round(p,4))
    p_result<-c(p_result,p)
  }
}


###柱状图
pp1 = ggplot(G_plot,aes(x=group,y=percent,fill = group))+
  geom_bar(stat = "identity", position = "dodge", width = 1) +
  ylab("percent") +
  # geom_errorbar(aes(ymin = len - 1.96 * sd, ymax = len + 1.96 * sd), width = 0.2, position = position_dodge(0.7)) +
  facet_grid(~cell_type+t_test+p_value) +
  theme_cowplot() +
  scale_y_continuous(expand = c(0,0))+
  # geom_signif(test="wilcox.test", comparisons = list(c("AD","SQ")), map_signif_level = T)
  scale_fill_manual(values = Science_col) +
  theme(axis.text = element_text(size = 10),axis.title = element_text(size = 10),legend.text = element_text(size = 10),
        legend.title = element_text(size = 10),plot.title = element_text(size = 10,face = 'plain'),legend.position = 'none')

pp1
save_picture(pp1, out_result , Pname = "cellbar_t_test",width_len = 550, height_len = 1000)

pp2 = ggplot(G_plot,aes(x=group,y=percent,fill = group))+
  geom_bar(stat = "identity", position = "dodge", width = 1) +
  ylab("percent") +
  # geom_errorbar(aes(ymin = len - 1.96 * sd, ymax = len + 1.96 * sd), width = 0.2, position = position_dodge(0.7)) +
  facet_grid(~cell_type+wilcox+p_value_wilcox) +
  theme_cowplot() +
  scale_y_continuous(expand = c(0,0))+
  # geom_signif(test="wilcox.test", comparisons = list(c("AD","SQ")), map_signif_level = T)
  scale_fill_manual(values = Science_col) +
  theme(axis.text = element_text(size = 10),axis.title = element_text(size = 10),legend.text = element_text(size = 10),
        legend.title = element_text(size = 10),plot.title = element_text(size = 10,face = 'plain'),legend.position = 'none')

pp2
save_picture(pp2, out_result , Pname = "cellbar_wilcox",width_len = 550, height_len = 1000)


#Group NS3/S3 =====
out_result<-paste(out,"other_barplot","Stage",sep = "/")
if(!dir.exists(out_result)){
  dir.create(out_result,recursive = T)
}

cell_type_name<-c("subcell_type2")###选择要分析的亚群分组列


#按病人FIGO分期的分群图=
library(openxlsx)
add_sample_data<-read.xlsx('E:/work_station/2023/3_6_HPV2/样本信息（简化）.xlsx')
add_sample_data$sample<-gsub('-','',add_sample_data$sample)
library(plyr)
seurat_data@meta.data$FIGO_group <- mapvalues(seurat_data@meta.data$orig.ident,
                                             from = add_sample_data$sample,
                                             to=add_sample_data$FIGO_group)

p<-DimPlot(seurat_data,group.by = 'FIGO_group',
           label = TRUE,label.size = 10,
           pt.size = 0.8,raster=FALSE,
           cols = c("#7830cc","#f2ca27"))+
  ggtitle('sample')
p
save_picture(p, out_result , Pname = "1_Group_UMAP",
             width_len = 1000, height_len = 900)

p<-DimPlot(seurat_data,reduction = "tsne",group.by = 'FIGO_group',
           label = TRUE,label.size = 10,
           pt.size = 0.8,raster=FALSE,
           cols = c("#7830cc","#f2ca27"))+ggtitle('sample')
p
save_picture(p, out_result , Pname = "1_Group_TSNE",
             width_len = 1000, height_len = 900)


p<-DimPlot(seurat_data,group.by = 'FIGO_group',
           label = FALSE,label.size = 10,
           pt.size = 0.8,raster=FALSE,
           cols = c("#7830cc","#f2ca27"))+
  ggtitle('sample')
p
save_picture(p, out_result , Pname = "2_Group_UMAP",
             width_len = 1000, height_len = 900)

p<-DimPlot(seurat_data,reduction = "tsne",group.by = 'FIGO_group',
           label = FALSE,label.size = 10,
           pt.size = 0.8,raster=FALSE,
           cols = c("#7830cc","#f2ca27"))+ggtitle('sample')
p
save_picture(p, out_result , Pname = "2_Group_TSNE",
             width_len = 1000, height_len = 900)

# table(seurat_data$FIGO_group)#查看各组细胞数
# table(Idents(seurat_data))#查看各种类型细胞数目
# table(Idents(seurat_data), seurat_data$FIGO_group)#各组不同细胞群细胞数
# Cellratio <- prop.table(table(Idents(seurat_data), 
#                               seurat_data$FIGO_group), margin = 2)#计算各组样本不同细胞群比例
# Cellratio <- data.frame(Cellratio)
# colnames(Cellratio)<-c("cell_type","group","percent")
# 
# G<-Cellratio

Group_1<-'S3'
Group_2<-'NS3'
Group_3<-NULL
Group_4<-NULL
level<-NULL  ###设置亚群排列顺序，不设置则系统按字典排序
#分期绘图
par1 = Group_1
par2 = Group_2
par3 = Group_3
par4 = Group_4
ggtj<-function(x,cell_types){
  gplot<-aggregate(x,by=list(x[,cell_types]),length) ###将meta.data按照cell_type进行分组并计算meta.data每列频率数量
  a<-rownames(gplot)
  gplot<-subset(gplot,select=c('Group.1',cell_types))##提取分组及细胞亚群数量两列
  colnames(gplot)<-c("cell_type","counts")
  gplot$percent<-as.numeric(round(gplot$counts/nrow(x),4)*100)##计算百分比
  return(gplot)
}
G1<-ggtj(subset(seurat_data@meta.data,FIGO_group ==par1),cell_types=cell_type_name)
G1$group<-par1
if(!(is.null(par2) | length(which(par2 == 'NULL'))>0)){
  G2<-ggtj(subset(seurat_data@meta.data,FIGO_group==par2),cell_types=cell_type_name)
  G2$group<-par2
  if (!(is.null(par3) | length(which(par3 == 'NULL'))>0)) {
    G3<-ggtj(subset(seurat_data@meta.data,FIGO_group==par3),cell_types=cell_type_name)
    G3$group<-par3
    if (!(is.null(par4)  | length(which(par4 == 'NULL'))>0)) {
      G4<-ggtj(subset(seurat_data@meta.data,FIGO_group==par4),cell_types=cell_type_name)
      G4$group<-par4
      G<-rbind(G1,G2) %>% rbind(.,G3) %>% rbind(.,G4)
    }else{
      G<-rbind(G1,G2) %>% rbind(.,G3)
    }
  }else{
    G<-rbind(G1,G2)
  }
}else{
  G<-G1
}

###设置亚群排序
if (!(is.null(level))) {
  G$cell_type<-factor(G$cell_type, levels = level)
}else{
  G$cell_type<-factor(G$cell_type,levels = levels(seurat_data@active.ident))
}

write.csv(G,file = paste0(out_result,"/","cellpre.csv"),quote = F,row.names = F)

###各大群样本分组细胞比例柱状图cellpre_change
library(ggplot2)
library(ggpubr)
P<-ggplot(G,aes(x=cell_type,y=percent,fill=group,group=group))+
  geom_bar(stat = "identity",position = "dodge",size=1.3)+
  theme_classic()+
  theme(axis.text.x = element_text(margin=margin(2,0,0,0, "mm"),size = 15,angle = 90,hjust = 1))+
  scale_y_continuous(expand = c(0.01,0))+
  # geom_text(aes(x=cell_type,y=percent,label=paste(percent,"%",sep = "")),size = 3,position = position_dodge(0.9))+
  scale_fill_manual(values = c("#7830cc","#f2ca27"))
P
save_picture(P,out_result,Pname = "cellpre_change",width_len = 550, height_len = 900)


#组间差异分析：仍然是使用这个比例数据进行分析，不过却是在各个样本中进行比例比较
scedata<-seurat_data
table(scedata$orig.ident)#查看各组细胞数
table(Idents(scedata))#查看各种类型细胞数目
table(Idents(scedata), scedata$orig.ident)#各组不同细胞群细胞数
Cellratio <- prop.table(table(Idents(scedata), 
                              scedata$orig.ident), margin = 2)#计算各组样本不同细胞群比例
Cellratio <- data.frame(Cellratio)
colnames(Cellratio)<-c("cell_type","Sample","percent")
Cellratio$group<-NA
sample <- add_sample_data$sample
group <- add_sample_data$FIGO_group
for (i in 1:length(sample)) {
  Cellratio$group[which(Cellratio$Sample == sample[i])]<-group[i]
}


###作图展示
library(ggplot2)
library(dplyr)
library(ggpubr)
library(cowplot)

Cellratio$group<-as.factor(Cellratio$group)
Cellratio$cell_type<-as.factor(Cellratio$cell_type)
write.csv(Cellratio,file = paste0(out_result,"/Sample_celltype_percent.csv"))


G_plot<-G
G_plot$t_test<-"t.test"
G_plot$p_value<-NA
p_result<-c()
for (i in levels(seurat_data)) {
  Cellratio_plot<-subset(Cellratio,cell_type == i)
  group1<-Cellratio_plot$percent[which(Cellratio_plot$group == "S3")]
  group2<-Cellratio_plot$percent[which(Cellratio_plot$group == "NS3")]
  if(length(group1) == 0|length(group2) == 0){
    p<-NA
    G_plot$p_value[which(G_plot$cell_type == i)]<-"p.value: NA"
    p_result<-c(p_result,p)
  }else{
    diff_result<-t.test(group1,group2)
    p<-diff_result$p.value
    G_plot$p_value[which(G_plot$cell_type == i)]<-paste0("p.value: ",round(p,4))
    p_result<-c(p_result,p)
  }
}

G_plot$wilcox<-"wilcox.test"
G_plot$p_value_wilcox<-NA
p_result<-c()
for (i in levels(seurat_data)) {
  Cellratio_plot<-subset(Cellratio,cell_type == i)
  group1<-Cellratio_plot$percent[which(Cellratio_plot$group == "S3")]
  group2<-Cellratio_plot$percent[which(Cellratio_plot$group == "NS3")]
  if(length(group1) == 0|length(group2) == 0){
    p<-NA
    G_plot$p_value_wilcox[which(G_plot$cell_type == i)]<-"p.value: NA"
    p_result<-c(p_result,p)
  }else{
    diff_result<-wilcox.test(group1,group2)
    p<-diff_result$p.value
    G_plot$p_value_wilcox[which(G_plot$cell_type == i)]<-paste0("p.value: ",round(p,4))
    p_result<-c(p_result,p)
  }
}


###柱状图
pp1 = ggplot(G_plot,aes(x=group,y=percent,fill = group))+
  geom_bar(stat = "identity", position = "dodge", width = 1) +
  ylab("percent") +
  # geom_errorbar(aes(ymin = len - 1.96 * sd, ymax = len + 1.96 * sd), width = 0.2, position = position_dodge(0.7)) +
  facet_grid(~cell_type+t_test+p_value) +
  theme_cowplot() +
  scale_y_continuous(expand = c(0,0))+
  # geom_signif(test="wilcox.test", comparisons = list(c("AD","SQ")), map_signif_level = T)
  scale_fill_manual(values = c("#7830cc","#f2ca27")) +
  theme(axis.text = element_text(size = 10),axis.title = element_text(size = 10),legend.text = element_text(size = 10),
        legend.title = element_text(size = 10),plot.title = element_text(size = 10,face = 'plain'),legend.position = 'none')

pp1
save_picture(pp1, out_result , Pname = "cellbar_t_test",width_len = 550, height_len = 1000)

pp2 = ggplot(G_plot,aes(x=group,y=percent,fill = group))+
  geom_bar(stat = "identity", position = "dodge", width = 1) +
  ylab("percent") +
  # geom_errorbar(aes(ymin = len - 1.96 * sd, ymax = len + 1.96 * sd), width = 0.2, position = position_dodge(0.7)) +
  facet_grid(~cell_type+wilcox+p_value_wilcox) +
  theme_cowplot() +
  scale_y_continuous(expand = c(0,0))+
  # geom_signif(test="wilcox.test", comparisons = list(c("AD","SQ")), map_signif_level = T)
  scale_fill_manual(values = c("#7830cc","#f2ca27")) +
  theme(axis.text = element_text(size = 10),axis.title = element_text(size = 10),legend.text = element_text(size = 10),
        legend.title = element_text(size = 10),plot.title = element_text(size = 10,face = 'plain'),legend.position = 'none')

pp2
save_picture(pp2, out_result , Pname = "cellbar_wilcox",width_len = 550, height_len = 1000)
```
