## Supplementary Data 1-annotated copies of script for "Single-cell profiling reveals the intratumor heterogeneity and immunosuppressive microenvironment in cervical adenocarcinoma": Neu_cluster_ann.R

```
###Neu细胞分群注释

#大群注释====
seurat_deal<-readRDS("E:/work_station/2023/3_6_HPV2/01_data/HPV2_seurat_deal.rds")
out<-"E:/work_station/2023/3_6_HPV2/03_ann"
Group_1<-'AD'
Group_2<-'SQ'
Group_3<-NULL
Group_4<-NULL

seurat_ann<-ann_and_plot(seurat_deal, out,
                         ann_file = 'E:/work_station/2023/3_6_HPV2/02_all/HPV2_ann.txt',
                         meta_names = "cell_type", label_size = 10, pt.size = 1)

#Neu分群
#human_after_CCA:亚群标准分析
human_after_CCA<-function(x,resolution,t=1,mem=1){
  suppressMessages(library("tidyverse"))
  suppressMessages(library('ggplot2'))
  suppressMessages(library("scales"))
  suppressMessages(library("gridExtra"))
  suppressMessages(library("Matrix"))
  suppressMessages(library("Seurat"))
  suppressMessages(library('harmony'))
  suppressMessages(library('cowplot'))
  suppressMessages(library('future'))
  if (!(exists("resolution"))) {
    stop("Error: resolution is null")
  }
  t_after_CCA<-Sys.time()
  set.seed(2022)
  message("**Statistical Period information")
  s.genes <- read.table(file = "~/细胞周期标志物S期.txt",sep="\t",header=T)
  g2m.genes <- read.table(file = "~/细胞周期标志物G2M期.txt",sep="\t",header=T)
  message('物种：human')
  g2m.genes<-g2m.genes$G2M_gene
  s.genes<-s.genes$S_gene
  x<- CellCycleScoring(object = x, g2m.features = g2m.genes, s.features = s.genes)
  message('**Normalization is under way...')
  x <- NormalizeData(x, normalization.method = "LogNormalize", scale.factor = 10000)
  x <- FindVariableFeatures(x, selection.method = "vst", nfeatures = 2000)
  message('**It is being standardized...')
  library(future)
  if (t==1) {
    plan("sequential")
  }else{
    plan("multiprocess", workers = t)
    options(future.globals.maxSize = as.numeric(mem)*1000 * 1024^2)
  }
  x <- ScaleData(x, features = rownames(x),vars.to.regress = c("percent.mt","S.Score","G2M.Score"))
  message('**Pca dimension reduction is in progress')
  x <- RunPCA(x, features = VariableFeatures(object = x),verbose = FALSE)
  message('**Harmony correction in progress...')
  x <- RunHarmony(x,"orig.ident", plot_convergence = TRUE)
  message('**Clustering and visualization in progress...')
  dim.use<-30
  #####降维+聚类+分群
  x <- x %>% 
    RunUMAP(reduction = "harmony", dims = 1:dim.use) %>% 
    FindNeighbors(reduction = "harmony", dims = 1:dim.use) %>% 
    FindClusters(resolution = resolution)
  x <- x %>% 
    RunTSNE(reduction = "harmony", dims = 1:dim.use)
  
  plan("sequential")
  print(Sys.time()-t_after_CCA)
  return(x)
}
temp_subseurat<-subset(seurat_ann, idents = "Neutrophils")
temp_subseurat_deal<-human_after_CCA(temp_subseurat, resolution = 0.1, t = 4, mem = 30)


###Neu注释：先进行差异分析，将分群的差异基因按照对应的avg_log2FC升序排序，取排序第一的差异基因名作为该群的名字
temp_DE<-FindAllMarkers(temp_subseurat_deal,logfc.threshold = 0,only.pos = T) ###差异分析

new_id<-1:length(levels(temp_subseurat_deal))
names(new_id)<-levels(temp_subseurat_deal)
for (i in 1:length(levels(temp_subseurat_deal))) {
  new_id[i]<-paste(subset(temp_DE, cluster==names(new_id)[i]) %>% arrange(-avg_log2FC) %>% .[1,"gene"],
                   names(subgroup)[[groupone]],sep = "+_")
}

new_id
temp_subseurat_deal <- RenameIdents(temp_subseurat_deal,new_id)
meta_names = "sub_celltype"
temp_subseurat_deal@meta.data[,meta_names]<-as.factor(temp_subseurat_deal@active.ident)
saveRDS(temp_subseurat_deal,file = "E:/work_station/2023/3_6_HPV2/05_subset/RDS/Neutrophils_ann.rds")

###Neu重新注释:根据“Neutrophils_ann.txt”注释分簇 ====
ann_and_plot<-function(mouse_deal, out, ann_file, meta_names,label_size, pt.size){
  if (!(dir.exists(paste(out,"RDS",sep = "/")))) {
    message("warning: don't exists RDS file, but already creat")
    dir.create(paste(out,"RDS",sep = "/"))
  }
  if (!(file.exists(ann_file))) {
    stop("Error: ann file isn't existsed")
  }
  #读取注释文件
  ann<-read.table(file = ann_file,
                  header = F,check.names = F)
  ##注释
  all.equal(as.character(levels(mouse_deal)),as.character(ann$V1))
  new_id<-ann$V2
  names(new_id)<-levels(mouse_deal)
  new_id
  mouse_deal <- RenameIdents(mouse_deal,new_id)
  all.equal(rownames(mouse_deal@meta.data),names(mouse_deal@active.ident))
  mouse_deal@meta.data[,meta_names]<-as.factor(mouse_deal@active.ident)
  #saveRDS(mouse_deal,file = "D:/work_station_space/2022/7_1_liver_luyao/7_1_liver_luyao/04_ann_before/RDS/mouse_ann.rds")
  
  ##注释后二维图
  P<-list()
  P[[1]]<-DimPlot(mouse_deal,label = TRUE,label.size = label_size,pt.size = pt.size,raster=FALSE,
                  cols = Science_col)
  P[[2]]<-DimPlot(mouse_deal,reduction = "tsne",label = TRUE,label.size = label_size,pt.size = pt.size,raster=FALSE,
                  cols = Science_col)
  names(P)<-c("umap","tsne")
  
  for (i in 1:length(P)) {
    while (!is.null(dev.list()))  dev.off()
    png(filename = paste(out,paste0("注释后",names(P)[i],"图.png"),sep = "/"),width = 1100,height = 900)
    print(P[[i]])
    while (!is.null(dev.list()))  dev.off()
    while (!is.null(dev.list()))  dev.off()
    pdf(file = paste(out,paste0("注释后",names(P)[i],"图.pdf"),sep = "/"),width = 11,height = 9)
    print(P[[i]])
    while (!is.null(dev.list()))  dev.off()
  }
  
  return(mouse_deal)
}

seurat_ann<-readRDS('E:/work_station/2023/3_6_HPV2/05_subset/RDS/Neutrophils_ann.rds')
seurat_ann<-ann_and_plot(seurat_ann,
                         'E:/work_station/2023/3_6_HPV2/05_subset/Neutrophils',
                         'E:/work_station/2023/3_6_HPV2/05_subset/Neutrophils_ann.txt',
                         meta_names = 'subcell_type',
                         label_size = 10,
                         pt.size = 1.5)
```
